# Leveraging heterogeneity for neural computation with fading memory in layer 2/3 cortical microcircuits

**DOI:** 10.1101/230821

**Authors:** Renato Duarte, Abigail Morrison

## Abstract

Complexity and heterogeneity are intrinsic to neurobiological systems, manifest in every process, at every scale, and are inextricably linked to the systems’ emergent collective behaviours and function. However, the majority of studies addressing the dynamics and computational properties of biologically inspired cortical microcircuits tend to assume (often for the sake of analytical tractability) a great degree of homogeneity in both neuronal and synaptic/connectivity parameters. While simplification and reductionism are necessary to understand the brain’s functional principles, disregarding the existence of the multiple heterogeneities in the cortical composition, which may be at the core of its computational proficiency, will inevitably fail to account for important phenomena and limit the scope and generalizability of cortical models. We address these issues by studying the individual and composite functional roles of heterogeneities in neuronal, synaptic and structural properties in a biophysically plausible layer 2/3 microcircuit model, built and constrained by multiple sources of empirical data. This approach was made possible by the emergence of large-scale, well curated databases, as well as the substantial improvements in experimental methodologies achieved over the last few years. Our results show that variability in single neuron parameters is the dominant source of functional specialization, leading to highly proficient microcircuits with much higher computational power than their homogeneous counterparts. We further show that fully heterogeneous circuits, which are closest to the biophysical reality, owe their response properties to the differential contribution of different sources of heterogeneity.

## 1 Introduction

Variability is the engine of evolution, endowing organisms with complex, heterogeneous features upon which natural selection can act (Darwin, 1859). As a consequence, heterogeneity and diversity are ubiquitous design principles in neurobiology (or in any biological system, for that matter), covering components and mechanisms at every descriptive scale (Koch, 1999). While many of these specializations, as well as their inherent complexity and diversity, are functionally meaningful, intrinsically linked and responsible for the brain’s computational capacity and efficiency (see e.g. Duarte et al. 2017a; Gjorgjieva et al. 2016; Singer 2015), others are bound to reflect epiphenomena, by-products of evolution, bearing little or no functional significance (Otopalik et al., 2017), or to subserve metabolic/maintenance tasks (Bélanger et al., 2011) that, while crucial for healthy function, are not directly involved in the computational process.

To address these issues and study the functional role of heterogeneity in cortical processing, we need to modularize complexity (Mappes and Lindstrom, 2012): exploit the deeply degenerate nature of the system (Edelman and Gally, 2001; Price and Friston, 2002) and heuristically identify groups of components that may behave as singular modules (depending on the scale and processes of interest). Once these tentative ‘building blocks’ are identified, we need to specify adequate levels of descriptive complexity that may shed light onto the underlying functional principles. These pursuits, however, pose severe epistemological problems as we currently have no clear intuition as to what ‘adequacy’ means in this context (see, e.g. (Hopfield, 1994; Krakauer et al., 2017; Marom, 2009)).

Despite substantial progress, our ability to clearly identify the system’s core component ‘building blocks’ (Getting, 1989; Gjorgjieva et al., 2016) and to systematically characterize their relative contributions and potential functional roles is still a daunting task given the multiple spatial and temporal scales at which they operate, their complex, nested interactions and the, often incomplete or inconsistent, empirical evidence. Nevertheless, when studying neuronal computation, one needs to keep in mind that, despite its tremendous complexity or because of it, the brain is a machine ‘fit-for-purpose’ and optimized to process information and operate in complex, dynamic and uncertain environments, whose spatiotemporal structure it must extract in order to reliably compute (Koch, 1999; Singer, 2015). As such, it is important to disentangle and quantify which and how the different components of the system may modulate functional neurodynamics in meaningful ways that can be paralleled with related experimental observations, and to which degree these specializations affect the system’s operations.

Combined with the use of large and complex datasets, computational modelling and numerical simulation are a central tool to make progress in this endeavour and attempt to discern adequate levels of descriptive complexity (Carandini, 2012; Gerstner et al., 2012; Piccinini and Shagrir, 2014). Doing so requires, on the one hand, the integration of data relating to processes at multiple scales (Churchland and Abbott, 2016; Gao and Ganguli, 2015; Markowetz, 2017) into mechanistic or phenomenological models of neuronal, synaptic and population dynamics and, on the other hand, the establishment of appropriate comparisons and benchmarks in order to systematically evaluate the functional consequences of complexity and heterogeneity in the different components. In the next section, we attempt to identify an appropriate partition of potential building blocks of complexity and heterogeneity in neocortical microcircuits. We then proceed to building a complex, biologically inspired and strongly data-driven microcircuit model that explicitly employs this partition in an attempt to understand and quantify the differential functional roles and consequences of these different sources of heterogeneity applying systematic and generic benchmarks. This approach also aims at providing an intermediate level of descriptive complexity for studying computation in cortical microcircuit models, as discussed below.

### 1.1 Heterogeneous building blocks in the neocortical circuitry

On a macroscopic level, hierarchical modularity is easily identifiable as a parsimonious design principle underlying various structural and functional aspects of cortical organization (Friston, 2005; Meunier et al., 2010; Mountcastle, 1978, 1997; Park and Friston, 2013). Different anatomophysiological (Shinomoto et al., 2003; VanEssen, 2013), genetic / biochemical (Hawrylycz et al., 2012; Kang et al., 2011; O’Rourke et al., 2012; Pletikos et al., 2014; Zilles et al., 2004) or functional (Mueller et al., 2013; Power et al., 2011; Yeo et al., 2011) criteria give rise to slightly different modular parcellations but, in combination, these criteria reveal the relevant ‘building blocks’, the most important features whose variations and recombinations give rise to the complexity and diversity of the cortical tissue (Harris and Shepherd, 2015).

For convenience, these features can be coarsely (and tentatively) grouped into neuronal, synaptic and structural components (see also Getting 1989). Neuronal features refer to the different cell classes and their laminar and regional distributions (Shinomoto et al., 2009) along with their characteristic electrophysiological and biochemical diversity (Poulin et al., 2016; Tripathy et al., 2015). Synaptic components refer to a molecular default organization characterized by variations in the differential expression and transcription of genes involved in synaptic transmission (Hawrylycz et al., 2012; Pletikos et al., 2014), which is reflected, for example, in regional receptor architectonics (Palomero-Gallagher and Zilles, 2017; Zilles et al., 2002a; Zilles and Amunts, 2009). Structural aspects include variations in cortical thickness and laminar depth (Wagstyl et al., 2015) along with neuronal and synaptic density (Collins et al., 2010; Finlay and Uchiyama, 2015) and input-output (both local and long-range) connectivity patterns (Brown and Hestrin, 2009; Stepanyants and Chklovskii, 2005). In combination, these features highlight default organizational principles whose variations across the cortical sheet are likely to contribute to the corresponding functional specializations.

Naturally, these decompositions are not entirely clear or uncontroversial and, as in every biological system, tend to be fuzzy and gradual. However, their heuristic identification constitutes a necessary first step in an attempt to disentangle reusable components and mechanisms.

Based either on morphological, electrophysiological or biochemical features (or, preferably, a combination thereof), several different classes of neurons can be identified throughout the neocortex (e.g. Mensi et al. 2012; Poulin et al. 2016; Shinomoto et al. 2009; Tripathy et al. 2015; Van Aerde and Feldmeyer 2015; Zaitsev et al. 2012). Apart from pronounced regional and laminar differences in the types of neurons that make up the cortex and their relative spatial distributions, every microcircuit in every cortical column is composed of diverse neuron types, with heterogeneous properties and heterogeneous behaviour.

Electrochemical communication between these diverse neuronal classes is an intricate, dynamic and very complex process involving a multitude of nested inter- and intracellular signalling networks (Lisman et al., 2007; Marx, 2014; Südhof and Malenka, 2008). Their functional range spans multiple spatial and temporal scales (Greengard, 2001; Sabatini and Regehr, 1999; Südhof, 2013) and has, arguably, the most critical role in modulating microcircuit dynamics and information processing within and across neuronal populations (Abbott and Regehr, 2004; Duarte et al., 2017a; Gjorgjieva et al., 2016). The specificities of receptor composition and kinetics underlie the substantial diversity observed in the elicited post-synaptic potentials (Hestrin, 1993; Voglis and Tavernarakis, 2006) across different synapse and neuronal types (see, e.g. Angulo et al. 1999; Destexhe et al. 1994, 1998; Moreau and Kullmann 2013; Nissen et al. 2010). This occurs because the receptors mediating these events have distinct biochemical and physiological properties depending on the type of neuron they are expressed in and, naturally, the type of neurotransmitter they are responsive to. Having very different genetic make-ups, different neuronal classes contain synapses with different properties in terms of, for example, the total number and relative ratios of different receptor types (Connors et al., 1988; Myme, 2003; Watt et al., 2000) and their specific sub-unit composition (Greger et al., 2017; Henley and Wilkinson, 2016; Kühr et al., 1993; Paoletti et al., 2013). These varying properties have known and non-negligible implications in the characteristic kinetics of synaptic transmission events occurring between different neurons (Kubota et al., 2016) and strongly constrain the circuit’s operations.

Additionally, cortical microcircuits are not randomly coupled, but over-express specific connectivity motifs (Perin et al., 2011; Shimono and Beggs, 2015; Song et al., 2005; Thomson, 2002; Yoshimura and Callaway, 2005; Yoshimura et al., 2005), which bias and skew the network’s degree distributions (Tomm et al., 2014) and/or introduce correlations among specific connections (Koulakov et al., 2009), thus selectively modifying the impact of specific pre-synaptic neurons on their post-synaptic targets. Analogously to the heterogeneities in neuronal and synaptic properties, such structural features are known to significantly impact the circuit’s properties (Harris and Mrsic-Flogel, 2013; Hoffmann and Triesch, 2017; Litwin-Kumar and Doiron, 2012; Pernice et al., 2013; Roxin, 2011).

### 1.2 Descriptive adequacy

As discussed above, the variations in the anatomy and physiology of a cortical microcircuit are experimentally well established and have been shown to influence computational properties. This notwithstanding, they are rarely accounted for in computational studies. One reason for this lies in our current inability to unambiguously uncover the ground truth. For example, despite significant progress, both methodological (Cardin, 2012; Deisseroth et al., 2006; Miesenböck, 2011) and empirical (Helmstaedter et al., 2008, 2009; Wertz et al., 2015; Yoshimura and Callaway, 2005; Yoshimura et al., 2005), it is still technically challenging to account for a sufficiently large number of synapses per neuron or even for a sufficiently large number of the neurons that make up the circuit to entirely disentangle the nature of structural connectivity and heterogeneity.

In the absence of complete descriptions and well-substantiated empirical evidence to support them, a simplified approach is often taken. This has multiple additional advantages, such as greater likelihood of analytical tractability and lower overhead for the researcher in specifying the network parameters. Such simplified, homogeneous models of spiking networks have proven to be a valuable tool for a theoretically grounded exploration of microcircuit dynamics, emerging from the interaction of excitatory and inhibitory populations (Amit and Brunel, 1997a; Tsotsos et al., 2000; van Vreeswijk and Sompolinsky, 1996, 1998).

The generic principles established by studying simple balanced random networks have subsequently been applied to model specific cortical microcircuits with integrated connectivity maps and realistic numbers of neurons and synapses (Potjans and Diesmann, 2014). This approach revealed that some prominent features of spontaneous and stimulus-evoked activity and its dynamic flows through a cortical column can be accounted for by the macroscopic connectivity structure, mediated by local and long-range interactions (Schmidt et al., 2015). However, by focusing on emergent dynamics, these studies neglect the functional aspects and the fact that cortical interactions serve computational purposes (but, see Cain et al. (2016) for a study on the computational properties of the Potjans and Diesmann (2014) microcircuit model). In addition, although there are good reasons for taking a minimalist approach, assuming uniformity and homogeneity on every component of the system tends to lack cogency with respect to established anatomical and physiological facts and to disregard biophysical and biochemical plausibility.

Some of these limitations were circumvented by Haeusler and Maass (2007), who not only accounted for detailed and empirically-informed connectivity maps, but also employed more biologically motivated models of neuronal and synaptic dynamics and placed them in an explicit functional/computational context. In line with the results obtained with the simpler microcircuit models (Cain et al., 2016; Potjans and Diesmann, 2014), this study demonstrated that considering realistic structural constraints is beneficial and significantly improves the computational capabilities of the circuit.

A completely different set of priorities for modelling cortical microcircuits are espoused by Markram et al. (2015) in the framework of the continuous efforts of the Blue Brain project (Markram, 2006). The Blue Brain approach lies on the other extreme of the descriptive scale, in that it attempts to model a cortical column in full detail, explicitly accounting for the complexities of cellular composition (based on neuronal morphology and electrophysiology), synaptic anatomy and physiology, as well as thalamic innervation, essentially constituting an *in silico* reconstruction of a cortical column (see also Helmstaedter et al. 2007). This approach is, naturally, extremely computationally expensive and its explanatory power is limited. The model complexity at this end of the spectrum is so close to the biophysical reality that it might not lend itself to a comprehensive understanding of dissociable and important functional principles any more readily than studying the real thing does. Nonetheless, it provides valuable insights in that it carefully replicates a lot of *in vivo* and *in vitro* responses of a real cortical column, while generating a wealth of complete and comprehensive data (Markram et al., 2015; Ramaswamy et al., 2015).

Thus, we conclude that while simpler models are preferable, as they are generic enough to be broadly insightful and allow us to uncover general principles, we should ask the question: what is the cost of simplification? If a model simplifies away the core computational elements of the system, our ability to account for its operations is lost. The findings discussed above indicate that heterogeneity may be critical for the mechanisms of computation; therefore models aiming at uncovering computational principles in specific biophysical systems, such as a cortical column or microcircuit, should at least account for these features.

In this study, we attempt to bridge this descriptive gap by building microcircuit models, inspired and constrained by the composition of Layer 2/3, that account for key heterogeneities in neuronal, synaptic and structural properties. We implement all types of heterogeneity such that they can be switched on or off, thus enabling us to systematically disentangle and evaluate the roles played by the different types of heterogeneity in the different tentative building blocks, and how they collectively interact to shape the circuit’s dynamics and information processing capacity.

The choices and characteristics of the models and parameter sets used throughout this study, as well as the general microcircuit composition are constrained and inspired by multiple sources of experimental data (see Section 2 and Supplementary Materials) and account for the prevalence of different neuronal subtypes and their heterogeneous physiological and biochemical properties, the specificities of instantaneous synaptic kinetics and its inherent diversity as well as specific structural biases in cortical micro-connectivity. All models and model parameters were, as far as possible, chosen to directly match relevant experimental reports and minimize the introduction of arbitrary model parameters, in order to ensure that the effects observed are caused by realistic forms of complexity and heterogeneity and avoid imposing excessive assumptions or preconceptions on the systems studied, i.e. to “allow biology to speak for itself”.

The circuits explored are not meant to provide an exact, detailed replica of a cortical microcircuit, but instead an intermediate stage of descriptive accuracy in between the abstract, simplified formalisms and the highly detailed ones, while retaining a primary focus on computation and functional aspects. The goal is to account for key features that are bound to play a significant role in the circuit’s dynamics and information processing properties while still maintaining a manageable degree of model/parameter complexity and computational cost. This approach also serves as a “proof of concept” to demonstrate that it is possible to explicitly use neurophysiological and biophysical insights to build and parameterize complex microcircuit models at the level of point-neuron models, as well as to devise and apply complex benchmark measurements which, being system-independent, allow us to directly compare systems with varied degrees of biological plausibility and thus systematically assert the functional and computational implications of several architectural features present in biology as well as the emergent phenomena they entail.

In Section 2 (complemented by Section 6 and the Supplementary Materials), we explain all the details of the models and model parameters used to build and constrain the microcircuit, as well as the underlying empirical observations that motivate the choices. After specifying and fixing all the relevant parameters to, as closely as possible, match multiple sources of empirical data, we study the effects of heterogeneity on population dynamics in a *quiet* state, where the circuit is passively excited by background noise (Section 3) and in an *active* state, where the circuit is directly engaged in information processing (Section 4). We evaluate the circuit’s sensitivity and responsiveness, as well as its memory and processing capacity, demonstrating a clear and unambiguous role of heterogeneity in shaping the proficiency of the system by greatly increasing the space of computable functions. Neuronal heterogeneity is shown to be the leading source of functional specialization, having a clear and marked impact on microcircuit dynamics. Structural heterogeneity, on the other hand, has marginal effects on population activity, but greatly increases the circuit’s capacity to process highly nonlinear functions of the input.

## 2 Building the microcircuits

In this section, we describe the process of building a complex data-driven cortical microcircuit model capturing some of the fundamental features of layer 2/3. We specify the detailed architecture, composition and dynamics of the microcircuits explored throughout this study as well as the motivation behind all model and parameter choices. In each relevant section, we highlight the differences between the respective homogeneous and heterogeneous conditions. A summarized, tabular description of the main models is provided in the Supplementary Materials, along with a list of the primary sources of experimental data used to constrain the model parameters.

All the circuits analysed throughout this study are composed of N = 2500 point neurons (roughly corresponding to the size of a layer 2/3 microcircuit in an anatomically defined cortical column; Lefort et al. 2009), divided into N_E_ = 0.8 N excitatory, glutamatergic neurons and N_I_ = 0.2 N inhibitory, GABAergic neurons. In addition, we further subdivide the inhibitory population in two sub-classes, I_1_ and I_2_ (with N_I_1__ = 0.35 N_I_ and N_I_2__ = 0.65 N_I_), corresponding to fast-spiking and non-fast-spiking interneurons, respectively (see Section 2.1). Accordingly, there are nine different synapse types (all possible connections between neuronal populations), with distinct, specific response properties (see Section 2.2). Similarly, there are nine connection probabilities from which random connections are drawn (see Section 2.3).

For each of the key features of neuronal, synaptic and structural properties, we differentiate between the homogeneous case, where all properties are identical, and the heterogeneous case, where properties are drawn from appropriately chosen distributions. In this way, we can tease apart the differential effects of the three sources of heterogeneity considered here: neuronal, synaptic and structural.

For consistency, all the circuits’ structural (and synaptic) features are constrained primarily by the composition of layer 2/3 in the C2 barrel column in the mouse primary somatosensory cortex (S1), given the availability of direct, complete and significantly explored experimental datasets (e.g. Avermann et al. 2012; Gentet et al. 2010; Lefort et al. 2009).

### 2.1 Neuronal properties

We divide the neurons into one excitatory (E: glutamatergic, pyramidal neurons) and two inhibitory (I_1_: GABAergic, fast spiking interneurons, I_2_: GABAergic, non-fast spiking) classes, consistent with the reports in Avermann et al. (2012), Lefort et al. (2009) and Gentet et al. (2010). The three classes differ in relative excitability and firing properties, providing substantially more electrophysiological diversity than commonly exhibited in cortical models on the abstraction level of point neurons (e.g. Amit and Brunel 1997b; Ecker et al. 2010; Potjans and Diesmann 2014; Renart et al. 2010; Shadlen and Newsome 1998), while still being of a manageable degree of complexity. All neurons are modelled using a simplified adaptive leaky integrate-and-fire scheme (see Section 6.1 and Gerstner et al. 2014; Koch 2004).

The parameters used for the different neuron types (summarized in Table 1) were chosen to match the respective ranges reported in the literature, considering both the data collected in the NeuroElectro^1^ database (Tripathy et al., 2014, 2015) encompassing tens of unique data points from different experimental sources (see Supplementary Materials) and those reported in Avermann et al. (2012); Harrison et al. (2015); Lefort et al. (2009); Lu et al. (2007), and Gentet et al. (2010) given the completeness of these reports and the direct similarities with our case study.

**Table 1.**
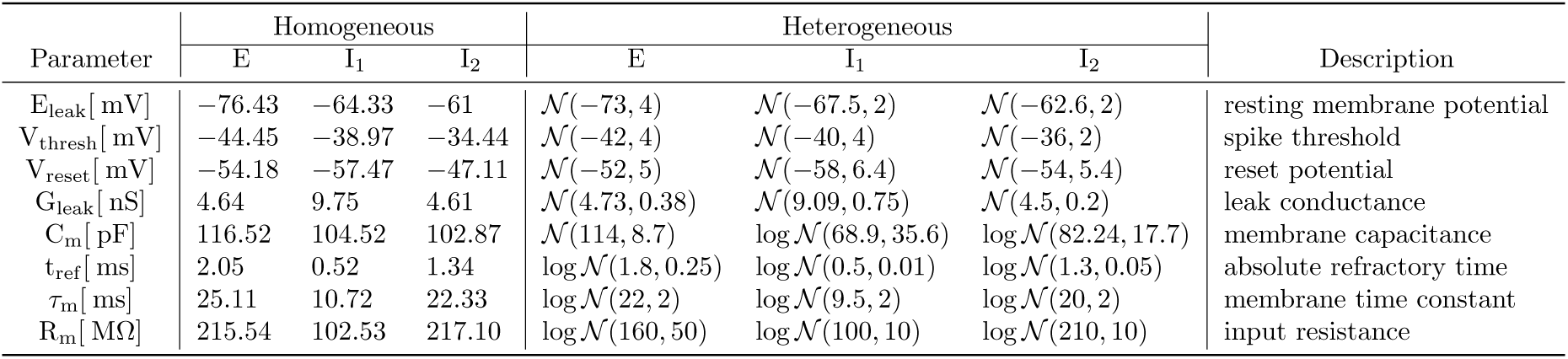
Single neuron parameter sets. In the heterogeneous condition, three neuronal types have the values of the described parameters randomly drawn from a normal (𝒩) or lognormal (log 𝒩) distribution. The parameter values for these distributions were determined taking into account multiple sources of experimental data (see Supplementary Materials). Note that, to make comparisons simpler, the values displayed for the lognormal distributions correspond to the mean and standard deviation of the distribution, not the actual *μ* and *σ* parameters (see Section 6).

Due to the nature of the chosen neuronal formalism (Section 6.1), some model parameters have no direct proxy with experimental measures and were instead determined considering their relations to other variables for which direct experimental data exists. For example, V_reset_ is not, strictly speaking, a biophysically meaningful variable; its value was chosen considering the data for afterhyperpolarization potentials E_AHP_ relative to the resting membrane potentials E_L_. Whenever such discrepancies occurred or when parameters had to be derived relative to others, we selected those values that better matched the experimental data. Overall, we observed a remarkable consistency in the ranges of parameter values considered across the different data sources.

#### 2.1.1 Neuronal heterogeneity

In the homogeneous condition, the parameters for all neurons of a given class are fixed and chosen as a representative example of that class (see left side of Table 1 and bold fI curves in Figure 1). To incorporate neuronal heterogeneity, the specific values of each parameter are independently drawn from a probability distribution, specific to each neuron class (see right side of Table 1 and individual fI curves in Figure 1).

**Figure 1.**
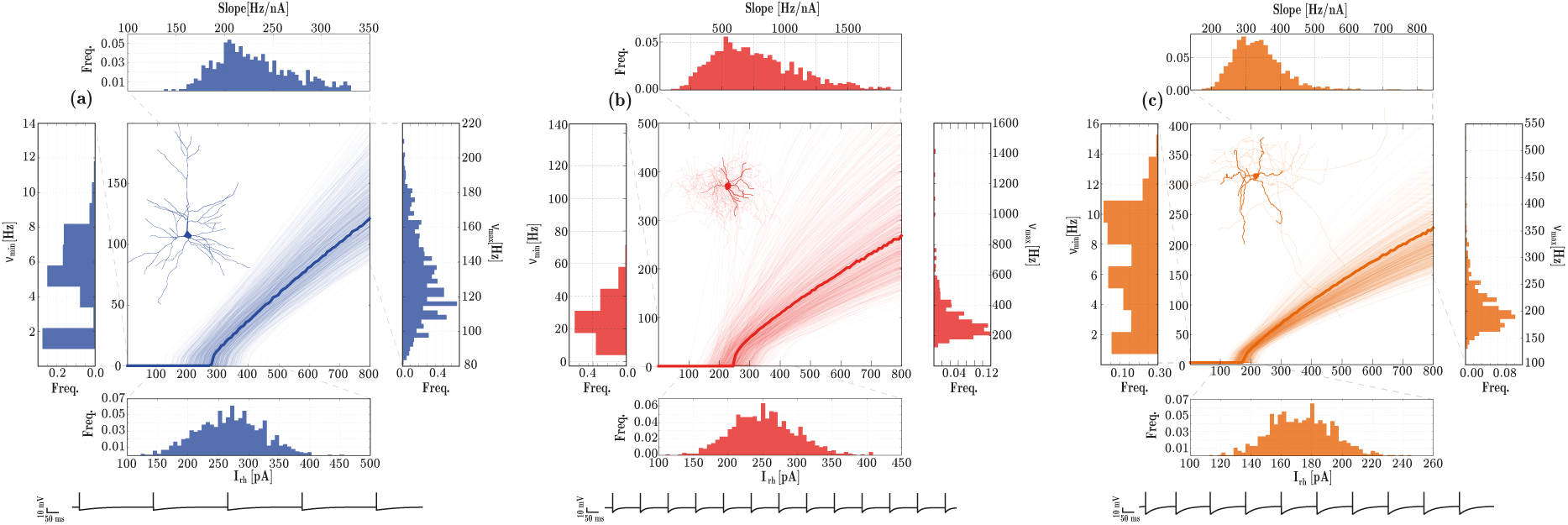
Response properties of the three different neuronal types, E (left), I_1_ (middle) and I_2_ (right). For each neuronal class, the central panels depict single neuron fI curves and the marginal panels give the corresponding distributions of the neuron’s rheobase currents (I_rh_[pA], bottom), minimum firing rates (*v*_min_[*H_z_*], left); maximum firing rates (*v*_max_[*H_z_*], right) and the slope of the fI curve (Slope[Hz/nA], top). The data was obtained from 1000 neurons of each class. The membrane potential traces depicted in the bottom correspond to the response of the homogeneous neurons (bold traces in the fI curves) to a stimulus step of amplitude I_rh_ + 10 pA, for a duration of 10 seconds.

To ensure comparability among conditions, we tuned the intrinsic adaptation parameters (*a*, *b*, see Section 6.1 and Supplementary Materials) independently for each neuron type (data not shown), in order to retain the following relations among neuronal classes:

- Rheobase current (excitability) - *I*_rh_(E) ≈ *I*_rh_(I_1_) > *I*_rh_(I_2_)
- Slope of f-I curve (gain) - *g*(I_1_) > *g*(I_2_) > *g*(E)
- Minimum firing rate - *v*_min_(I_1_) > *v*_min_(I_2_) > *v*_min_(E)
- Maximum firing rate - *v*_max_(I_1_) > *v*_max_(I_2_) > *v*_max_(E)

This led to the following values (*a*, *b*) for sub-threshold and spike-triggered adaptation, respectively: E = (4, 30), I_1_ = (0, 0), I_2_ = (2, 10).

One of the advantages of using the modelling approach presented in this study is the possibility to directly interpret the biophysical meaning of the different parameters. In this case, these results highlight the fact that fast spiking interneurons (I_1_) do not exhibit any form of intrinsic adaptation, which is a reasonable result for a neuron whose primary role is to respond quickly and provide dense and fast, feed-forward inhibition (Szabadics, 2006). It should be noted that, after fixing all the neuronal parameters as described above, the absolute values of these four properties did not exactly match the corresponding experimental reports. For example, the values obtained for the rheobase currents were, on average, larger than the ranges reported experimentally, for all 3 neuron classes (see Supplementary Materials).

### 2.2 Synaptic properties

With three neuronal populations, as described above, there are nine possible types of synaptic connection, i.e. syn ∈ { EE, E I_1_, E I_2_, I_1_ E, I_2_ E, I_1_ I_1_, I_1_ I_2_, I_2_ I_1_, I_2_ I_2_}. Synapse types can be grouped by transmitter composition and/or post-synaptic effect, as excitatory syn_E_ = { EE, I_1_ E, I_2_ E} and inhibitory syn_I_ = { EI_1_, EI_2_, I_1_ I_1_, I_1_ I_2_, I_2_ I_1_, I_2_ I_2_}, as illustrated in Figure 2a. For simplicity, we consider all synaptic transmission as being mediated by either glutamate (excitatory synapses) or gamma-aminobutyric acid (GABA, inhibitory synapses), as illustrated in Figure 2b. This is a reasonable simplification given that these are, by far, the primary neurotransmitters used in the neocortex, as demonstrated by immunohistochemistry (Hill et al., 2000) and receptor autoradiography studies (Palomero-Gallagher et al., 2015; Zilles et al., 2002a, b). Additionally, this is a common assumption underlying the great majority of theoretical and computational studies.

**Figure 2.**
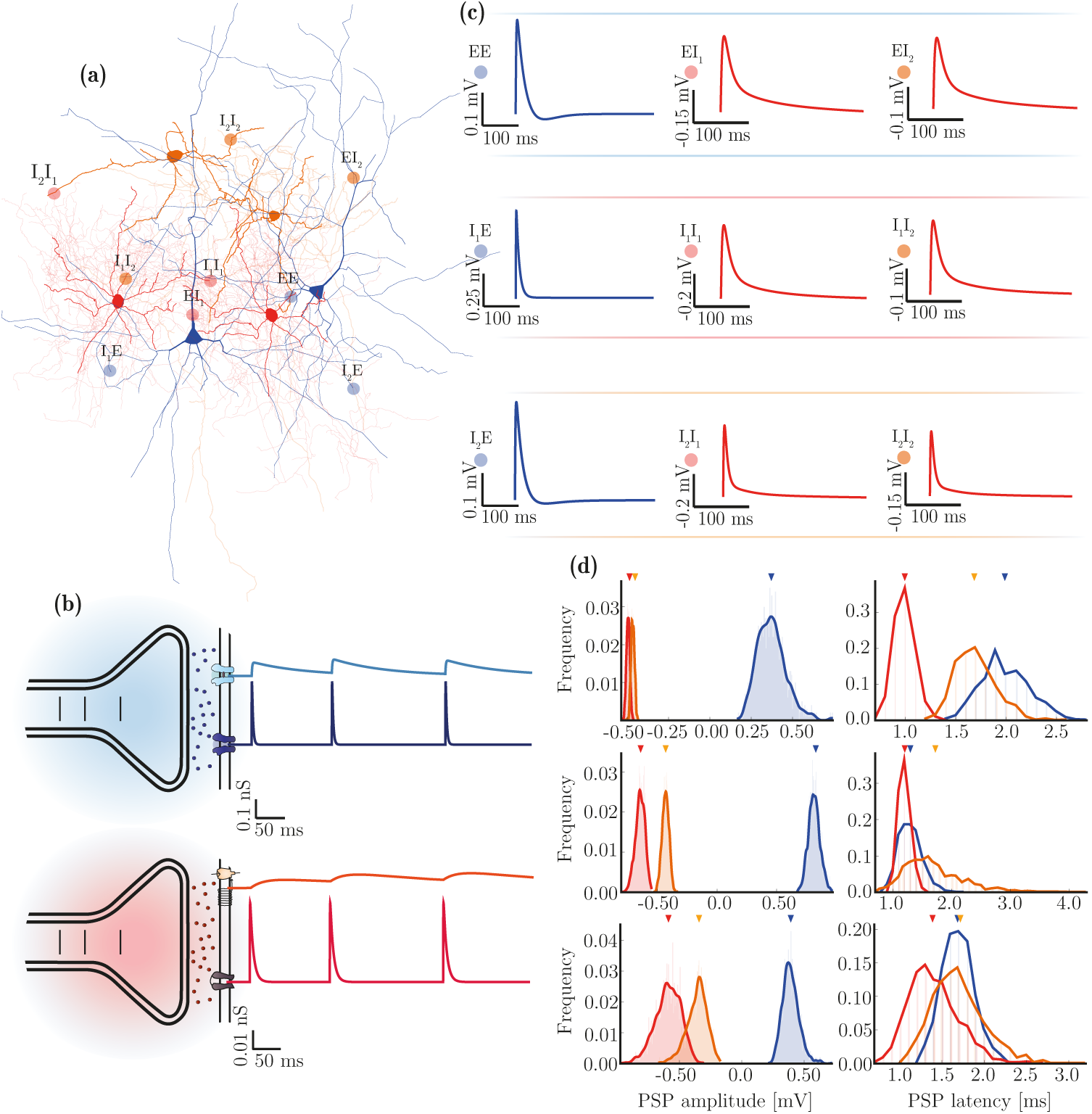
Diversity of synaptic transmission in the microcircuit. **(a)** Illustration of the three neuron and nine synapse types considered in this study. **(b)** Characteristics of synaptic transmission in excitatory (blue) and inhibitory (red) synapses, comprising fast and slow components. **(c)** Kinetics of spike-triggered PSPs dependent on pre- and post-synaptic neuron type and determined by the specific receptor kinetics and composition in the postsynaptic neuron. The depicted traces are the PSPs at rest in E neurons (top row) and at a fixed holding potential of –55 mV for inhibitory neurons (I_1_: middle row, I_2_: bottom row), as a function of receptor composition and correspond to the values reported in Table 2 (last column). **(d)** Distributions of PSP amplitudes and latencies after rescaling (by *w* ^syn^ and *d*^syn^, respectively, see Table 3) in the homogeneous (top arrows) and heterogeneous (distributions) conditions, for synapses onto E (top), I_1_ (middle) and I_2_ (bottom) neurons. Note that the latency distributions are discrete in that for technical reasons, they can only assume values that are a multiple of the simulation resolution.

**Table 2.**
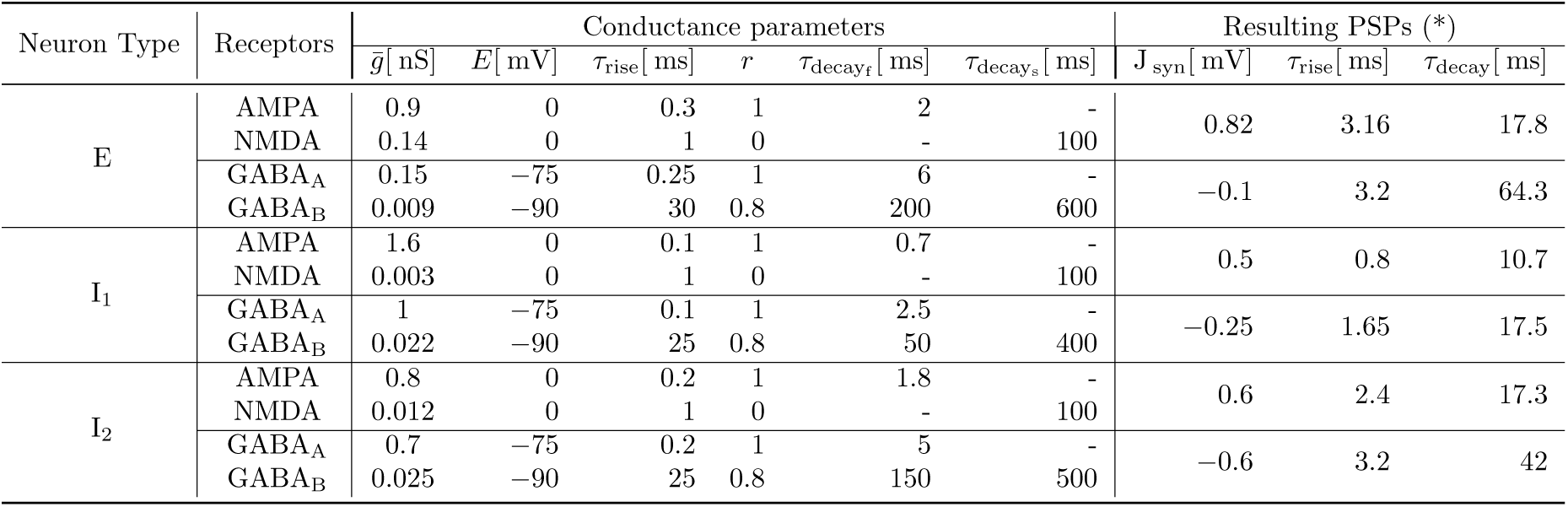
Differential receptor expression in the different neuron types. The kinetics and relative conductance of the different receptors that make up an inhibitory or excitatory synapse onto each neuron results in post-synaptic potentials with equally discernible kinetics. The parameters were chosen based on the corresponding receptor conductance data, if directly available and/or on the characteristics of the resulting PSPs, resulting in a substantial diversity of postsynaptic responses (see Figure 2c). (*) The PSP values reported in this table were obtained by fitting a double exponential function to single, spike-triggered PSPs, recorded at rest for E neurons and at a fixed holding potential of –55 mV for I neurons.

| Neuron Type | Receptors | Conductance parameters |  |  |  |  |  | Resulting PSPs (*) |  |  |
| --- | --- | --- | --- | --- | --- | --- | --- | --- | --- | --- |
| | | $\bar{g}$ [nS] | $E$ [mV] | $\tau_{\text{rise}}$ [ms] | $r$ | $\tau_{\text{decay}_f}$ [ms] | $\tau_{\text{decay}_s}$ [ms] | $J_{\text{syn}}$ [mV] | $\tau_{\text{rise}}$ [ms] | $\tau_{\text{decay}}$ [ms] |
| E | AMPA | 0.9 | 0 | 0.3 | 1 | 2 | - | 0.82 | 3.16 | 17.8 |
|  | NMDA | 0.14 | 0 | 1 | 0 | - | 100 |  |  |  |
|  | GABA<sub>A</sub> | 0.15 | -75 | 0.25 | 1 | 6 | - | -0.1 | 3.2 | 64.3 |
|  | GABA<sub>B</sub> | 0.009 | -90 | 30 | 0.8 | 200 | 600 |  |  |  |
| I<sub>1</sub> | AMPA | 1.6 | 0 | 0.1 | 1 | 0.7 | - | 0.5 | 0.8 | 10.7 |
|  | NMDA | 0.003 | 0 | 1 | 0 | - | 100 |  |  |  |
|  | GABA<sub>A</sub> | 1 | -75 | 0.1 | 1 | 2.5 | - | -0.25 | 1.65 | 17.5 |
|  | GABA<sub>B</sub> | 0.022 | -90 | 25 | 0.8 | 50 | 400 |  |  |  |
| I<sub>2</sub> | AMPA | 0.8 | 0 | 0.2 | 1 | 1.8 | - | 0.6 | 2.4 | 17.3 |
|  | NMDA | 0.012 | 0 | 1 | 0 | - | 100 |  |  |  |
|  | GABA<sub>A</sub> | 0.7 | -75 | 0.2 | 1 | 5 | - | -0.6 | 3.2 | 42 |
|  | GABA<sub>B</sub> | 0.025 | -90 | 25 | 0.8 | 150 | 500 |  |  |  |

**Table 3.**
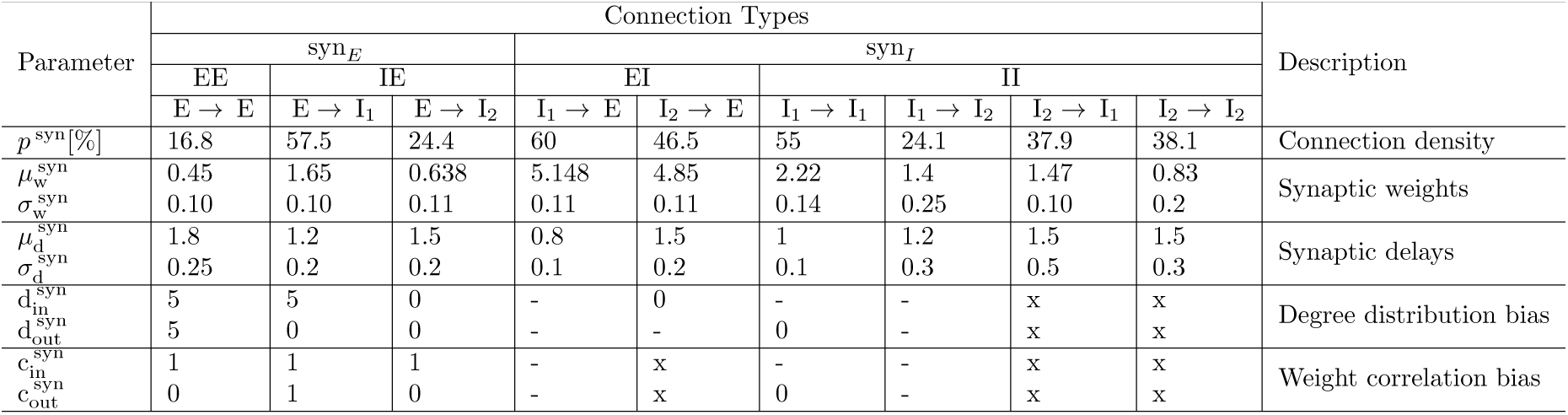
Synaptic and structural parameters in the microcircuit. Each of the nine connection types (which can be grouped as indicated) is characterized by a specific connection density, weight and delay. In the homogeneous condition, weights and delays are fixed and equal to the mean values 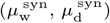 for all synapses of a given type, whereas in the heterogeneous condition they are independently drawn from lognormal distributions with the corresponding mean and standard deviation. The last two rows in the table are the connection-specific structural bias parameters, used to skew the network’s degree and weight distributions. The indicated values were taken directly from Tomm (2012) and Tomm et al. (2014). The cases marked with - or x, correspond to connections that were either tested, revealing no significant effect (-) or untested due to missing data (x). In both cases, we set the corresponding values to 0.

To accommodate the wealth of data available regarding the phenomenology of synaptic transmission and to provide a significant step forward from the traditional approaches, we chose a relatively complex biophysical model (Gerstner and Kistler, 2002; Gerstner et al., 2014; Hoffmann et al., 2015; McCormick et al., 1993); primarily due to its plausibility but also due to the availability of direct parameters in the experimental literature (e.g. Hoffmann et al. 2015). The model, described fully in Section 6.2, captures the detailed kinetics of single receptor conductances (Figure 2b).

The use of this model allows us to specify different receptor parameters depending on the neuron type they are expressed in (see Table 2), in order to directly match empirical data on receptor kinetics and relative conductance ratios in different neuronal classes. In the absence of such direct data for specific synapses (particularly those involving I_2_ neurons), we tuned these parameters to match the resulting PSP/PSC kinetics, as well as the relative ratios of total charge elicited by the receptors that compose such synapses. For a detailed list of the data sources used to constrain these parameters, consult the Supplementary Materials.

Having fixed the kinetics of post-synaptic responses according to neuron class (Table 2), we finally rescale the PSP amplitudes (*w*^syn^) and delays (*d*^syn^) independently for each synapse type (see Table 3), in order to account for the effects of different presynaptic neuron classes and to explicitly match the data reported in (Avermann et al., 2012). As a result of this parameter fitting process, the responses generated by the synaptic model are good matches to the responses experimentally observed in the nine types of biological synapses represented in this study.

#### 2.2.1 Synaptic heterogeneity

As all the receptor parameters are fixed and neuron-specific (Table 2), we introduce synaptic heterogeneity by simply distributing the individual values of weights and delays (Figure 2d and Table 3). Whereas in the homogeneous condition, synaptic efficacies (*w*^syn^) and conduction delays (*d*^syn^) are fixed and equal for all connections of a given type, in the heterogeneous condition, these values are randomly drawn from lognormal distributions, left-truncated at 0 for weight distributions and 0.1 for delay distributions, and parameterized such that the distributions’ means are equal to the homogeneous value. For consistency among the various data sources, we fix the connectivity parameters, including not only structural aspects, but also synaptic weights and delays, to match the data reported in (Avermann et al., 2012).

For that purpose, the mean weight values were chosen to rescale the PSP amplitude of each synapse type to the target value. Additionally, as described below in Section 2.3, if both structural and synaptic heterogeneity conditions are considered simultaneously, the weight distributions are skewed in order to introduce structural weight correlations (see Section 6.3.1).

### 2.3 Structural Properties

The structure of the network is defined by the density parameter *p*^syn^, which is specific for each of the nine connection types. For consistency with the results and methods presented in Tomm (2012) and Tomm et al. (2014), we set the connection densities (*p*^syn^) to the values reported in Avermann et al. (2012) (see Table 3). However, it is worth noting that the values reported in the literature can vary substantially, possibly reflecting methodological/technical limitations and/or the fact that connection density is a highly region- / species-specific feature. In fact, among all the complex parameter sets used throughout this study, the single parameter that was most difficult to reconcile across multiple sources was *p*^syn^. In the homogeneous case, random connections are created between neurons in a source population *pre* and target population *post* (with *pre*,*post* ∊ { E, I_1_, I_2_}) with a probability given by *p*^syn^.

#### 2.3.1 Structural heterogeneity

In order to account for structural heterogeneity, we bias the network’s degree distributions by modifying the structure of the connectivity matrix *A*^syn^, following the methods introduced in (Tomm, 2012; Tomm et al., 2014) and validated against the same primary sources of experimental data used in this study.

By controlling the skewness of the out-/in-degree distributions (d^out/in^ parameters, see Section 6.3), we can generate completely random, uniform connectivity (d^out/in^ = 0, Figure 3a) or highly structured in-/out-degree distributions, with a larger variance in the number of connections per neuron (d^out/in^ > 0, Figure 3b). For the structural heterogeneity condition, these parameters were fixed to the values that were shown to provide a better fit for the experimentally determined connectivity data (see Tomm 2012; Tomm et al. 2014 and Table 3).

**Figure 3.**
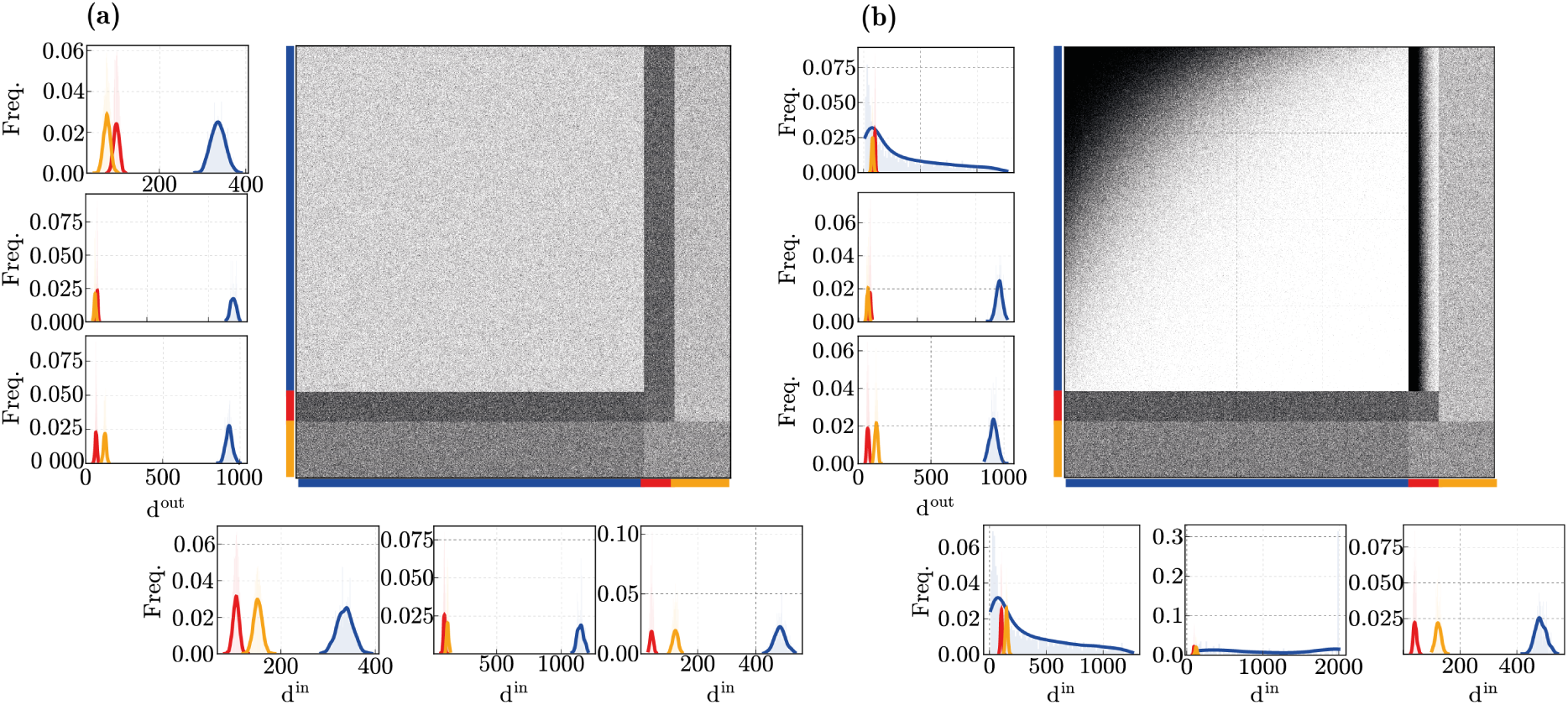
Microcircuit connectivity for **(a)**homogeneous and **(b)** heterogeneous network structures. The large panels show the circuit’s complete connectivity matrix *A*^syn^, comprising all connections among all neuron classes. The panels on the left sides show the corresponding out-degree (d^out^) distributions for the neuronal populations (E, I_1_ and I_2_ from top to bottom); the panels at the bottom show the corresponding in-degree (d^in^) distributions (E, I_1_ and I_2_ from left to right)

Additionally, in conditions where synaptic heterogeneity is also present (fully heterogeneous circuit), structural heterogeneity is further expressed as a bias in the synaptic efficacies for all incoming and outgoing connections to a given neuron. Following Tomm (2012), Tomm et al. (2014) and Koulakov et al. (2009), this bias is implemented by rescaling individual synaptic efficacies in order to introduce correlations between them, see Table 3 and Section 6.3.1.

## 3 Emergent population dynamics

In this section, we set out to quantify and evaluate the specific impact of the different forms of heterogeneity (neuronal, synaptic, structural, and the combination of all forms) on the characteristics of population activity. To do so, we consider the circuit’s responses to an unspecific and stochastic external input, modelling cortical background / ongoing activity (see Section 6.4.1). We determine and compare the circuit’s responsiveness by looking at the population rate transfer functions, as exemplified in Figure 4a for I_2_ neurons (complete results are provided in the Supplementary Materials), and summarize the results by the change in absolute gain (ΔGain) and offset (ΔOffset) introduced by each source of heterogeneity, relative to the homogeneous condition (Figure 4b).

**Figure 4.**
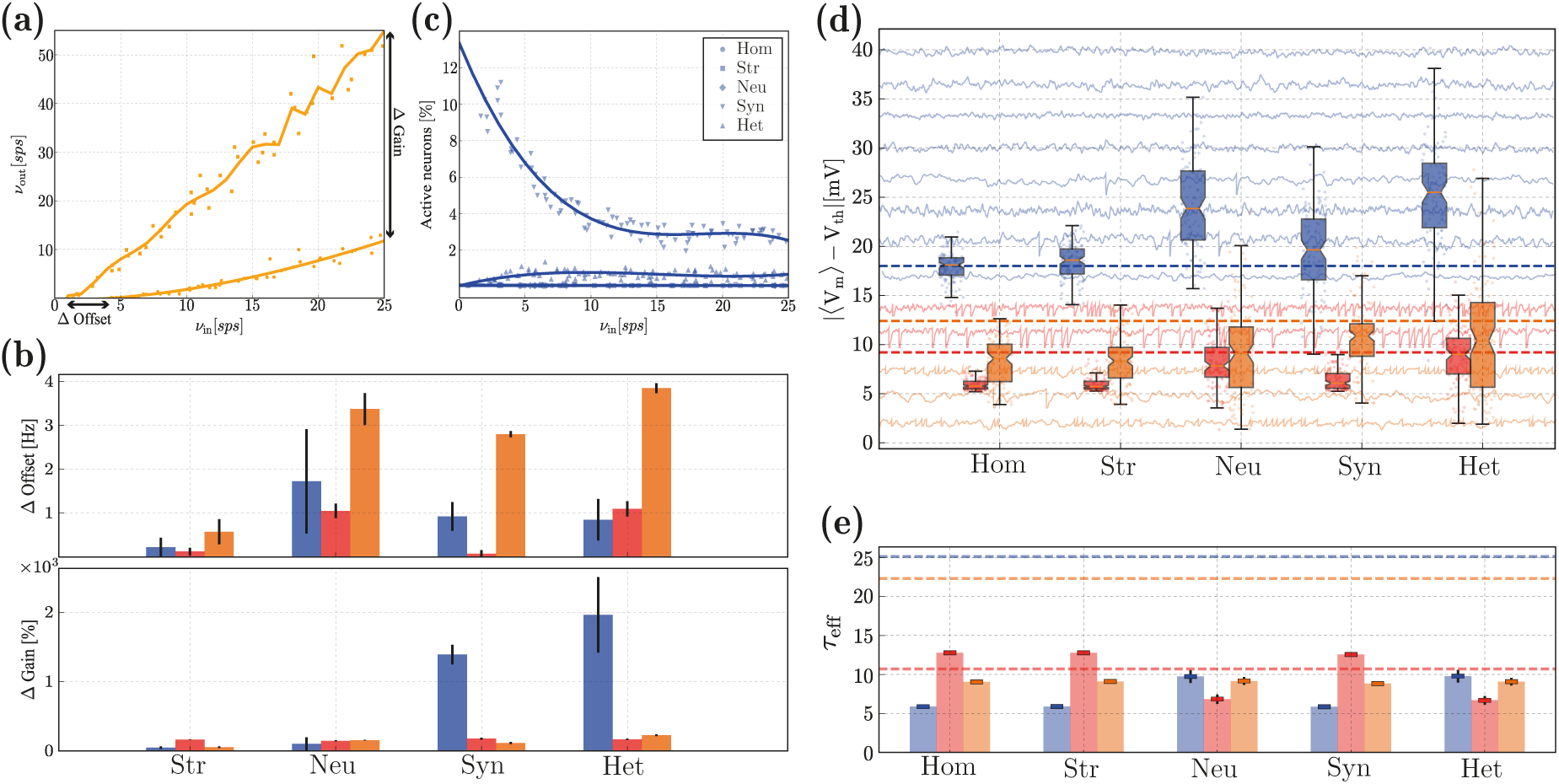
Characteristics of population activity in the *quiet* state. **(a)**: Rate transfer function of the I_2_ neurons in the homogeneous (dots) and structurally heterogeneous (squares) conditions. **(b)**: Change in absolute gain (ΔGain) and offset(ΔOffset), see annotations in (a), for the E (blue), I_1_ (red) and I_2_ (orange) populations, for each heterogeneous condition (structural, neuronal, synaptic, and all forms combined), relative to the homogeneous condition. Error bars indicate the standard deviation across ten simulations per condition. **(c)**: Fraction of active E neurons in the different conditions as a function of input rate *ν_in_*. **(d)**: Distributions of absolute distances between neurons’ mean membrane potentials 〈V_m_〉 and their firing threshold V_th_, for the three neuronal populations considered, under the different heterogeneity conditions. The dashed lines indicate the corresponding values reported in (Gentet et al., 2010). For illustrative purposes, we depict *V_m_* traces for a small set of randomly chosen neurons in the background. **(e)**: Effective membrane time constants (*τ*_eff_ = 〈G_total_〉/ C_m_) in the noise-driven regime, for the different neuronal classes and conditions. Error bars correspond to the means and standard deviations across each population. The dashed lines indicate the baseline values (*τ*_o_ = g_leak_/ C_m_). All results depicted were calculated over an observation period of 10 s. The results depicted in **(d, e)** were acquired with a fixed input rate of *ν_in_* = 10 spikes/s.

All heterogeneous conditions, particularly neuronal and synaptic, cause a slight offset for all neuron types (more significant for I_2_ neurons), making them slightly more responsive (firing at lower input rates) but the effect is not substantial (Figure 4b, top). In most of the conditions analysed, the E population is rather unresponsive, with less than 1% of the neurons active (Figure 4c) and firing at rates inferior to 1 spikes/s, regardless of the input rate. While structural and neuronal heterogeneity are incapable of circumventing this effect, synaptic heterogeneity appears to be strictly required for the network to fire at more reasonable rates (albeit, still very sparsely) and for the E population to become responsive to variations in the input rate, resulting in a significant modulation of the gain of the rate transfer function (Figure 4b, bottom).

It should be noted that the impact of structural heterogeneity alone is mitigated by the low E rates, since the structural bias exists only within excitatory synapses or between excitatory neurons and fast-spiking interneurons (i.e. EE, I_1_ E, see Table 3). So, if the E population rarely fires, it is difficult to ascertain the effects of structural heterogeneity, suggesting either that its relevance pertains mostly to active states, when population activity is slightly higher, or that it is negligible at this scale.

The extremely sparse firing of E neurons that we observe is consistent with physiological measurements in layer 2/3 (e.g. Benedetti et al. 2013; Crochet et al. 2011; Gentet et al. 2010; Lefort et al. 2009; O’Connor et al. 2010; Petersen and Crochet 2013), but it significantly limits the degree to which we can quantify the effects of heterogeneity on population activity. So, in order to obtain a greater insight, we look at the sub-threshold responses and characteristics of membrane potential dynamics (Figure 4d, e). Excitatory neurons are always significantly hyperpolarized, with their mean membrane potentials kept far from threshold (Figure 4d, blue) and thus require much stronger depolarizing inputs to fire, compared with both inhibitory types. The inhibitory populations are, on average, much more depolarized and their membrane potentials fluctuate closer to their firing thresholds, particularly I_1_ (Figure 4d, red). Qualitatively, the ratio of average degree of depolarization among the different populations is retained across all conditions, with I_1_ neurons being strongly depolarized, followed by I_2_ and E and is consistent with experimental reports for circuits in a state of *quiet wakefulness* (Figure 4d, dashed lines). This feature stems directly from the electrophysiological properties of the different neuronal classes and the interactions among the 3 populations (given that it is already observed in the homogeneous circuit). Both synaptic and neuronal heterogeneity greatly increase the variability in the distribution of mean membrane potentials across all the neurons and cause a significant overlap between E and I_2_ populations, an effect that is also consistent with experimental evidence (Gentet et al., 2010).

The relative impact of intense synaptic bombardment on the integrative properties of the different neuronal classes is also critically influenced by the presence of neuronal heterogeneity, as illustrated by the distribution of effective membrane time constants (Figure 4e). Active synapses contribute to the total membrane conductance and cause a significant deviation from the resting membrane time constant (Destexhe et al., 2003; Kumar et al., 2008). In the absence of synaptic input, I_1_ neurons have faster responses, characterized by a short baseline membrane time constant (*τ*_0_ = g_leak_/ C_m_ ≈ 10.7ms), whereas I _2_ and E neurons are slower (*τ*_o_ ≈ 22.3 and 25.1 ms, respectively) and can thus integrate their synaptic inputs over a larger time scale. This relationship between the neuronal classes (*τ*_eff_ (I_1_) < *τ*_eff_ (I_2_) < *τ*_eff_ (E)) is a consequence of the neurons’ physiological properties (and consistent with empirical evidence Angulo et al. 1999; Destexhe et al. 1999; Leger et al. 2005). It is thus important that, under conditions of intense synaptic bombardment, the different neuronal classes respond consistently and retain this ratio. This is only observed in the presence of neuronal heterogeneity (Figure 4e).

### 3.1 Excitation / inhibition balance

The balance of excitation and inhibition is one of the most important and widely observed features in the neocortex. It plays a pervasive role in modulating and stabilizing circuit dynamics (Humphries, 2016), shifting the population state (Poulet, 2014; Tsodyks and Sejnowski, 1995; Zucca et al., 2017), selectively gating and routing signals (Kremkow et al., 2010; Vogels, 2005; Vogels and Abbott, 2009) and maintaining sparse, distributed dynamics (Crochet et al., 2011; Waters and Helmchen, 2006), critical for adequate processing and computation (Deneve and Machens, 2016; Duarte and Morrison, 2014; Rubin et al., 2017).

As demonstrated in the previous section (Figure 4), the different sources of heterogeneity significantly influence the circuit’s responsiveness, partially by modifying how the different neurons and neuronal populations receive and integrate their synaptic inputs. These differences can also be observed in the amplitude and time course of the total excitatory and inhibitory drive onto each neuron, as can be seen in Figure 5. The results shown are a compound effect of the kinetics of the specific receptors involved and the post-synaptic currents they elicit, the physiological properties of the different neuronal classes as well as the density of specific connections.

**Figure 5.**
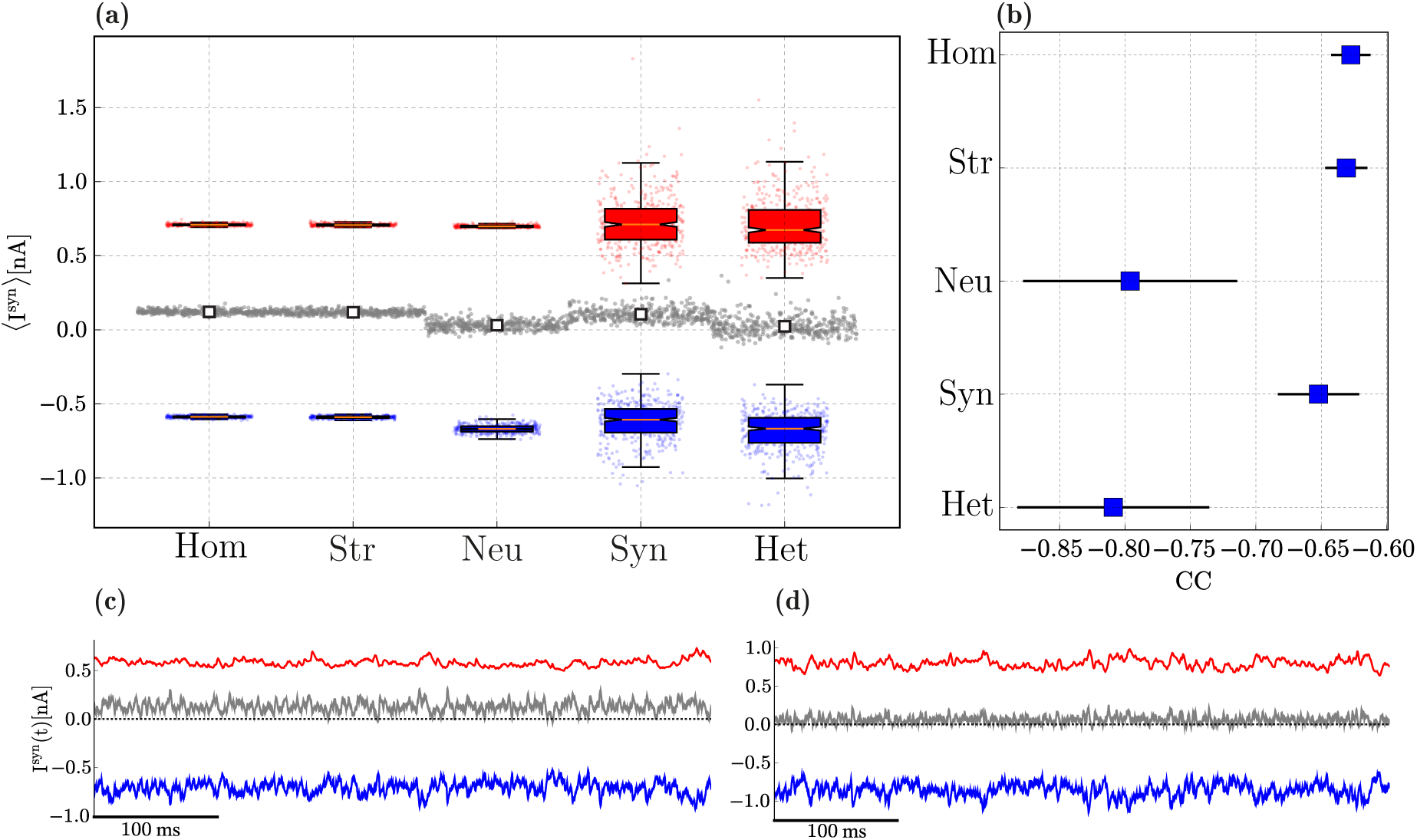
Balance of excitation and inhibition in excitatory neurons driven by background, Poissonian input at *ν_in_* = 10 spikes/s. **(a)**: Distribution of mean amplitudes of excitatory (blue) and inhibitory (red) synaptic currents onto E neurons in the different conditions, as well as the absolute difference between them (grey). For each condition, we show the single data points, where the currents onto a given neuron are summarized as a set of 3 points (〈I^exc^〉 in blue, 〈I^inh^〉 in red and |〈I^inh^) — 〈I^exc^〉| in grey). Overlaid on top of these data points are the distributions across all the neurons, summarized as box-plots: the box represents the first and third quartiles (IQR); the median is marked in red; the whiskers are placed at 1.5 IQR and the outliers can be seen in the underlying data points. The white markers in the middle display the mean difference of synaptic amplitudes across all neurons, for each condition. **(b)**: zero-lag correlation coefficient between excitatory and inhibitory synaptic currents (mean and standard deviation across the E population). **(c, d)**: Examples of the total excitatory and inhibitory synaptic currents received by a randomly chosen E neuron in the homogeneous and fully heterogeneous conditions, respectively.

In a globally balanced state, the amplitudes of excitatory and inhibitory synaptic currents cancel each other on average. This occurs in our microcircuit model only in the presence of neuronal heterogeneity (Figure 5a). Variability in connectivity structure is indistinguishable from the homogeneous condition, whereas variability in synaptic weights and delays significantly increases the variance in the distribution of post-synaptic current amplitudes, but does not shift the mean. This results in an inhibition-dominated synaptic input, resembling that of the homogeneous condition (see also Figure 5c), despite the substantially different distributions.

Apart from being balanced on average, a condition of “detailed” balance (Vogels et al., 2011; Vogels and Abbott, 2009) is characterized by E and I currents that closely track each other and are strongly anti-correlated (Figure 5b). In the homogeneous circuit, excitatory and inhibitory currents are only weakly anti-correlated (*CC* ≈ —0.63). Synaptic heterogeneity causes a slight improvement, but the most significant contribution to this effect comes from neuronal heterogeneity. In this condition, the mean correlation coefficient reaches *CC* ≈ –0.8, although it also introduces a greater variance than synaptic or structural heterogeneity (see also Figure 5d). Both global and detailed balance thus appear to be emergent properties of heterogeneous microcircuits, primarily due to neuronal diversity, but encompassing also a significant influence of synaptic diversity. As is the case with all results presented so far, the fully heterogeneous circuit retains several key properties of interest, but appears to inherit them from different sources, the most significant of which is neuronal heterogeneity. As before, the effects of structural heterogeneity are mitigated by the very sparse firing of the E population, which render its effects moot and no significant deviation from the homogeneous condition is observed.

## 4 Active processing and computation

In order to induce a functional state, engaging the circuit in active processing, we introduce an additional input signal (see Section 6.4.1). We began by tuning the input amplitudes (of both background input firing rate *v*_in_ and external input current *ρ_u_*) independently, for each condition, in order to approximate the relative ratio of mean firing rates among the different populations (see Supplementary Materials), i.e. we attempt to find a combination of input parameters that allows the mean firing rates to remain within realistic bounds (*v*_E_ ∊ [0.5, 5], *v*_I_1__ ∊ [10, 25], *v*_I_2__ ∊ [3,15], considering the values reported in Avermann et al. 2012; Crochet et al. 2011; Gentet et al. 2010; O’Connor et al. 2010).

We consider the circuit’s responses to this input signal as an *active* state, as opposed to the condition explored in the previous sections, where the circuit was driven solely by background, stochastic input (noise). It is worth noting, however, that the similarities between what we call *quiet* and *active* states and their biological counterparts are limited (see Section 5.1). In the following, we show that despite these limitations, the actively engaged circuit operates in similar dynamic regimes to its biological counterpart and maintains the key statistical features that are most likely to play a significant role in modulating the circuit’s processing capacity.

In this section, we assess the microcircuit’s capacity to compute complex functions of the input signal, as described in Section 6.4.3. Note that we purposefully removed any predetermined structure in the input signal, such that the measurements reflect the properties of the system and not the acquisition of structural information in the input (see Section 6.4.3). If we were to consider naturalistic sensory input as the driving signal, this would not be the case.

### 4.1 Spiking activity in the *active* state

Due to the extremely sparse firing observed in the quiet state, an adequate comparison of spiking statistics is only sensible in the active condition. While no explicit effort was taken to constrain the circuit’s operating point when tuning the input parameters (the focus was purely on average firing rates), all conditions operate on an asynchronous irregular regime (exemplified by the raster plots in Figure 6a). This regime is characterized by low pairwise correlations (CC < 0.03) and high coefficients of variation (CV_ISI_ ≈ 0.9), for all neuron classes, in all conditions. Each condition generates different spiking responses with slight variations in activity statistics. The profiles of the activity statistics for the three neuronal classes are summarized in Figure 6 (c)-(g) for the five different conditions. The complete results, displaying the specific profiles for the different classes and conditions separately can be consulted in the Supplementary Materials.

**Figure 6.**
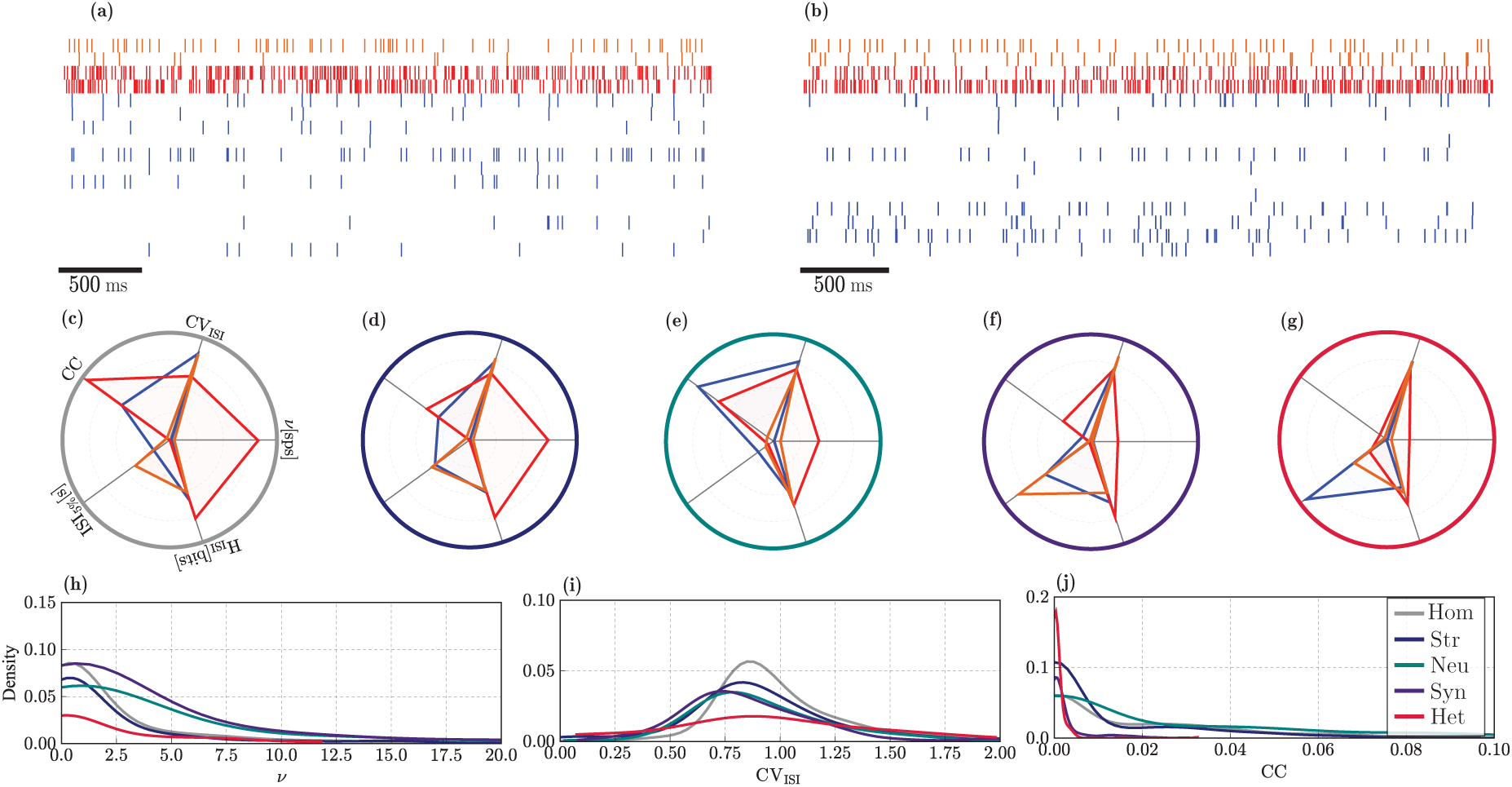
Statistical properties of spiking activity in the active state for the E (blue), I_1_ (red) and I_2_ (orange) populations. **(a, b)**: Example raster plots depicting the activity of a small, randomly chosen subset of neurons, over a recording period of 2.5s in a homogeneous and fully heterogeneous circuit, respectively. **(c)-(g)**: Activity statistics profiles for the different populations (overlaid) in the different conditions (homogeneous (gray), structural (blue), neuronal (green), synaptic (purple), and all (red), respectively). The radial axes represent, as depicted in **(c)**: mean pairwise correlation coefficient (CC), coefficient of variation of the inter-spike intervals (CV_ISI_), mean firing rate (*v*[spikes/s]), entropy (H_ISI_[bits]) and burstiness of the firing patterns (ISI_5%[s]_). The units have been removed for better legibility (see Supplementary Materials). **(h)-(j)**: Distribution of firing rates *v*, CVisi and CC for the E population in the different heterogeneity conditions (see legend in **(j)**).

In the homogeneous condition (Figure 6a and c), I_1_ neurons exhibit the most distinctive profile, with the highest amount of synchrony and the largest firing rates as well as a very bursty and regular firing relative to any other neuronal class and any other condition. Consistent with empirical observations (Crochet and Petersen, 2006; Crochet et al., 2011; Gentet et al., 2010; Poulet and Petersen, 2008), E neurons retain an extremely low firing rate (*v*_E_ ≤ 1 spikes/s), even when stimulated by the extra input current that characterizes the active state. The main difference is that a larger fraction of the population is engaged and actively firing (data not shown). This result demonstrates that sparse firing of E neurons is a stable characteristic of layer 2/3 microcircuits, emerging from the strong and dense inhibition provided primarily by I_1_ neurons that, firing at very high rates, strongly inhibit the E population, regardless of the variations introduced by different sources of heterogeneity and the addition of the extra excitatory input.

In addition, E neurons exhibit the most irregular spiking activity among all neuronal classes. The impact of the various heterogeneity conditions particularly affects the degree of synchronization and burstiness across the different populations. Only the firing rates of I_1_ neurons are significantly modified by heterogeneity, whereas for all other neuron classes, they remain consistently low. Irregularity and randomness in the firing patterns are mostly unaffected as is clear by observing the similarity in the respective axes (H_ISI_ and CV_ISI_ in Figure 6c-g).

The effects of structural heterogeneity (Figure 6d) are only noticeable on the neuronal classes that are directly affected (E and I_1_, see Section 2.3.1 and Table 3); no changes in activity statistics are observable for the I_2_ population (orange profiles in Figure 6c and d). Excitatory neurons fire less synchronously and exhibit a much lower tendency to fire in short spike bursts, compared with the homogeneous condition. On the other hand, I_1_ neurons show a slight decrease in synchrony and firing rate.

Diversity in neuronal parameters (Figure 6e), on the other hand, strongly affects the response properties of I_2_ neurons, slightly increasing their firing rates and correlation coefficient. The most noticeable effect, however is a greatly increased tendency to fire in short bursts (ISI_5%_ ≈ 1*s*) which is the most significant deviation of the standard profile exhibited by this neuronal class in all other conditions (for a more complete comparison, consult the Supplementary Materials). Heterogeneous E neurons have a higher tendency to fire synchronously (albeit still with very low CC), compared to any other condition. As for the I_1_ population, the most significant effect of neuronal heterogeneity is a reduction of the mean firing rate.

Synaptic heterogeneity (Figure 6f) causes a very significant alteration of the firing profile of all neuron classes, particularly E and I_1_, resulting in a very significant decrease in synchronization in all populations, thus having a marked decorrelating effect. It also produces a substantial reduction in the tendency for burst spiking in the E and I_2_ populations. The firing rates of both inhibitory populations are reduced (due to the chosen input parameters, see Supplementary Materials) which, consequently, leads to a slight increase in the E neurons’ firing rates that also fire slightly (but not significantly) more regularly.

Interestingly, the firing profile observed in the fully heterogeneous circuit (Figure 6b and g) exhibits some unique features, different from any of those created by the individual sources of heterogeneity in isolation. Particularly prominent is the complete abolishment of any degree of synchronization in any of the neuronal populations, which show the smallest correlation coefficients of all the cases considered. This effect is likely primarily acquired from synaptic heterogeneity, but goes further than the effect observed there. The firing profile of I_2_ neurons in the fully heterogeneous circuit retains all the features observed in the homogeneous circuit, indicating that the variations introduced by neuronal and synaptic heterogeneity are counteracted by the complex interactions between the different sources of heterogeneity.

Overall, the statistics of population activity clearly demonstrate that the fully heterogeneous circuit is more than the sum of its parts, i.e. the variations introduced by the combination of multiple sources of heterogeneity cannot be fully accounted for by their individual effects and lead to more complex interactions that strongly modulate the circuit’s operating point. In addition, all heterogeneity conditions give rise to similar distributions (Figure 6(h)-(j)), i.e. lognormal distributions of firing rates (argued to be a beneficial feature Buzséki and Mizuseki 2014; Koulakov et al. 2009) and correlation coefficients as well as a Gaussian distribution of CV_ISI_, with mean close to 1. The different conditions simply modulate the parameters of the distributions: synaptic and neuronal heterogeneity (that are responsible for the most significant effects referred so far) broaden the firing rate distributions. Synaptic heterogeneity alone skews the CCs to smaller values, an effect that is stronger and more pronounced in the fully heterogeneous circuit.

### 4.2 Temporal tuning and memory capacity

Layer 2/3 pyramidal neurons receive spatiotemporally structured synaptic inputs from several thousand pre-synaptic sources, through multiple channels conveying different top-down, bottom-up and local information streams that must be integrated in meaningful ways. Whereas the spatial structure is conveyed primarily through non-random projections (conserved topographic maps (Silver and Kastner, 2009; Thivierge and Marcus, 2007) and/or clustered dendritic inputs (Kastellakis et al., 2015; Morita, 2008; Takahashi et al., 2012)), the acquisition and processing of temporal information requires the circuits to integrate their inputs over a broad range of timescales (dynamic range). In this section, we investigate the effect of heterogeneity on the ability of our microcircuit model to retain information over long timescales, enabling it to operate over a broad dynamic range and relatively long timescales.

The baseline, homogeneous circuit already exhibits these properties (Figure 7a), with an average intrinsic time constant (measured as the decay of the membrane potential autocorrelation functions, see Section 6.4) of ≈ 126.75 ms in the quiet state and a relatively broad dynamic range. Thus, even when all the system’s components are homogeneous, neurons appear to operate at relatively long time scales. This can be partially attributed to the fact that each input elicits very small, sub-millivolt responses (Bruno, 2006; Okun and Lampl, 2009) and these neurons are strongly hyperpolarized (Section 3). Therefore, the microcircuit complexity, in combination with the nature of the sub-threshold dynamics, is inherently sufficient to allow it to operate over a large and long temporal range and to rapidly switch to the timescales of its primary driving input (Figure 7a, top).

**Figure 7.**
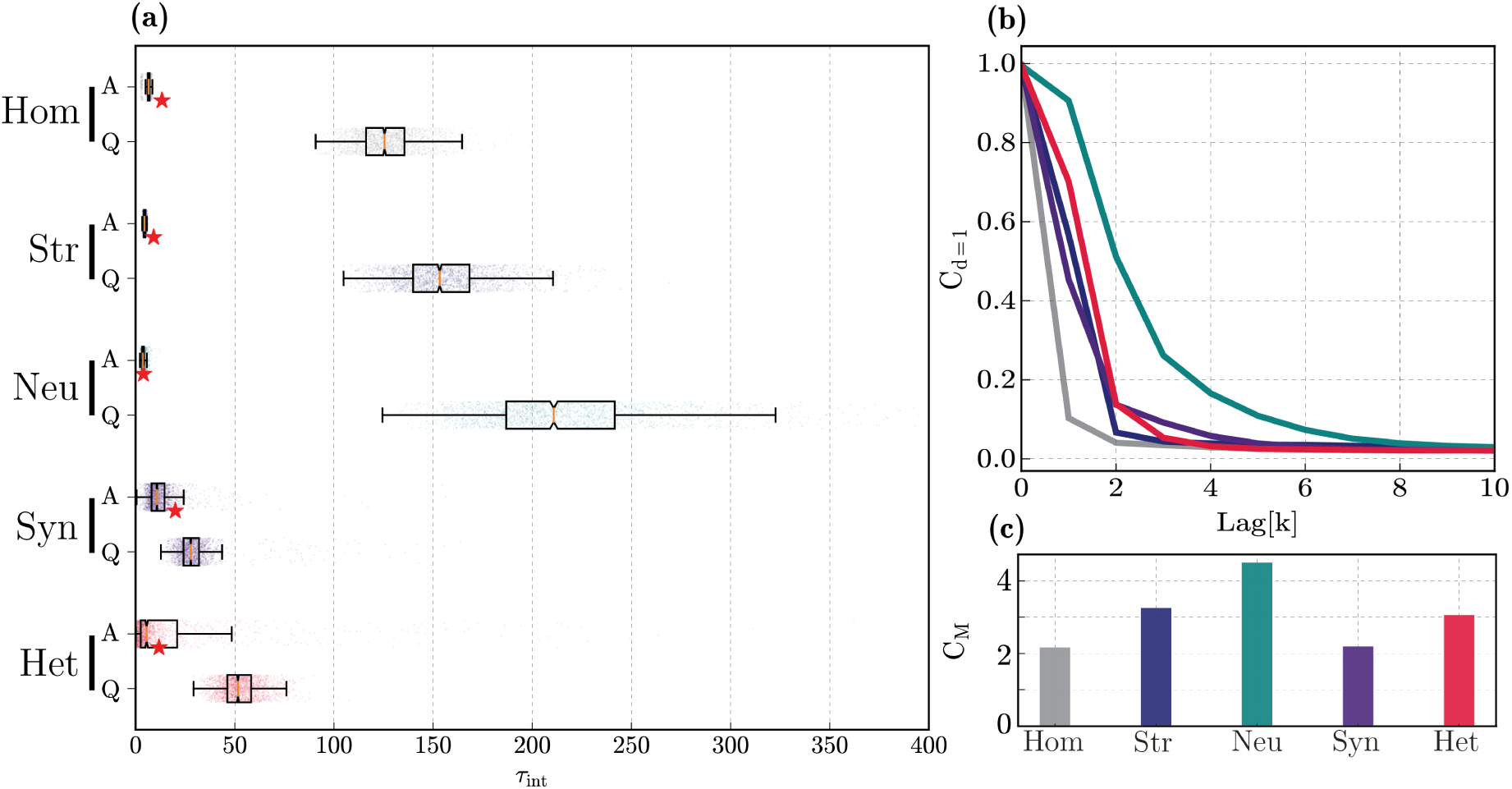
Intrinsic timescales and linear memory capacity. **(a)**: distributions of intrinsic timescales (τ_int_) in the quiet (bottom) and active (top) states, for each of the different conditions. The red stars in the active condition mark the optimal stimulus resolution (Δ*t*) that allows the circuit to perfectly track its input signal. **(b)**: Fading memory functions, determined as the ability to reconstruct the input signal at different delays (k). Colours as in panel (c) below. **(c)**: Total memory capacity, corresponding to the area under the curves in **(b)**.

In order to reliably compute, it is beneficial if the system’s high-dimensional dynamics become transiently ‘enslaved’ by the input (Thalmeier et al., 2016). This would correspond to a switch in the system’s intrinsic timescale to the dominant input time scale, allowing the intrinsic fluctuations to help the circuit track the input dynamics. The observed features of having a broad dynamic range in the quiet state, along with the ability to rapidly switch to the shorter timescale of the driving input during active processing appear to stem primarily from the microcircuit’s composition and dynamics (homogeneous condition). However, they are present to greater extents in the presence of structural and neuronal heterogeneity (Figure 7). The same pattern is true for memory capacity, whereby neuronal heterogeneity causes a large extension of the circuit’s memory range, leading to a slowly fading and relatively long memory store (Figure 7b, c).

The memory range of structurally heterogeneous networks is similar to that of the fully heterogeneous circuit (C_m_ ≈ 3.75 and 3.56, respectively) and both are significantly larger than in the corresponding homogeneous case (C_M_ ≈ 2.67, Figure 7b, c). Thus, despite having a barely noticeable effect on microcircuit dynamics and state transitions, the functional effects of structural heterogeneity appear to be very significant. This is surprising, in light of all the results discussed so far, but consistent with Chaudhuri et al. 2014, who proposed that a heterogeneous network structure can give rise to broad and diverse temporal tuning.

The fact that physiological diversity in single neuron properties (neuronal heterogeneity) extends the dynamic range and memory capacity, is to be expected since it directly decreases redundancy and adds variability to the population responses. However, the magnitude of the effect is very significant, making neuronal heterogeneity the most functionally relevant condition. In the presence of neuronal heterogeneity alone, the circuit becomes much more responsive, with broader temporal tuning and memory capacity (Figure 7) and capable of achieving a significant performance in reconstructing the input signal, even when it varies at short timescales (peak reconstruction performance is achieved with Δ*t* ≈ 3.9 ms versus Δ*t* ≈ 13.3ms in the homogeneous circuit, see Supplementary Materials).

Counter-intuitively, it appears that synaptic heterogeneity makes the circuit ‘sluggish’, incapable of tracking fast fluctuations in the input signal (Δ*t* ≈ 20 ms, see markers in Figure 7a and Supplementary Materials) and endowed with a very short memory capacity (C_M_ ≈ 2.7) that is inferior to that of the homogeneous circuit. Accordingly, diversity in synaptic components enforces a very narrow temporal tuning, skewed towards short timescales in the quiet state (mean *τ*_int_ ≈ 30 ms) and reduces the circuit’s ability to acquire the input timescale (Figure 7a).

These results demonstrate a clear relation between the system’s responsiveness to temporal fluctuations in the input signal, the intrinsic timescales that characterize the neurons’ activity and the circuit’s memory capacity. There appears to be a ‘push-and-pull’ phenomenon caused by the interactions of neuronal and synaptic heterogeneity, whereby the first significantly boosts the circuit’s dynamic range and memory capacity, whereas the second pulls it back to values worse than the homogeneous condition.

#### 4.2.1 Receptor composition

It has been suggested (Duarte et al., 2017a, and references therein) that temporal tuning and memory properties in cortical microcircuits may stem from their biochemical default organization, whose primary consequences are reflected in the regional synaptic composition. In turn, these regional synaptic variations may reflect evolutionary constraints that prime specific circuits to operate on specific timescales, depending on their position in the global processing hierarchy. We note that the main sources we use for synaptic data were recorded in primary sensory cortices. Therefore, the range of synaptic responses could reflect an evolutionary tuning to short timescales, thus reducing its memory capacity. This would partially explain the ‘pull’ effect referred in the previous section.

Accordingly, we hypothesize that different relative concentrations of receptors, such as may be present in other cortical areas, will lead to different intrinsic timescales and memory capacities. To investigate this, we therefore manipulate the ratio of peak conductances in excitatory and inhibitory synapses, globally rescaling *g̅*(NMDA) by *α*_E_ and *g̅*(GABA_B_) by *α* _I_, thus changing the balance of synaptic time constants in the circuit and skewing them to slower excitatory and/or inhibitory transmission by overpowering the relative contributions of NMDA and GABA_B_ receptors (the normal baseline condition corresponds to *α* _E_ = *α* _I_ = 1). The results for memory capacity of systematically emulating the different relative receptor concentrations is shown in Figure 8. Both the fully heterogeneous circuit (Figure 8a) and a circuit with only neuronal heterogeneity (Figure 8b) show a clear dependence of memory capacity on this relative balance. The fully heterogeneous circuit, which has the highest degree of biological verisimilitude of all the examined conditions, shows a significant and consistent effect of *α* _E_, meaning that the balance of excitatory synaptic time constants is particularly important and can dramatically modify the circuit’s memory capacity. Interestingly, the effect is different from what one would expect, as having a larger NMDA component (longer excitatory timescales) essentially halves the memory capacity. This is due to the de-stabilization of the circuit’s operating point (data not shown). Memory is maximized for circuits with a smaller contribution of slow excitatory transmission (*α* _E_ = 1), but a larger contribution of slow inhibition (*α* _I_ =4), see black marker in Figure 8a. Nevertheless, and despite this clear dependence, the memory gains are modest, i.e. the maximum capacity obtained (C_M_ ≈ 4.08) is not significantly larger than the baseline condition (*α* _E_ = *α* _I_ = 1, C_M_ ≈ 3.56) condition, see white marker in Figure 8a.

**Figure 8.**
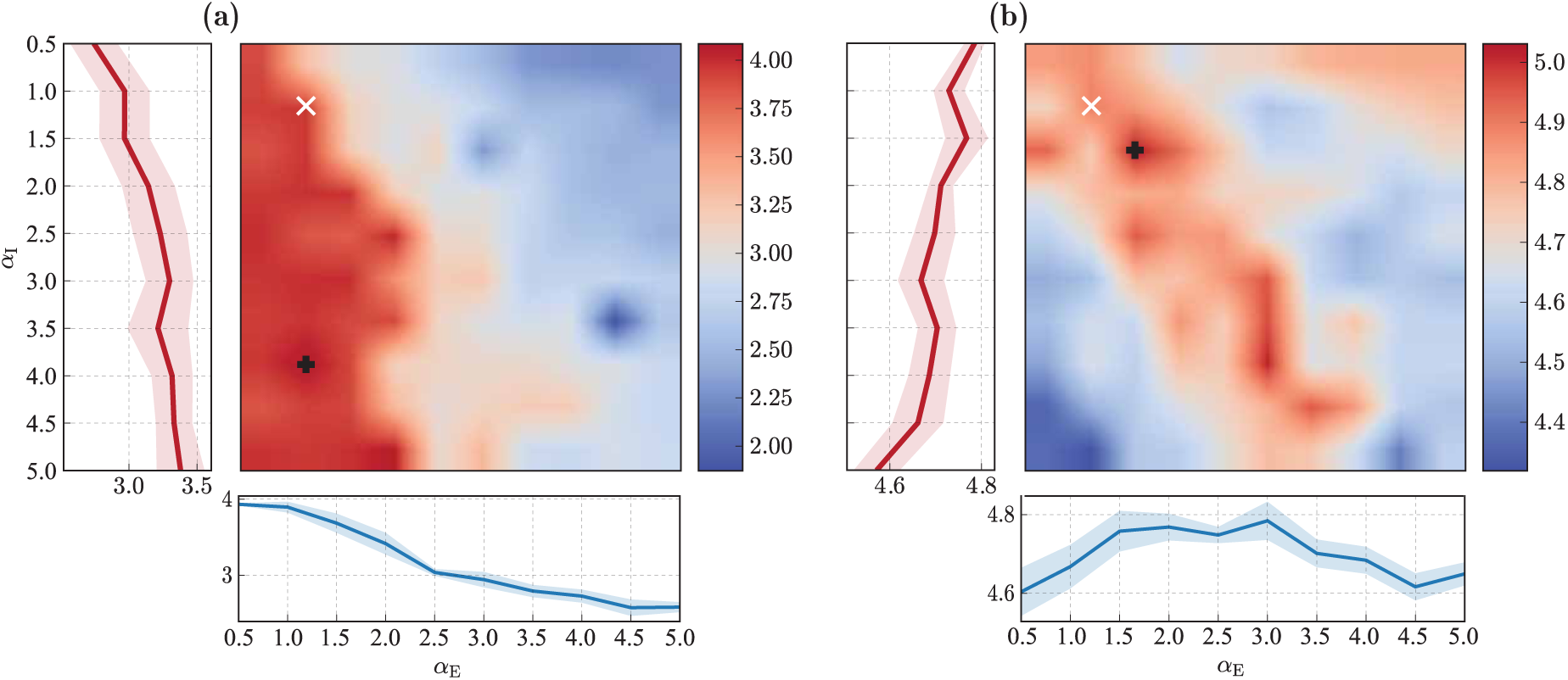
Impact of receptor ratios on memory capacity for the fully heterogeneous circuit. **(a)** and in a circuit with only neuronal heterogeneity **(b)**. Peak conductances of NMDA (horizontal axis) and GABA_B_ (vertical axis) are rescaled for all synapses. Main panels indicate the memory capacity of the circuit (see 7c)’ marginal plots show mean and standard deviation of the memory capacity as a function of the corresponding scaling variable. The markers highlight the default receptor composition (white) and the point where memory capacity is maximized (black). Note the different scales.

For the already high-capacity condition (neuronal heterogeneity, Figure 8b), the combination of synaptic timescales that maximizes memory capacity occurs when both slow excitatory and inhibitory components are slightly larger than the baseline (*α* _E_ = *α* _I_ = 1.5), reaching a maximum value of C_M_ ≈ 5.03. However, the default receptor composition (*α* _E_ = *α* _I_ = 1), already achieves close to maximum capacity (with C_M_ ≈ 5.00). Nevertheless, the dependence on these scaling variables is less straightforward in a circuit with only neuronal heterogeneity, resulting from the effects of both slow inhibitory and excitatory synapses and exhibiting an optimum range where memory capacity is maximized. Although these results confirm that there is a complex dependence of memory capacity and the characteristics of receptor kinetics, the magnitude of the variations are not very pronounced suggesting that other, related processes (short-term synaptic plasticity, for example) contribute to the modulation of memory capacity.

### 4.3 Processing capacity

To determine the microcircuit’s suitability for online processing with fading memory, we adopt the notion of information processing capacity introduced in Dambre et al. (2012), which allows us to quantify the system’s ability to employ different modes of information processing and, by combining them, determine the total computational capacity of the circuit (see Section 6.4.3). By this definition, the memory capacity discussed in the previous sections corresponds to the capacity to reconstruct the set of *k* different linear functions (corresponding to degree 1 Legendre polynomials, see Section 6.4.3) of the input *u*, each corresponding to a specific time lag, see also Jaeger (2002). As such, it corresponds to the fraction of the total capacity associated with linear functions (since no products are involved) and measures the circuit’s linear processing capacity (*d* = 1 in Figure 9). Accordingly, degrees *d* ≥ 2 correspond to larger and increasingly complex sets of non-linear functions (products of Legendre polynomials) and thus require increasingly more sophisticated computational capabilities.

**Figure 9.**
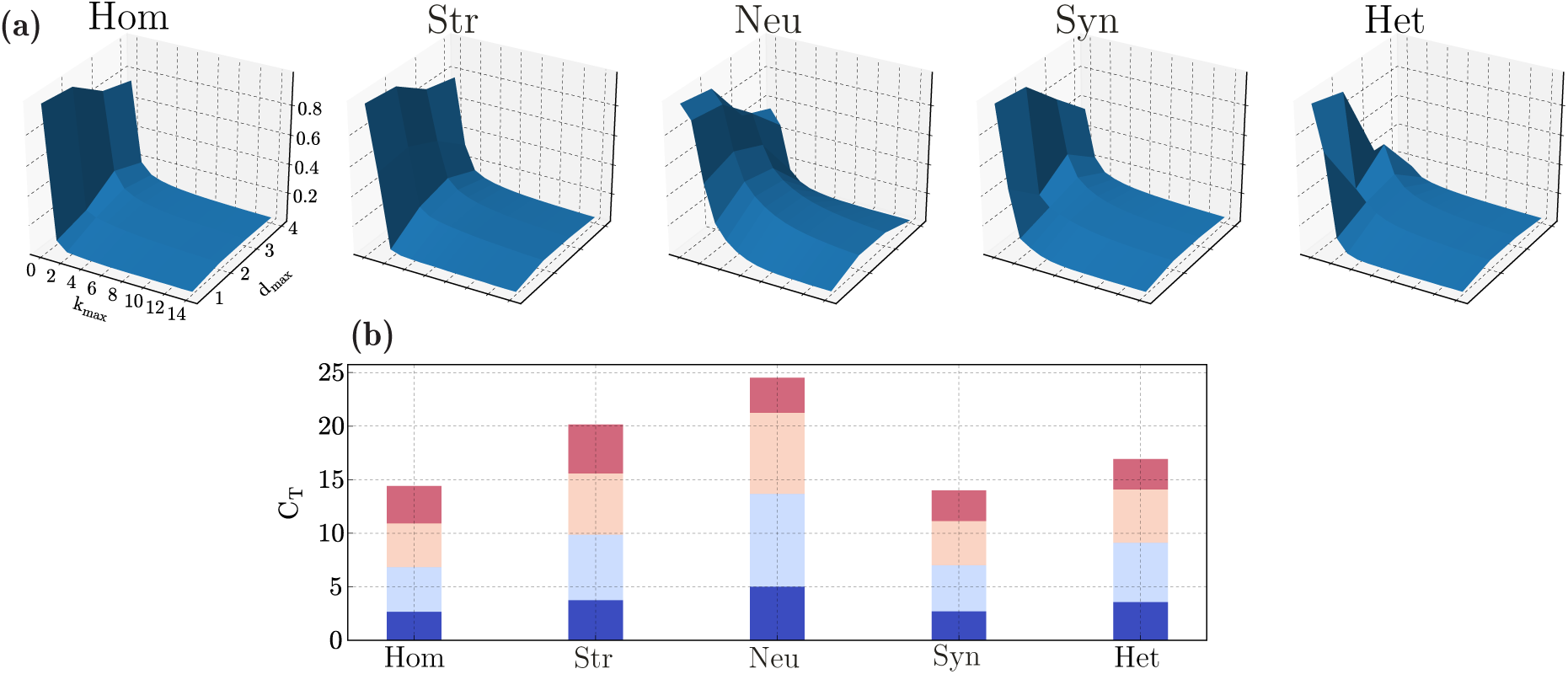
Computational capacity in the different heterogeneity conditions. **(a)** Normalized capacity space, i.e. ability to reconstruct functions of *u* at different maximum delays (k_max_, memory) and degrees (d_max_, complexity/nonlinearity). **(b)** Total processing capacity, expressed as the sum of all capacities for a given degree (the incremental color code in each bar correspond to the maximum degree for each segment, varying from 1 to 4).

We can thus distinguish between computational complexity / non-linearity (specified by the maximum degree of the basis functions used, *d*_max_) and memory (maximum delay taken into consideration, *k*_max_). By evaluating a very large set of functions of *u*, we can approximate the circuit’s capacity space as a probability distribution over the space of basis functions.

These results are shown in Figure 9a. In line with the previous results (Section 4.2), heterogeneity in neuronal parameters has the most significant effect, greatly extending the space of computable functions, both linear and nonlinear. By allowing the circuit to retain contextual information for longer (extended memory range), these circuits have a high capacity even for relatively large delays, as demonstrated by the slowly decaying memory curves in Figure 9a. As a consequence, the total capacity of microcircuits with heterogeneous neurons is the largest among all the conditions (Figure 9b).

Despite its very modest effects on population activity, structural heterogeneity has very interesting consequences on the microcircuit’s processing capacity, particularly in the ability to compute very complex nonlinear functions. Although the memory functions decay abruptly (almost as abruptly as in the homogeneous condition, Figure 9a), the circuits achieve a larger capacity for more complex functions (Figure 9b) and the total capacity at *d* = 4 (largest degree evaluated) is the largest among all conditions tested, as can be seen by comparing the top crimson bars in Figure 9b.

Also in line with the results for *d* =1 (linear memory capacity), synaptic heterogeneity has an unexpected deleterious effect on processing capacity, reducing it to values smaller than the homogeneous case (C_T_ ≈ 13.9 versus C_T_ ≈ 14.4 for the homogeneous condition, Figure 9b). Consequently, the beneficial effects introduced by both structural and neuronal heterogeneity are counteracted by the negative effect of synaptic heterogeneity, as previously observed in Section 4.2. These cancelling effects result in the fully heterogeneous circuit having a total capacity that is only modestly superior to the homogeneous case (C_T_ ≈ 16.9).

Under idealized conditions, the total capacity is bound by the number of linearly independent state variables of the dynamical system, which, in the limit of *T* → ∞, equals N (the number of neurons), in systems that perfectly obey the fading memory property and whose neurons’ activity is linearly independent (for proofs, see Dambre et al. 2012). In that respect, the values we have obtained for the total capacity are very modest and close to only 1% of the theoretical limit. On the one hand, this is due to methodological limitations (we could only investigate a short range of d_max_), and on the other, suggests that there may be important aspects that were neglected in this study that significantly boost the total capacity and, at least partially explain the difference (see Section 5.3).

Nevertheless, the results demonstrate a consistent pattern to that observed in Section 4.2, i.e. the functional consequences of the different sources of heterogeneity are consistent for both the ability to compute linear and non-linear functions with fading memory, as circuits with the largest linear memory capacity are also the ones with the largest non-linear processing capacity (namely neuronal heterogeneity). However, these results indicate that neuronal heterogeneity has its main effect on memory (k_max_), greatly extending the capacity to compute functions of *u*_–*k*_ for larger values of *k* (Figure 9a) while structural heterogeneity (that has the second largest functionally beneficial effects) significantly boosts the ability to compute more complex functions, with a main effect on d_max_.

## 5 Discussion

Heterogeneity and diversity in cellular, biochemical and physiological properties seen within and across cortical regions and layers exerts a significant influence on population dynamics. Although often disregarded by the reduced models commonly used in computational neuroscience, these features of the neural tissue may well be partially responsible for the high computational proficiency and functional properties of these systems. In order to understand the functional relevance of the different ‘building blocks’ and their inherent complexity and diversity, it is important to start from relatively simple formalisms and gradually account for the biological complexity while maintaining coherence with the relevant empirical observations at multiple levels. The present study proposes a data-driven modelling approach as an exploratory strategy to systematically uncover the computational benefits of different microcircuit features in an attempt to elucidate and quantify the biophysical substrates of neural computation.

We have focused on the composition of layer 2/3 cortical microcircuits (Section 2), since their highly recurrent connectivity (Feldmeyer et al., 2006; Lewis and Gonzalez-Burgos, 2000) and sparse, asynchronous activity (Avermann et al., 2012; Lewis and Gonzalez-Burgos, 2000; Neske et al., 2015; O’Connor et al., 2010; Petersen and Crochet, 2013; Waters and Helmchen, 2006) are ideally suited to study the nature of sparse distributed processing in cortical microcircuit models. The small extent of the neuritic processes (in comparison with deeper layers), makes it more reasonable to assume that the potential role of dendritic compartmentalization (Branco and Hausser, 2010; Morita, 2008) and other effects caused by the detailed neuronal morphology (Kubota et al., 2015; Spruston, 2008) are negligible, allowing us to use simple point-neuron models with limited loss - which would not be the case if we accounted for the deeper layers. Additionally, the input/output relations and unique position of layer 2/3 in a cortical column suggests a particularly prominent computational role as it must integrate and process multiple streams of information in meaningful ways.

In an attempt to disentangle the role played by heterogeneity in different components of the system, we tentatively partitioned it into neuronal, structural and synaptic components (see Section 1.1, Section 2 and Supplementary Materials). These different sources of heterogeneity differentially influence the characteristics of population responses: from introducing variability in how different neurons and neuronal classes respond to and integrate their synaptic inputs, to variations in the magnitude and distribution of those inputs, among other effects. These, often subtle, differences have complex effects at the population level and strongly condition the system’s operating point. The fully heterogeneous circuit provides the closest approximation to the biophysical reality, exhibiting important commonalities, and appears to inherit different features from different sources of heterogeneity. Naturally, given the simplifications required, our conclusions on the effects of the various heterogeneities are primarily qualitative. However, the extent to which the model responses differ from the empirical observations can also be informative about the potential impact of microcircuit features and processes that were not explicitly considered or were overlooked or oversimplified, as we will discuss below.

### 5.1 Cortical states

The state of any given cortical microcircuit, both in terms of macroscopic spiking statistics and, particularly, membrane potential dynamics can differ dramatically between behavioural states (Crochet and Petersen, 2006; Crochet et al., 2011; Gentet et al., 2010; Poulet and Petersen, 2008) given that they require different levels of active ‘engagement’. The three neuronal classes behave in very specific ways, with specialized response features providing differential contributions to the different circuit states. These neuron-class-specific contributions play an important role in the observed dynamics, providing a potential mechanism to support state modulations (Pachitariu et al., 2016; Poulet, 2014).

Spontaneous cortical activity during states of *quiet wakefulness* (a quiescent state in which the animal is awake but the circuit is not directly engaged in active processing), is commonly characterized by short-lasting, large amplitude depolarizations (DeWeese and Zador, 2006; Petersen et al., 2003; Poulet and Petersen, 2008) that reflect the presence of strongly synchronized excitatory inputs and resemble the dynamics observed under light anaesthesia (Crochet et al., 2011; Neske et al., 2015; Okun and Lampl, 2009; Poulet and Petersen, 2008). Naturally, driven by a homogeneous Poisson process (Section 6.4.1), the system does not exhibit such behaviour (see Section 3), which indicates that such effects are partially inherited by the spatiotemporal structure of the background input (Poulet et al., 2012), which in turn may reflect the structure of the sensory input (Luczak et al., 2007). Additionally, or alternatively, this may be a consequence of propagating waves of excitation (Petersen et al., 2003) which are likely related to spatial connectivity features that were not taken into consideration in this study (see Section 5.3).

Nevertheless, our *quiet* state, where the circuit is driven by background noise, highlights relevant features of population activity and their relations among different neuronal classes, emerging from the effects of the different sources of heterogeneity (Section 3). The most prominent feature is the extremely sparse firing of E neurons (Figure 4c), which appears to stem directly from the circuit’s composition (homogeneous condition) and is a robust and replicable effect emerging as a direct consequence of dense, strong and fast inhibition. While structural heterogeneity has no measurable effects, synaptic heterogeneity makes the E population more responsive and places some of these neurons closer to their firing thresholds (Figure 4d). Neuronal heterogeneity, on the other hand, leads to more strongly hyperpolarized E and I_1_ populations, compared to all other conditions. This has the positive effect of shifting the distribution of membrane potential in the I_1_ population to a range that overlaps with the empirical values in Gentet et al. (2010)). However, E neurons become excessively hyperpolarized and their membrane potentials are kept farther from threshold and farther from the corresponding experimental value (to which all other conditions provide a better match). Despite these differences, neuronal heterogeneity is responsible for placing all three neuronal populations operating within the range of values reported in the literature (Figure 4d) and, simultaneously maintaining the correct ratio of effective membrane time constants, thus significantly modulating how the circuit responds to synaptic input (Figure 4e).

Neocortical pyramidal neurons (particularly in layer 2/3) fire very sparsely and are never driven to saturation, despite a large and constant synaptic bombardment. For this to occur, excitatory and inhibitory input currents onto each neuron must be carefully balanced such that, on average, they cancel each other, allowing the net mean input to be small and the output rates moderate (Shadlen and Newsome, 1998, 1994). Co-active and balanced excitation and inhibition thus stabilizes and shapes the circuit’s activity and must be actively maintained to allow the networks to operate in stable regimes (Duarte and Morrison, 2014; Ernst and Pawelzik, 2011; Vogels et al., 2005). In a balanced configuration, local cortical dynamics is mostly driven by fluctuations of synaptic inputs, which provides a mechanism to explain a fundamental feature of cortical dynamics: variability. Importantly, it also plays a critical role in active processing and computation, with the most clear experimental evidence coming form the development of input selectivity in visual and auditory cortices (see e.g. Dorrn et al. 2010; Marino et al. 2005; Okun and Lampl 2009 and references therein). Any disruption of balance can have dramatic consequences, leading to epileptogenic activity (Dehghani et al., 2016; Dichter and Ayala, 1987), with the associated impairment in information processing and behaviour (Yizhar et al., 2011). In Section 3.1, we demonstrate that such balance condition is an emergent property from circuits with heterogeneous neurons, without the need for changing any of the system’s parameters.

Despite limitations in our study, discussed below, our results highlight the importance of developing new theories of cortical function and dynamics based on the complex interactions of multiple neuronal sub-populations, as different neuronal classes have a non-negligible differential contribution to the circuit’s dynamics. Additionally, the prominent functional role of structural and neuronal heterogeneity suggest that they are part of a critical minimum necessary to account for computation in cortical microcircuit models.

### 5.2 Heterogeneity and information processing

At any given point in time, the state of the circuit reflects a nonlinear combination of the current and past inputs, mediated by complex recurrent interactions. The state of each neuron is thus a nonlinear, fading memory function of the input (the characteristics of which are determined by the circuit’s specificities and input encoding) and the population state a set of *N* basis functions that can be linearly combined to approximate arbitrary nonlinear functions with fading memory. In that sense, these circuits are endowed with universal computing power on time invariant functions (Enel et al., 2016; Maass et al., 2002, 2004; Thalmeier et al., 2016). This is where complexity and heterogeneity play a particularly prominent role, as they can greatly extend the space of computable functions by diversifying population responses and, consequently, the richness of the circuit’s high-dimensional state-space.

With specific functions in mind, circuits can be "designed” to perform certain computations by explicitly solving the credit-assignment problem, i.e. determining how each neuron ought to contribute to the computation (Abbott et al., 2016) in order to achieve the desired outcome. This is typically achieved by constraining the microcircuit connectivity (Memmesheimer and Timme, 2006a,b) and/or by postulating and building-in specific functionality (e.g. efficient / predictive coding; Boerlin et al. 2013; Deneve and Machens 2016). The great majority of these approaches, however, assumes idealized conditions and neglects the complexities of real biophysics (but see, e.g. Schwemmer et al. 2014), which limits their scope and generalizability.

Since we were not interested in specific functions, but in universal computational properties, instead of "designing” functional microcircuits or assuming specific computations, we sought to mimic fundamental design principles of the real neocortical microcircuitry and systematically evaluate how they affect the circuit’s computational capabilities. While this exploratory approach has its limitations, we were able to partially disentangle the computational role of complexity and heterogeneity in the microcircuit’s building blocks and pinpoint potential sources of functional specialization. Globally, the functional analysis on the computational benefits of the different sources of heterogeneity revealed the same effect: neuronal diversity, on its own, significantly boosts linear and nonlinear processing capacity and memory (see Section 4.2 and Section 4.3) and dramatically increases its dynamic range and sensitivity (Section 4.2). Surprisingly, and even though its effects on population activity were barely noticeable, structural heterogeneity has the second largest computational effect, particularly boosting the ability to compute highly nonlinear functions (capacity at *d* = 4 was much larger than any other condition, see Section 4.3). Surprisingly, the benefits introduced by neuronal and structural heterogeneity are not reflected in the fully heterogeneous circuit, as synaptic heterogeneity prevents this from happening. It would expectable and desirable that the computational benefits would combine in a way that could dramatically increase the total capacity of the most realistic condition. As discussed below, these synaptic effects likely reflect important limitations in our ability to capture their real influence in the biological system.

The complete dominance of neuronal heterogeneity, not only in terms of computational proficiency, but any other property analysed, suggests that it should be taken into consideration in studies focused on reduced, simple models of cortical microcircuits as its effects appear to underlie a variety of important phenomena.

### 5.3 Limitations and future work

As discussed in Section 1, the myriad of complex features that characterize any given cortical microcircuit entail considerable constraints on our ability to define an adequate level of descriptive complexity, and discern which elements are functionally meaningful and partially responsible for the system’s computational proficiency. Despite providing a significant step towards biological verisimilitude, away from the idealized conditions typically assumed in computational and theoretical studies of cortical dynamics, our results demonstrate important limitations that ought to be addressed in future work.

Even though we consider three different neuronal populations, including two separate inhibitory classes, further sub-divisions have been reported in neocortical layer 2/3 populations, both for glutametergic (Zaitsev et al., 2012) and, in particular, for GABAergic neurons (Ascoli et al., 2008; Helmstaedter et al., 2009; Jiang et al., 2016; Neske et al., 2015; Petersen and Crochet, 2013). It is possible that these reflect regional specializations particularly prominent in specific cortical areas (such as the prefrontal cortical regions; Harris and Shepherd 2015; Shepard and Grillner 2010; Zaitsev et al. 2012) or that they represent separate instances of broader classes and can, for simplicity, be grouped together. We conclude that it is important to understand the role of multiple interacting populations (e.g. Lagzi and Rotter 2015), particularly including inhibitory sub-types and their different physiological properties and interactions, given their clearly distinct contributions.

As in every modelling study, one needs to consider the trade-off between realism and computational cost (Herz et al., 2006). Parameterized correctly, our choice of neuron model proved to be sufficient for the purposes of this study and allowed us to account for the most important physiological characteristics of the different neuronal classes and their relations (Section 2.1). Such simplifying assumptions, however, are bound to miss relevant structural and functional features, particularly when it comes to specialization of inhibitory neurons and synapses (Fino et al., 2013; Gupta, 2000; Isaacson and Scanziani, 2011; Pi et al., 2013; Wilson et al., 2012), the effects of dendritic nonlinearities (Branco and Hausser, 2010; Kubota et al., 2015; Morita, 2008; Spruston, 2008) and intrinsic adaptation processes (Pozzorini et al., 2013), to name a few. It is also important, in future work along this direction, to consider the intricate relations between model parameters, i.e. explicitly include not only the empirical variability but also the covariance across multiple parameters (as e.g. Harrison et al. 2015).

When it comes to synaptic transmission, we have focused on the specificities of instantaneous response kinetics and its inherent diversity, disregarding any form of synaptic plasticity. However, in our model synaptic heterogeneity was shown to severely constrain the microcircuit’s processing capacity and memory (Section 4.3), counteracting the benefits introduced by neuronal and structural heterogeneity. Additionally, the fact that the total measured capacity is very modest even in the best-performing systems (only about 1% of the theoretical maximum), and considering the computational requirements posed on these systems in ecological conditions, this suggests that there are important aspects of synaptic transmission that we have failed to consider, but contribute significantly to the circuit’s processing capacity. Adaptation and plasticity are likely to be important missing components, due to their critical roles in learning and memory processes (Abbott and Regehr, 2004). It would be important, in future work, to capture, at least, the dynamics of short-term plasticity (Hennig, 2013), which also differ substantially among different synapse types (Gupta et al., 2000) and is one of the processes which, operating at these timescales can significantly affect the fading memory property. Additionally, long-term plasticity (Holtmaat and Svoboda, 2009; Maffei, 2011; Morrison et al., 2008) and homeostasis (Turrigiano, 1999), shaping circuit dynamics and processing (Duarte and Morrison, 2014) over long timescales (Tetzlaff et al., 2012; Zenke and Gerstner, 2016) lead to cumulative changes that may result in an improved domain-specific processing capacity.

In our study we investigated the behaviour of our microcircuit model in two dynamic regimes, which we associated with the biological *quiet* and *active* states. However, the stimulation applied to bring the circuits into the active state was not biologically realistic, as we purposefully removed any spatiotemporal structure in order to measure the computational properties of the system and not the acquisition of structural information present in the input signal. Thus, the degree to which we are able to account for and explicitly compare empirical observations with the model is restricted and only qualitative. Moreover, whilst measuring the capacity of the network, we significantly under-sampled the space, as the results clearly demonstrate (Figure 9a). A more complete set of basis functions would lead the capacity along both axes to decay to 0: as the complexity and memory requirements increase, the capacity to compute these functions decrease to negligible values in all systems. While accounting for delays of up to k_max_ = 100 allowed us to capture this effect (since the memory range in all conditions is inferior to that), we failed to account for a sufficiently large d_max_. The primary reason for this was computational cost, as our current implementation is extremely time-consuming (see Supplementary Materials). As a consequence, the capacity space is sub-normalized, incomplete and underestimated, due to the relatively small number of basis functions tested. Additionally, the limited sample size (*T* = 10^5^) may bias the individual results.

### 5.4 Data-driven computational neuroscience

Collecting, validating and organizing experimental data relevant for these type of studies is still a monumental challenge. Manual annotation and parameter extraction are cumbersome, error-prone strategies and only feasible on well-constrained systems and well-defined problems. The creation and active curation of stable and reliable large-scale databases (of which good examples exist: Allen Brain Atlas^2^ (Sunkin et al., 2013), NeuroMorpho^3^ (Ascoli et al., 2007), NeuroElectro^4^ (Tripathy et al., 2014), NMC^5^ (Ramaswamy et al., 2015), ICGenealogy^6^ (Podlaski et al., 2016), to name a few), along with standard and widely accepted registration and sharing practices (Peng, 2011; Zehl et al., 2016) are increasingly a priority in a community-driven effort to better constrain neuroscience models and integrate knowledge from multiple disciplines. In addition, automated parameter extraction, estimation as well as model fitting and comparison is strictly and increasingly necessary for studies in this direction. These are complex challenges as they must meet the requirements of an ever-changing scientific field that, consequently, doesn’t lend itself easily to standardization.

As carefully curated and organized datasets become increasingly available, it will become possible in the near future to apply increasingly realistic constrains and comparatively study the properties of realistic microcircuits, built to model specific cortical regions, their local input-output relations and systematically assess the impact of different modifications in the circuit’s function as an information processing, general-purpose device.

Apart from the considerable efforts to explicitly include and account for experimental data to constrain the microcircuit models and make use of publicly available datasets, we additionally emphasize the importance of ensuring transparency, openness and reproducibility. To this end, the complete materials for this study are publicly available through the Open Science Collaboration (2015) (see Supplementary Materials for details). Our efforts in that direction are a mere example and proof-of-concept, but demonstrate that the field is mature enough to embrace these practices, which should become a standard in computational neuroscience. Given the complexity of these studies, the ability to reproduce and verify are paramount not only to impose the scientific ‘golden standards’, but to extend and build upon existing work.

## 6 Materials and methods

### 6.1 Neuronal dynamics

In all systems analysed, the neuronal dynamics is modelled using a common, simplified adaptive leaky integrate-and-fire scheme (Koch, 2004), where the total current flow across the membrane of neuron *i* is governed by:

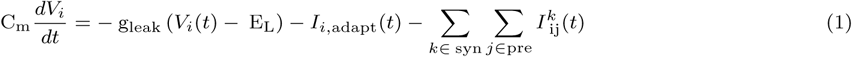

The spike times of neuron *i* are defined as the set *F*(*i*) = {*t_f_* | *V_i_*(*t_f_*) ≥ *V_thresh_*}. At these times, the membrane potential is reset to the constant value *V*(*t*) = *V*_reset_ for all times *t* ∊ (*t_f_*, *t_f_* + *t_refr_*], after which integration is resumed as above. *I*(*t*) is the total synaptic current generated by inputs from all pre-synaptic neurons *j* ∊ *pre* mediated by synapse type *k* ∊ *syn* (see Section 6.2).

To provide greater control over neuronal excitability properties and a more realistic account of cortical neuronal dynamics, we model intrinsic adaptation processes as proposed by Gerstner et al. (2014):

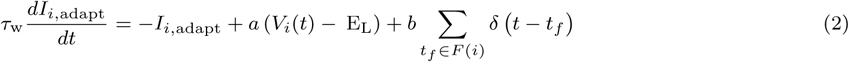

where the parameters *a* and *b* determine the relative contribution of sub-threshold and spike-triggered adaptation processes, respectively.

### 6.2 Synaptic dynamics

The synaptic current 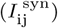 elicited by each spike from presynaptic neuron *j* is determined by the conductivity (*G*^rec^) of the corresponding, responsive receptors (each synapse type being composed of a pre-determined set of receptors, see below):

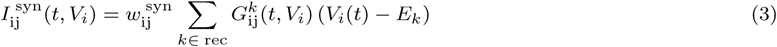

The amplitude of post-synaptic currents is rescaled by the dimensionless weight parameter 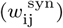, specific to each connection type and whose value was chosen, such that the PSP amplitudes matched the data reported in (Avermann et al., 2012; Lefort et al., 2009) (see Table 3). The synaptic conductivity 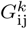 in Equation 3 models the response of receptor *k* to spike arrival from pre-synaptic neuron *j* with a total conduction delay of 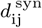:

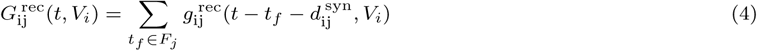

The conductance transient elicited by a single pre-synaptic event on a single post-synaptic receptor is then modelled as (Gerstner and Kistler, 2002; Gerstner et al., 2014; Hoffmann et al., 2015; McCormick et al., 1993):

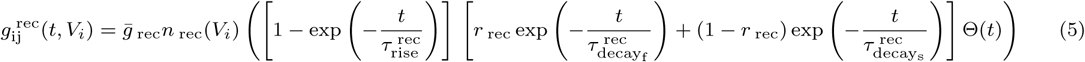

where *g̅*_rec_ is the peak conductance of the corresponding receptor, *n* _rec_(*V*) is a voltage-gating function assuming a constant value of 1 for all receptor types, except NMDA, in which case (Jahr and Stevens, 1990):

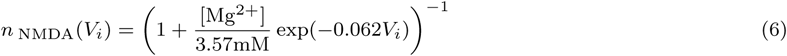

This gating function is thus used to model the voltage-dependent magnesium block at NMDA receptors (see below). For simplicity, we assume a fixed [Mg^2+^] = 1mM. The remaining parameters in Equation 5 correspond to the receptors’ characteristic time constants, namely the rise, fast and slow decay times, as well as the relative balance between fast and slow decay (*r* _rec_). In order to account for the differential receptor composition and expression across different neuronal classes, all these parameters are specific for each receptor, synapse and neuron type.

### 6.3 Generating structural heterogeneity

Consider the sparse adjacency matrix *A* ^syn^, specifying the anatomical connectivity between all neurons in source population *pre* and target population *post* (with *pre*, *post* ∊ {E, I_1_, I_2_}). The indices i, j of the nonzero entries in *A* ^syn^ are independently drawn from normalized, truncated exponential distributions, with probability:

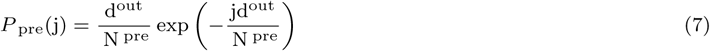

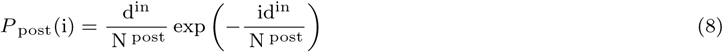

for pre- and postsynaptic neuron indices, respectively. N ^pre/ post^ is the total number of pre-/postsynaptic neurons and d^out/in^ are the parameters used to define the skewness of the out-/in-degree distributions, respectively. Setting d^out/in^ = 0 corresponds to a random, uniform connectivity, whereas values > 0 generate structured in-/out-degree distributions, with a larger variance in the number of connections per neuron.

#### 6.3.1 Weight correlations

For each existing connection, the individual synaptic efficacies 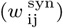 can be equal to a fixed scalar value (homogeneous synaptic condition) or randomly drawn from a lognormal distribution (heterogeneous condition). To introduce weight correlations, we specify additional scaling variables *ζ* for each pre- or postsynaptic neuron (*ζ*_j_, *ζ*_i_), whose values are independently drawn from 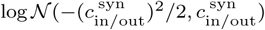. The parameters 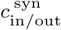 determine the strength of induced correlations, which we fix and set to the values reported in Tomm (2012) and Tomm et al. (2014). The original weight values 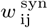 are then re-scaled by *ζ* of the corresponding pre- and postsynaptic partner, i.e.:

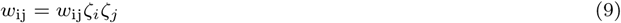

For completeness, and given that lognormal distributions are widely employed throughout this study, it is worth noting that the lognormal probability density function has the following form:

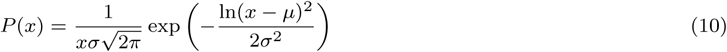

parameterized by scale (*μ*) shape (*σ*) values.

### 6.4 Profiling the microcircuits

To adequately quantify the relevant functional properties of the microcircuits and the impact of the different features analysed, we employ metrics that are system-agnostic, i.e. independent from the specificities of the circuit analysed and, preferably, parameter-free such that the choices of metric parameters do not influence the measured outcome and any results obtained are unbiased and objectively reflect the circuit’s properties. Of particular interest, for the purposes of this study, is the adequate quantification of the characteristics of population dynamics, under active synaptic bombardment as well as the circuit’s capacity for stimulus processing and computation, in order to establish links between the features of population dynamics, the circuit’s composition and complexity and its ability to perform complex computations.

#### 6.4.1 Input specifications

We model cortical background / ongoing activity as an unspecific and stochastic external input driving the circuit, considered to be excitatory, i.e. mediated by glutamatergic synapses, and to consist of independent Poissonian spike trains, at a fixed rate *v_in_* spikes/s. Given the small network size and, in order to compensate for the relatively small numbers of synapses involved, we rescale the input rates by a constant factor, K_in_ = 1000, such that each neuron receives, on average, background input through K_in_ synapses, with each presynaptic source firing at a fixed rate of *v_in_* spikes/s. The postsynaptic neuron’s responsiveness to this background input is then determined by its specific receptor composition (the kinetics of AMPA and NMDA receptors), with synaptic weights and delays equal to any other excitatory synapse onto that neuron, i.e. 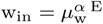 and 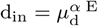, where *α* refers to the postsynaptic neuron class (see Table 3).

In addition, to evaluate the microcircuit’s processing capacity (see Section 6.4.3), we introduce an additional input signal, directly encoded as a piece-wise constant (in intervals of Δ*t* ms), somatic input current (*I_in_*(*t*)), delivered to a randomly chosen sub-set of the E population, comprising 0.25 N _E_ neurons (Figure ??a). We choose this direct encoding strategy in order to ensure that the input signal has a direct influence on the membrane dynamics, making it easier to decode (Eliasmith and Anderson, 1991; van den Broek et al., 2017; Weidel et al., 2016).

#### 6.4.2 Population dynamics

To quantify and characterize the population responses to different input conditions and assess how the different sources of heterogeneity modulate those responses, we look at the statistics of spiking activity as well as the relevant sub-threshold dynamics across the different neuronal populations.

The circuit’s state or operating point is typically determined primarily by the firing rate and the degree of population-wide synchrony and regularity (Tsotsos et al., 2000), as measured by the following statistics:

**Synchrony** - average spike count correlation coefficient (CC), computed pair-wise, for a large number of n^pairs^ = 500 randomly sampled, disjoint, neuronal pairs (see, e.g. (Kumar et al., 2008))
**Regularity** - degree of dispersion of the inter-spike interval (ISI) distribution, as measured by the coefficient of variation (CV_ISI_), averaged across the population. A value of 0 indicates a clock-like, regular firing pattern, whereas a completely irregular, Poisson process has a value of 1. Values larger than 1 are obtained for very bursty firing patterns.
**Burstiness** — degree of burstiness in the firing patterns can be captured by the 5-th percentile of the ISI distribution (ISI5%), averaged across the population. This measure has been successfully used to classify neuronal firing patterns in identified populations (Ruigrok et al., 2011; van Dijck et al., 2013). A low value indicates higher burstiness, for a given rate.
**Randomness** - The entropy of the log-ISI distribution is an important metric that captures the randomness in a spike train (Dorval, 2008). It was recently demonstrated that this metric was one of the key features of cerebellar neurons’ spike trains, allowing their accurate identification (van Dijck et al., 2013). This metric is defined on the probability density of the natural logarithm of the ISIs:

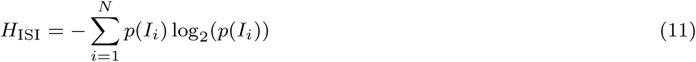 This last metric is added for completeness, and can be seen as an additional measure of regularity. The higher the entropy value, the more irregular the firing pattern. On the sub-threshold level, we assess and summarize the characteristics of membrane potential dynamics, synaptic currents and conductances. We analyse the distributions of mean membrane potentials (〈V_m_〉) and their variances (σ^2^ (V_m_)), as well as the mean excitatory and inhibitory synaptic currents (〈I^syn^〉) onto each neuron. For the computational analyses, we consider the dynamics of the membrane potentials the main state variable (van den Broek et al., 2017).
**Intrinsic timescale** (τ_int_) - to quantify the characteristic timescale of population activity, we look athe autocorre lation function of each neuron’s membrane potentials:

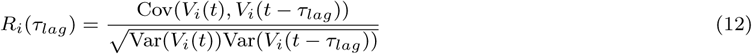 For each neuron, we fit the autocorrelation by an exponential function:

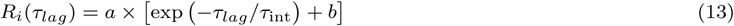

where *τ*_int_ specifies the decay time constant, characterizing the neuron’s intrinsic timescale.

#### 6.4.3 Processing capacity

To analyse the processing capacity of the networks, following (Dambre et al., 2012), we begin by defining an input sequence *u*[*n*], of finite total length *T*, comprising values independent- and identically drawn (i.i.d.) from a pre-determined probability distribution *p*(*u*). Since we are interested in measuring the circuit’s generic processing properties and not specific transformations on specific inputs, considering the input a random variable ensures that it has no pre-imposed structure so that any measured structure reflects only the system’s intrinsic properties and not the acquisition of structural relations present in the input. We set *p*(*u*) to be the uniform distribution over the interval [0,1]. This input sequence is then directly encoded into the circuit as explained above (Section 6.4.1).

The circuit’s initial states are randomized (*V*_o_ ~ *𝒰*_[*EL, Vth*]_) and the circuit is driven by the input, for a total simulation time of *T* × Δ*t*. In order to obtain accurate results and diminish potential errors and biases, we use a large sample size of *T* = 10^5^. The circuit state in response to the input is sampled at every Δ*t* ms, resulting in a collection of state vectors *x*[*n*] corresponding to a sample of the circuit state at time point *t*\* = *n* × Δ*t* ms. The resulting state matrix *X* ∊ ℝ _N _E_ ×*T*_ and the corresponding input *u* ∊ ℝ_1×*T*_ will then be used to estimate the capacity.

The aim of the analysis is to quantify the system’s ability to carry out computations on *u*. For that purpose, we measure the capacity *C* to reconstruct time-dependent functions *z* on finite sequences of *k* inputs, *z*[*n*] = *z*(*u*^−k^ [*n*]), from the state of the system, using a simple linear estimator:

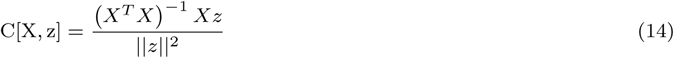

The numerator in the capacity measure corresponds to the linear estimator that minimizes the quadratic error between the target function to be reconstructed *z* and its linear estimate *ẑ:*

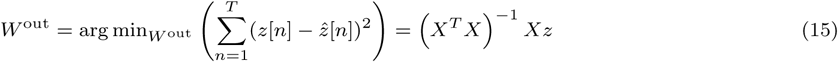

where

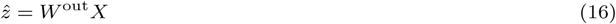

For any given function *z* and observed states *X*, C[X, z] is normalized, such that, in a system that allows perfect reconstruction of *z*, C[X, z] = 1, meaning that there exists a linear combination of *x*[*n*] that equals *z*[*n*], for all *n* On the other hand, a capacity of 0 indicates that it is not possible to even partially reconstruct the target function. Evaluating C[X, z] for large sets of target functions *y*_{_*l*} = {*z*_1_,…, *z_L_*}, allows us to gain insights into the information processing capacity of the system. If the evaluated functions are sufficiently distinct (preferably orthogonal), their corresponding capacities measure independent properties and provide independent information about how the system computes. As such,we systematically probe the capacity space by evaluating the complete set of orthonormal basis functions of *u*, using finite products of normalized Legendre polynomials:

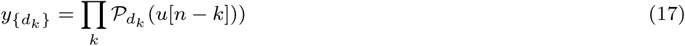

where 𝒫*_d_k__* (.) is the Legendre polynomial of degree *d_k_* ≥ 0:

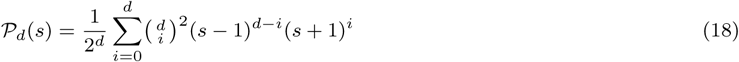

and is a function of input *u* delayed by *k* steps. The total capacity then corresponds to the sum of the individual capacities for a given set of target functions *y*{*d_k_*}:

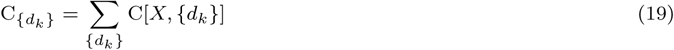

Naturally, we use finite data and a finite set of indices *d* to evaluate the capacities, leading to an unavoidable underestimation of the total capacity.

**Linear memory and nonlinearity** Using the notations introduced above, and following (Dambre et al., 2012; Jaeger, 2002), we can consider the linear memory capacity as the total capacity associated with linear functions:

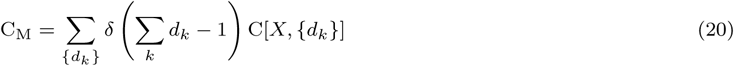

for a maximum polynomial degree of *d* = 1, i.e. each of the functions tested corresponds to a delayed version of the input:

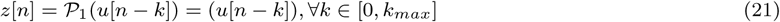

Accordingly, the capacity associated with non/linear functions corresponds to *d* ≥ 2. For more details on the implementation, consult (Dambre et al., 2012) and the code we provide in the Supplementary Materials.

### 6.5 Numerical simulations, implementation and data analysis

All the work presented in this manuscript was implemented using the Neural Microcircuit Simulation and Analysis Toolkit (NMSAT) (Duarte et al., 2017b), a python package designed to provide the first steps towards complex microcircuit benchmarking, as suggested and exemplified in this study. The core simulation engine running all the numerical simulations is NEST. Due to the specificities of this project, we used a modified version of NEST 2.10.0 (Bos et al., 2015), which includes all the models used in this manuscript (some of which are not available in the main release). A complete code package is provided in the supplementary materials that implements project-specific functionality to the framework, allowing the reproduction of all the numerical experiments presented in this manuscript. Computing resources were provided by the JARA-HPC Vergabegremium on the supercomputer JURECA (Krause and Thornig, 2016) at Forschungszentrum Jülich. All numerical simulations were performed at a resolution of 0.1 ms, using the GSL implementation of the adaptive fourth-order Runge-Kutta method.

## 7 Acknowledgements

The authors would like to acknowledge the contributions of Alexander Seeholzer in the initial conception and implementation of the neuron model used throughout this study. The authors would like to thank Maximillian Schmidt, Sven Goedecke and, in particular, Jannis Schuecker, for important suggestions and discussions at various stages of the project. The authors gratefully acknowledge the computing time granted by the JARA-HPC Vergabegremium on the supercomputer JURECA at Forschungszentrum Jülich, as well as the Neurobiology of Language group at the Max Plank Institute for Psycholinguistics, for valuable and useful discussions throughout the project. We acknowledge partial support by the Erasmus Mundus Joint Doctoral Program EuroSPIN, the German Ministry for Education and Research (Bundesministerium für Bildung und Forschung) BMBF Grant 01GQ0420 to BCCN Freiburg, the Initiative and Networking Fund of the Helmholtz Association and the Helmholtz Portfolio theme “Supercomputing and Modeling for the Human Brain”.

## Footnotes

1 http://www.neuroelectro.org/

2 http://www.brain-map.org

3 http://neuromorpho.org

4 http://neuroelectro.org

5 https://bbc.epfl.ch/nmc-portal

6 https://icg.neurotheory.ox.ac.uk

## References

L. F. Abbott and Wade G. Regehr. Synaptic computation. Nature, 431(7010):796–803, 2004. ISSN 0028-0836. doi: 10.1038/nature03010. URL http://www.nature.com/doifinder/10.1038/nature03010.

L F Abbott, Brian DePasquale, and Raoul-Martin Memmesheimer. Building functional networks of spiking model neurons. Nature Neuroscience, 19(3):350–355, 2016. ISSN 1097-6256. doi: 10.1038/nn.4241. URL http://www.nature.com/doifinder/10.1038/nn.4241.

D J Amit and N Brunel. Model of global spontaneous activity and local structured activity during delay periods in the cerebral cortex. Cerebral Cortex, 7(3):237–252, 1997a. ISSN 1047-3211. doi: 10.1093/cercor/7.3.237.

D J Amit and N Brunel. Full-Text. Cerebral Cortex, 7(3):237–252, 1997b.

M C Angulo, J Rossier, and E Audinat. Postsynaptic glutamate receptors and integrative properties of fast-spiking interneurons in the rat neocortex. Journal of neurophysiology, 82(3):1295–1302, sep 1999. ISSN 0022-3077. URL http://www.ncbi.nlm.nih.gov/pubmed/10482748.

Giorgio A Ascoli, Duncan E Donohue, and Maryam Halavi. NeuroMorpho.Org: a central resource for neuronal morphologies. The Journal of neuroscience: the official journal of the Society for Neuroscience, 27(35):9247–51, aug 2007. ISSN 1529-2401. doi: 10.1523/JNEUROSCI.2055-07.2007. URL http://www.ncbi.nlm.nih.gov/pubmed/17728438.

Giorgio A. Ascoli, Lidia Alonso-Nanclares, Stewart A. Anderson, German Barrionuevo, Ruth Benavides-Piccione, Andreas Burkhalter, György Buzsáki, Bruno Cauli, Javier DeFelipe, Alfonso Fairén, Dirk Feldmeyer, Gord Fishell, Yves Fregnac, Tamas F. Freund, Daniel Gardner, Esther P. Gardner, Jesse H. Goldberg, Moritz Helmstaedter, Shaul Hestrin, Fuyuki Karube, Zoltán F. Kisvárday, Bertrand Lambolez, David A. Lewis, Oscar Marin, Henry Markram, Alberto Muñoz, Adam Packer, Carl C. H. Petersen, Kathleen S. Rockland, Jean Rossier, Bernardo Rudy, Peter Somogyi, Jochen F. Staiger, Gabor Tamas, Alex M. Thomson, Maria Toledo-Rodriguez, Yun Wang, David C. West, and Rafael Yuste. Petilla terminology: nomenclature of features of GABAergic interneurons of the cerebral cortex. Nature Reviews Neuroscience, 9(7):557–568, jul 2008. ISSN 1471-003X. doi: 10.1038/nrn2402. URL http://www.nature.com/doifinder/10.1038/nrn2402.

M. Avermann, C. Tomm, C. Mateo, W. Gerstner, and C. C. H. Petersen. Microcircuits of excitatory and inhibitory neurons in layer 2/3 of mouse barrel cortex. Journal of Neurophysiology, 107(11):3116–3134, 2012. ISSN 0022-3077. doi: 10.1152/jn.00917.2011. URL http://jn.physiology.org/cgi/doi/10.1152/jn.00917.2011.

Mireille Bélanger, Igor Allaman, and Pierre J. Magistretti. Brain energy metabolism: Focus on Astrocyte-neuron metabolic cooperation, dec 2011. ISSN 15504131. URL http://www.sciencedirect.com/science/article/pii/S1550413111004207.

Brett L. Benedetti, Yoshio Takashima, Jing A. Wen, Joanna Urban-Ciecko, and Alison L. Barth. Differential wiring of layer 2/3 neurons drives sparse and reliable firing during neocortical development. Cerebral Cortex, 23(11):2690–2699, nov 2013. ISSN 10473211. doi: 10.1093/cercor/bhs257. URL http://www.ncbi.nlm.nih.gov/pubmed/22918982http://www.pubmedcentral.nih.gov/articlerender.fcgi?artid=PMC3792743.

Martin Boerlin, Christian K. Machens, and Sophie Denéve. Predictive Coding of Dynamical Variables in Balanced Spiking Networks. PLoS Computational Biology, 9(11), 2013. ISSN 1553734X. doi: 10.1371/journal.pcbi.1003258. URL http://journals.plos.org/ploscompbiol/article/file?id=10.1371/journal.pcbi.1003258&type=printable. 29/46

Hannah Bos, Jan Morrison, Abigail Peyser, Alexander Hahne, Moritz Helias, Susanne Kunkel, Tammo Ippen, Jochen Martin Eppler, Maximilian Schmidt, Alex Seeholzer, Mikael Djurfeldt, Sandra Diaz, Janne Morén, Rajalekshmi Deepu, Teo Stocco, Moritz Deger, Frank Michler, and Hans Ekkehard Plesser. Nest 2.10.0. page DOI: 10.5281/ZENODO.44222, 2015. doi: 10.5281/ZENODO.44222. URL https://zenodo.org/record/44222%5Cnhttps://doi.org/10.5281/zenodo.44222{#}.WNkIR4JBJ5o.mendeley&title=NEST2.10.0&description=NESTisahighlyscalablesimulatorfornetworksofpointorfew-compartmentspikingneuronmodels. Itincludesmultiplesynapticp https://doi.org/10.5281/zenodo.44222{#}.WNkIR4JBJ5o.mendeley&title=NEST2.10.0&description=NESTisahighlyscalablesimulatorfornetworksofpointorfew-compartmentspikingneuronmodels. Itincludesmultiplesynapticp.

Tiago Branco and Michael Häusser. The single dendritic branch as a fundamental functional unit in the nervous system. Current Opinion in Neurobiology, 20(4):494–502, 2010. ISSN 09594388. doi: 10.1016/j.conb.2010.07.009.

Solange P. Brown and Shaul Hestrin. Intracortical circuits of pyramidal neurons reflect their long-range axonal targets. Nature, 457(7233):1133–1136, feb 2009. ISSN 0028-0836. doi: 10.1038/nature07658. URL http://www.nature.com/doifinder/10.1038/nature07658.

R. M. Bruno. Cortex Is Driven by Weak but Synchronously Active Thalamocortical Synapses. Science, 312(5780):1622–1627, jun 2006. ISSN 0036-8075. doi: 10.1126/science.1124593. URL http://www.sciencemag.org/cgi/doi/10.1126/science.1124593.

György Buzsáki and Kenji Mizuseki. The log-dynamic brain: how skewed distributions affect network operations. Nature Reviews Neuroscience, 15(4):264–278, feb 2014. ISSN 1471-003X. doi: 10.1038/nrn3687. URL http://www.nature.com/doifinder/10.1038/nrn3687.

Nicholas Cain, Ramakrishnan Iyer, Christof Koch, and Stefan Mihalas. The Computational Properties of a Simplified Cortical Column Model. PLOS Computational Biology, 12(9):e1005045, sep 2016. ISSN 1553-7358. doi: 10.1371/journal.pcbi.1005045. URL http://dx.plos.org/10.1371/journal.pcbi.1005045.

Matteo Carandini. From circuits to behavior: a bridge too far? Nature Neuroscience, 15(4):507–509, mar 2012. ISSN 1097-6256. doi: 10.1038/nn.3043. URL http://www.nature.com/doifinder/10.1038/nn.3043.

Jessica A. Cardin. Dissecting local circuits in vivo: Integrated optogenetic and electrophysiology approaches for exploring inhibitory regulation of cortical activity. Journal of Physiology Paris, 106 (3-4):104–111, may 2012. ISSN 09284257. doi: 10.1016/j.jphysparis.2011.09.005. URL http://www.ncbi.nlm.nih.gov/pubmed/21958624http://www.pubmedcentral.nih.gov/articlerender.fcgi?artid=PMC3277809http://linkinghub.elsevier.com/retrieve/pii/S0928425711000313.

Rishidev Chaudhuri, Alberto Bernacchia, and Xiao Jing Wang. A diversity of localized timescales in network activity. eLife, 3:e01239, 2014. ISSN 2050084X. doi: 10.7554/eLife.01239.

Anne K Churchland and L F Abbott. Conceptual and technical advances define a key moment for theoretical neuroscience. Nature Neuroscience, 19(3):348–349, feb 2016. ISSN 1097-6256. doi: 10.1038/nn.4255. URL http://www.nature.com/doifinder/10.1038/nn.4255.

Christine E Collins, David C Airey, Nicole A Young, Duncan B Leitch, and Jon H Kaas. Neuron densities vary across and within cortical areas in primates. Proceedings of the National Academy of Sciences of the United States of America, 107(36):15927–32, sep 2010. ISSN 1091-6490. doi: 10.1073/pnas.1010356107. URL http://www.ncbi.nlm.nih.gov/pubmed/20798050http://www.pubmedcentral.nih.gov/articlerender.fcgi?artid=PMC2936588.

B W Connors, R C Malenka, and L R Silva. Two inhibitory postsynaptic potentials, and GABAA and GABAB receptor-mediated responses in neocortex of rat and cat. The Journal of Physiology, 406(1): 443–468, 1988. ISSN 00223751. doi: 10.1113/jphysiol.1988.sp017390. URL http://doi.wiley.com/10.1113/jphysiol.1988.sp017390.

Sylvain Crochet and Carl C H Petersen. Correlating whisker behavior with membrane potential in barrel cortex of awake mice. Nature Neuroscience, 9(5):608–610, 2006. ISSN 1097-6256. doi: 10.1038/nn1690. URL http://www.nature.com/doifinder/10.1038/nn1690.

Sylvain Crochet, James F A Poulet, Yves Kremer, and Carl C H Petersen. Synaptic mechanisms underlying sparse coding of active touch. Neuron, 69(6):1160–1175, mar 2011. ISSN 08966273. doi: 10.1016/j.neuron.2011.02.022. URL http://www.ncbi.nlm.nih.gov/pubmed/21435560http://linkinghub.elsevier.com/retrieve/pii/S0896627311001206.

Joni Dambre, David Verstraeten, Benjamin Schrauwen, and Serge Massar. Information Processing Capacity of Dynamical Systems. Scientific Reports, 2(1):514, jan 2012. ISSN 2045-2322. doi: 10.1038/srep00514. URL http://www.nature.com/articles/srep00514.

Charles Darwin. On the origin of the species by means of natural selection. 1859. ISBN 9781402756399 1402756399 9781402789595. URL http://www.ncbi.nlm.nih.gov/pubmed/15182499.

Nima Dehghani, Adrien Peyrache, Bartosz Telenczuk, Michel Le Van Quyen, Eric Halgren, Sydney S. Cash, Nicholas G. Hatsopoulos, and Alain Destexhe. Dynamic Balance of Excitation and Inhibition in Human and Monkey Neocortex. Scientific Reports, 6(1):23176, 2016. ISSN 2045-2322. doi: 10.1038/srep23176. URL http://www.nature.com/articles/srep23176.

K. Deisseroth, G. Feng, A. K. Majewska, G. Miesenbock, A. Ting, and M. J. Schnitzer. Next-Generation Optical Technologies for Illuminating Genetically Targeted Brain Circuits. Journal of Neuroscience, 26(41):10380–10386, oct 2006. ISSN 0270-6474. doi: 10.1523/JNEUROSCI.3863-06.2006. URL http://www.jneurosci.org/cgi/doi/10.1523/JNEUROSCI.3863-06.2006.

Sophie Denève and Christian K Machens. Efficient codes and balanced networks. Nature Neuroscience, 19 (3):375–382, 2016. ISSN 1097-6256. doi: 10.1038/nn.4243. URL http://www.nature.com/doifinder/10.1038/nn.4243.

a Destexhe, D Paré, Quebec Gk, and Denis Pare. Impact of network activity on the integrative properties of neocortical pyramidal neurons in vivo. Journal of neurophysiology, 81(4):1531–47, 1999. ISSN 0022-3077. URL http://www.ncbi.nlm.nih.gov/pubmed/10200189.

Alain Destexhe, Zachary F. Mainen, and Terrence J. Sejnowski. Synthesis of models for excitable membranes, synaptic transmission and neuromodulation using a common kinetic formalism. Journal of Computational Neuroscience, 1(3):195–230, 1994. ISSN 09295313. doi: 10.1007/BF00961734. URL http://link.springer.com/article/10.1007/BF00961734.

Alain Destexhe, Z F Mainen, and Terrence J. Sejnowski. Kinetic models of synaptic transmission. In Christof Koch and Idan Segev, editors, Methods in Neuronal Modeling, pages 1–25. MIT Press, Cambridge, MA, 2nd editio edition, 1998. URL http://cns.iaf.cnrs-gif.fr/abstracts/KSchap96.html.

Alain Destexhe, Michael Rudolph, and Denis Paré. The high-conductance state of neocortical neurons in vivo. Nature Reviews Neuroscience, 4(12):1019–1019, sep 2003. ISSN 1471-003X. doi: 10.1038/nrn1289. URL http://www.nature.com/doifinder/10.1038/nrn1289.

M. R. DeWeese and A. M. Zador. Non-Gaussian Membrane Potential Dynamics Imply Sparse, Synchronous Activity in Auditory Cortex. Journal of Neuroscience, 26(47):12206–12218, 2006. ISSN 0270-6474. doi: 10.1523/JNEUROSCI.2813-06.2006. URL http://www.jneurosci.org/cgi/doi/10.1523/JNEUROSCI.2813-06.2006.

M. Dichter and G. Ayala. Cellular mechanisms of epilepsy: a status report. Science, 237(4811):157–164, 1987. ISSN 0036-8075. doi: 10.1126/science.3037700. URL http://www.sciencemag.org/cgi/doi/10.1126/science.3037700.

Anja L. Dorrn, Kexin Yuan, Alison J. Barker, Christoph E. Schreiner, and Robert C. Froemke. Developmental sensory experience balances cortical excitation and inhibition. Nature, 465(7300):932–936, jun 2010. ISSN 0028-0836. doi: 10.1038/nature09119. URL http://www.nature.com/doifinder/10.1038/nature09119.

Alan D. Dorval. Probability distributions of the logarithm of inter-spike intervals yield accurate entropy estimates from small datasets. Journal of Neuroscience Methods, 173(1):129–139, aug 2008. ISSN 01650270. doi: 10.1016/j.jneumeth.2008.05.013. URL http://www.pubmedcentral.nih.gov/articlerender.fcgi?artid=2610469&tool=pmcentrez&rendertype=abstract.

Renato Duarte and Abigail Morrison. Dynamic stability of sequential stimulus representations in adapting neuronal networks. Frontiers in Computational Neuroscience, 8(October):124, 2014. ISSN 1662-5188. doi: 10.3389/fncom.2014.00124. URL http://journal.frontiersin.org/article/10.3389/fncom.2014.00124/abstract.

Renato Duarte, Alexander Seeholzer, Karl Zilles, and Abigail Morrison. Synaptic patterning and the timescales of cortical dynamics. Current Opinion in Neurobiology, 43:156–165, 2017a. ISSN 18736882. doi: 10.1016/j.conb.2017.02.007. URL http://www.sciencedirect.com/science/article/pii/S0959438817300545.

Renato Duarte, Barna Zajzon, and Abigail Morrison. Neural Microcircuit Simulation And Analysis Toolkit. Zenodo, jan 2017b. doi: 10.5281/ZEN0D0.582645. URL https://zenodo.org/record/582645.

A. S. Ecker, P. Berens, G. A. Keliris, M. Bethge, N. K. Logothetis, and A. S. Tolias. Decorrelated Neuronal Firing in Cortical Microcircuits. Science, 327(5965):584–587, jan 2010. ISSN 0036-8075. doi: 10.1126/science.1179867. URL http://www.sciencemag.org/cgi/doi/10.1126/science.1179867.

G. M. Edelman and J. A. Gally. Degeneracy and complexity in biological systems. Proceedings of the National Academy of Sciences, 98(24):13763–13768, nov 2001. ISSN 0027-8424. doi: 10.1073/pnas.231499798. URL http://www.pnas.org/cgi/doi/10.1073/pnas.231499798.

Chris Eliasmith and Charles H Anderson. Neural Engineering: Computation Representation and Dyamics in Neurobiological Systems, volume 19. MIT Press, 1991. ISBN 9781461452263. doi: 10.1007/BF02367401.

Pierre Enel, Emmanuel Procyk, René Quilodran, and Peter Ford Dominey. Reservoir Computing Properties of Neural Dynamics in Prefrontal Cortex. PLoS Computational Biology, 12(6):e1004967, 2016. ISSN 15537358. doi: 10.1371/journal.pcbi.1004967. URL http://dx.plos.org/10.1371/journal.pcbi.1004967.

U. Ernst and K. Pawelzik. Sensible Balance. Science, 334(6062):1507–1508, dec 2011. ISSN 0036-8075. doi: 10.1126/science.1216483. URL http://www.ncbi.nlm.nih.gov/pubmed/22174239.

Dirk Feldmeyer, Joachim Lübke, and Bert Sakmann. Efficacy and connectivity of intracolumnar pairs of layer 2/3 pyramidal cells in the barrel cortex of juvenile rats. The Journal of Physiology, 575(2): 583–602, sep 2006. ISSN 00223751. doi: 10.1113/jphysiol.2006.105106. URL http://doi.wiley.com/10.1113/jphysiol.2006.105106.

Barbara L. Finlay and Ryutaro Uchiyama. Developmental mechanisms channeling cortical evolution. Trends in Neurosciences, 38(2):69–76, 2015. ISSN 1878108X. doi: 10.1016/j.tins.2014.11.004. URL http://dx.doi.org/10.1016/j.tins.2014.11.004.

Elodie Fino, Adam M. Packer, and Rafael Yuste. The Logic of Inhibitory Connectivity in the Neocortex. The Neuroscientist, 19(3):228–237, 2013. ISSN 1073-8584. doi: 10.1177/1073858412456743. URL http://journals.sagepub.com/doi/10.1177/1073858412456743.

Karl Friston. A theory of cortical responses. Philosophical transactions of the Royal Society of London. Series B, Biological sciences, 360(1456):815–36, 2005. ISSN 0962-8436. doi: 10.1098/rstb.2005.1622. URL http://www.pubmedcentral.nih.gov/articlerender.fcgi?artid=1569488{%}7B&{%}7Dtool=pmcentrez{%}7B&{%}7Drendertype=abstract.

Peiran Gao and Surya Ganguli. On simplicity and complexity in the brave new world of large-scale neuroscience. Current Opinion in Neurobiology, 32:148–155, 2015. ISSN 18736882. doi: 10.1016/j.conb.2015.04.3. URL http://dx.doi.org/10.1016Zj.conb.2015.04.003.

Luc J. Gentet, Michael Avermann, Ferenc Matyas, Jochen F. Staiger, and Carl C H Petersen. Membrane Potential Dynamics of GABAergic Neurons in the Barrel Cortex of Behaving Mice. Neuron, 65(3): 422–435, 2010. ISSN 08966273. doi: 10.1016/j.neuron.2010.01.006. URL http://dx.doi.org/10.1016/j.neuron.2010.01.006.

W. Gerstner, H. Sprekeler, and G. Deco. Theory and Simulation in Neuroscience. Science, 338(6103): 60–65, 2012. ISSN 0036-8075. doi: 10.1126/science.1227356. URL http://www.sciencemag.org/cgi/doi/10.1126/science.1227356.

Wulfram Gerstner and Werner M. Kistler. Spiking Neuron Models. Cambridge University Press, 2002. ISBN 9780511815706. doi: 10.1017/CBO9780511815706. URL http://ebooks.cambridge.org/ref/id/CB09780511815706.

Wulfram Gerstner, Werner M. Kistler, Richard Naud, and Liam Paninski. Neuronal Dynamics - from single neurons to networks and models of cognition. Cambridge University Press, 2014. ISBN 9780874216561. doi: 10.1007/s13398-014-0173-7.2. URL https://books.google.de/books?id=D4j2AwAAQBAJ.

P. Getting. Emerging Principles Governing The Operation Of Neural Networks. Annual Review of Neuroscience, 12(1):185–204, mar 1989. ISSN 0147006X. doi: 10.1146/annurev.neuro.12.1.185. URL http://neuro.annualreviews.org/cgi/doi/10.1146/annurev.neuro.12.1.185.

Julijana Gjorgjieva, Guillaume Drion, and Eve Marder. Computational implications of biophysical diversity and multiple timescales in neurons and synapses for circuit performance. Current Opinion in Neurobiology, 37(Table 1):44–52, 2016. ISSN 18736882. doi: 10.1016/j.conb.2015.12.008. URL http://dx.doi.org/10.1016/j.conb.2015.12.008.

P. Greengard. The Neurobiology of Slow Synaptic Transmission. Science, 294(5544):1024–1030, nov 2001. ISSN 00368075. doi: 10.1126/science.294.5544.1024. URL http://www.sciencemag.org/cgi/doi/10.1126/science.294.5544.1024.

Ingo H. Greger, Jake F. Watson, and Stuart G. Cull-Candy. Structural and Functional Architecture of AMPA-Type Glutamate Receptors and Their Auxiliary Proteins. Neuron, 94(4):713–730, may 2017. ISSN 10974199. doi: 10.1016/j.neuron.2017.04.009. URL http://www.ncbi.nlm.nih.gov/pubmed/28521126.

A. Gupta. Organizing Principles for a Diversity of GABAergic Interneurons and Synapses in the Neocortex. Science, 287(5451):273–278, jan 2000. ISSN 00368075. doi: 10.1126/science.287.5451.273. URL http://www.sciencemag.org/cgi/doi/10.1126/science.287.5451.273.

Anirudh Gupta, Yun Wang, and Henry Markram. Organizing Principles for a Diversity of GABAergic Interneurons and Synapses in the Neocortex Diversity of GABAergic Synapses. Science, 287(5451): 273–8, 2000. URL http://science.sciencemag.org/content/sci/287/5451/273.full.pdf.

Stefan Haeusler and Wolfgang Maass. A statistical analysis of information-processing properties of lamina-specific cortical microcircuit models. Cerebral Cortex, 17(1):149–162, 2007. ISSN 10473211. doi: 10.1093/cercor/bhj132.

Kenneth D. Harris and Thomas D. Mrsic-Flogel. Cortical connectivity and sensory coding. Nature, 503(7474):51–58, 2013. ISSN 0028-0836. doi: 10.1038/nature12654. URL http://www.nature.com/doifinder/10.1038/nature12654.

Kenneth D Harris and Gordon M G Shepherd. The neocortical circuit: themes and variations. Nature Neuroscience, 18(2):170–181, 2015. ISSN 1097-6256. doi: 10.1038/nn.3917. URL http://www.nature.com/doifinder/10.1038/nn.3917.

Paul M. Harrison, Laurent Badel, Mark J. Wall, and Magnus J.E. Richardson. Experimentally Verified Parameter Sets for Modelling Heterogeneous Neocortical Pyramidal-Cell Populations. PLoS Computational Biology, 11(8):e1004165, aug 2015. ISSN 15537358. doi: 10.1371/journal.pcbi.1004165. URL http://dx.plos.org/10.1371/journal.pcbi.1004165.

Michael J Hawrylycz, Ed S Lein, Angela L Guillozet-Bongaarts, Elaine H Shen, Lydia Ng, Jeremy A Miller, Louie N van de Lagemaat, Kimberly A Smith, Amanda Ebbert, Zackery L Riley, Chris Abajian, Christian F Beckmann, Amy Bernard, Darren Bertagnolli, Andrew F Boe, Preston M Cartagena, M Mallar Chakravarty, Mike Chapin, Jimmy Chong, Rachel A Dalley, Barry David Daly, Chinh Dang, Suvro Datta, Nick Dee, Tim A Dolbeare, Vance Faber, David Feng, David R Fowler, Jeff Goldy, Benjamin W Gregor, Zeb Haradon, David R Haynor, John G Hohmann, Steve Horvath, Robert E Howard, Andreas Jeromin, Jayson M Jochim, Marty Kinnunen, Christopher Lau, Evan T Lazarz, Changkyu Lee, Tracy A Lemon, Ling Li, Yang Li, John A Morris, Caroline C Overly, Patrick D Parker, Sheana E Parry, Melissa Reding, Joshua J Royall, Jay Schulkin, Pedro Adolfo Sequeira, Clifford R Slaughterbeck, Simon C Smith, Andy J Sodt, Susan M Sunkin, Beryl E Swanson, Marquis P Vawter, Derric Williams, Paul Wohnoutka, H Ronald Zielke, Daniel H Geschwind, Patrick R Hof, Stephen M Smith, Christof Koch, Seth G N Grant, and Allan R Jones. An anatomically comprehensive atlas of the adult human brain transcriptome. Nature, 489(7416):391–9, sep 2012. ISSN 1476-4687. doi: 10.1038/nature11405. URL http://dx.doi.org/10.1038/nature11405.

M. Helmstaedter, C. P J de Kock, D. Feldmeyer, R. M. Bruno, and B. Sakmann. Reconstruction of an average cortical column in silico. Brain Research Reviews, 55(2 SPEC. ISS.):193–203, 2007. ISSN 01650173. doi: 10.1016/j.brainresrev.2007.07.011. URL http://linkinghub.elsevier.com/retrieve/pii/S0165017307001361.

M. Helmstaedter, J. F. Staiger, B. Sakmann, and D. Feldmeyer. Efficient Recruitment of Layer 2/3 Interneurons by Layer 4 Input in Single Columns of Rat Somatosensory Cortex. Journal of Neuroscience, 28(33):8273–8284, 2008. ISSN 0270-6474. doi: 10.1523/JNEUROSCI.5701-07.2008. URL http://www.jneurosci.org/cgi/doi/10.1523/JNEUROSCI.5701-07.2008.

Moritz Helmstaedter, Bert Sakmann, and Dirk Feldmeyer. L2/3 Interneuron groups defined by multiparameter analysis of axonal projection, dendritic geometry, and electrical excitability. Cerebral Cortex, 19(4):951–962, 2009. ISSN 10473211. doi: 10.1093/cercor/bhn130.

Jeremy M. Henley and Kevin A. Wilkinson. Synaptic AMPA receptor composition in development, plasticity and disease. Nature Reviews Neuroscience, 17(6):337–350, apr 2016. ISSN 1471-003X. doi: 10.1038/nrn.2016.37. URL http://www.nature.com/doifinder/10.1038/nrn.2016.37.

Matthias H Hennig. Theoretical models of synaptic short term plasticity. Frontiers in computational neuroscience, 7(April):45, 2013. ISSN 1662-5188. doi: 10.3389/fncom.2013.00045. URL http://www.pubmedcentral.nih.gov/articlerender.fcgi?artid=3630333&tool=pmcentrez&rendertype=abstract.

A. V. M. Herz, T. Gollisch, C. K. Machens, and D. Jaeger. Modeling Single-Neuron Dynamics and Computations: A Balance of Detail and Abstraction. Science, 314(5796):80–85, oct 2006. ISSN 0036-8075. doi: 10.1126/science.1127240. URL http://www.sciencemag.org/cgi/doi/10.1126/science.1127240.

Shaul Hestrin. Different glutamate receptor channels mediate fast excitatory synaptic currents in inhibitory and excitatory cortical neurons. Neuron, 11(6):1083–1091, 1993. ISSN 08966273. doi: 10.1016/0896-6273(93)90221-C.

E Hill, M Kalloniatis, and S S Tan. Glutamate, GABA and precursor amino acids in adult mouse neocortex: cellular diversity revealed by quantitative immunocytochemistry. Cerebral cortex (New York, N.Y.: 1991), 10(11):1132–42, 2000. ISSN 1047-3211. URL http://www.ncbi.nlm.nih.gov/pubmed/11053233.

Felix Z. Hoffmann and Jochen Triesch. Nonrandom network connectivity comes in pairs. Network Neuroscience, 1(1):31–41, feb 2017. ISSN 2472-1751. doi: 10.1162/NETN_a_00004. URL http://www.mitpressjournals.org/doi/10.1162/NETN{_}a{_}00004.

Jochen H.O. Hoffmann, H. S. Meyer, Arno C. Schmitt, Jakob Straehle, Trinh Weitbrecht, Bert Sakmann, and Moritz Helmstaedter. Synaptic conductance estimates of the connection between local inhibitor interneurons and pyramidal neurons in layer 2/3 of a cortical column. Cerebral Cortex, 25(11):4415–4429, 2015. ISSN 14602199. doi: 10.1093/cercor/bhv039. URL http://www.ncbi.nlm.nih.gov/pubmed/25761638.

Anthony Holtmaat and Karel Svoboda. Experience-dependent structural synaptic plasticity in the mammalian brain. Nature Reviews Neuroscience, 10(9):647–658, sep 2009. ISSN 1471-003X. doi: 10.1038/nrn2721. URL http://www.ncbi.nlm.nih.gov/pubmed/19693029.

J.J. Hopfield. Physics, Computation, and Why Biology Looks so Different, jul 1994. ISSN 00225193. URL http://www.sciencedirect.com/science/article/pii/S0022519384712112.

Mark D Humphries. The Goldilocks zone in neural circuits. eLife, 5, 2016. ISSN 2050-084X. doi: 10.7554/eLife.22735. URL http://elifesciences.org/lookup/doi/10.7554/eLife.22735.

Jeffry S. Isaacson and Massimo Scanziani. How inhibition shapes cortical activity. Neuron, 72(2):231–243, oct 2011. ISSN 08966273. doi: 10.1016/j.neuron.2011.09.027. URL http://www.pubmedcentral.nih.gov/articlerender.fcgi?artid=3236361&tool=pmcentrez&rendertype=abstract.

H Jaeger. Short term memory in echo state networks. GMD Report 152, page 60, 2002. URL papers: http//78a99879-71e7-4c85-9127-d29c2b4b416b/Paper/p14153{%}5Cnhttp://neuron-ai.tuke.sk/{∼}bundzel/diploma{_}theses{_}students/2006/MartinSramko-EchoStateNNinPrediction/STMEchoStatesTechRep.pdf.

C E Jahr and C F Stevens. Voltage dependence of NMDA-activated macroscopic conductances predicted by single-channel kinetics. The Journal of neuroscience, 10(9):3178–3182, 1990. ISSN 0270-6474. doi: 1990/09/0100:01.

X. Jiang, S. Shen, F. Sinz, J. Reimer, C. R. Cadwell, P. Berens, A. S. Ecker, S. Patel, G. H. Denfield, E. Froudarakis, S. Li, E. Walker, and A. S. Tolias. Response to Comment on “Principles of connectivity among morphologically defined cell types in adult neocortex”. Science, 353(6304):1108–1108, 2016. ISSN 0036-8075. doi: 10.1126/science.aaf6102. URL http://www.sciencemag.org/cgi/doi/10.1126/science.aaf6102.

Hyo Jung Kang, Yuka Imamura Kawasawa, Feng Cheng, Ying Zhu, Xuming Xu, Mingfeng Li, André M M Sousa, Mihovil Pletikos, Kyle A Meyer, Goran Sedmak, Tobias Guennel, Yurae Shin, Matthew B Johnson, Zeljka Krsnik, Simone Mayer, Sofia Fertuzinhos, Sheila Umlauf, Steven N Lisgo, Alexander Vortmeyer, Daniel R Weinberger, Shrikant Mane, Thomas M Hyde, Anita Huttner, Mark Reimers, Joel E Kleinman, and Nenad Sestan. Spatio-temporal transcriptome of the human brain. Nature, 478(7370):483–9, oct 2011. ISSN 1476-4687. doi: 10.1038/nature10523. URL http://www.ncbi.nlm.nih.gov/pubmed/22031440http://www.pubmedcentral.nih.gov/articlerender.fcgi?artid=PMC3566780http://www.pubmedcentral.nih.gov/articlerender.fcgi?artid=3566780&tool=pmcentrez&rendertype=abstract.

George Kastellakis, Denise J Cai, Sara C Mednick, Alcino J Silva, and Panayiota Poirazi. Synaptic clustering within dendrites: an emerging theory of memory formation. Progress in neurobiology, 126:19–35, mar 2015. ISSN 1873-5118. doi: 10.1016/j.pneurobio.2014.12.002. URL http://www.ncbi.nlm.nih.gov/pubmed/25576663http://www.pubmedcentral.nih.gov/articlerender.fcgi?artid=PMC4361279.

C. Koch. Complexity and the Nervous System. Science, 284(5411):96–98, apr 1999. ISSN 00368075. doi: 10.1126/science.284.5411.96. URL http://www.sciencemag.org/cgi/doi/10.1126/science.284.5411.96.

Christof Koch. Biophysics of Computation Information Processing in Single Neuron, volume 11. Oxford University Press, USA, 2004. ISBN 0-19-510491-9. doi: 10.3902/jnns.11.157_2. URL http://www.amazon.de/Biophysics-Computation-Information-Computational-Neuroscience/dp/0195181999.

G Köhr, Y De Koninck, and I Mody. Properties of NMDA receptor channels in neurons acutely isolated from epileptic (kindled) rats. The Journal of neuroscience: the official journal of the Society for Neuroscience, 13(8):3612–3627, 1993. ISSN 0270-6474.

A. A. Koulakov, T. Hromadka, and A. M. Zador. Correlated Connectivity and the Distribution of Firing Rates in the Neocortex. Journal of Neuroscience, 29(12):3685–3694, mar 2009. ISSN 0270-6474. doi: 10.1523/JNEUROSCI.4500-08.2009. URL http://www.jneurosci.org/cgi/doi/10.1523/JNEUR0SCI.4500-08.2009.

John W. Krakauer, Asif A. Ghazanfar, Alex Gomez-Marin, Malcolm A. MacIver, and David Poeppel. Neuroscience Needs Behavior: Correcting a Reductionist Bias. Neuron, 93(3):480–490, feb 2017. ISSN 10974199. doi: 10.1016/j.neuron.2016.12.041. URL http://www.ncbi.nlm.nih.gov/pubmed/28182904.

Dorian Krause and Philipp Thörnig. JURECA: General-purpose supercomputer at Jülich Supercomputing Centre. Journal of large-scale research facilities JLSRF, 2(0):A62, mar 2016. ISSN 2364-091X. doi: 10.17815/jlsrf-2-121. URL http://jlsrf.org/index.php/lsf/article/view/121.

J. Kremkow, A. Aertsen, and A. Kumar. Gating of Signal Propagation in Spiking Neural Networks by Balanced and Correlated Excitation and Inhibition. Journal of Neuroscience, 30(47):15760–15768, nov 2010. ISSN 0270-6474. doi: 10.1523/JNEUROSCI.3874-10.2010. URL http://www.jneurosci.org/cgi/doi/10.1523/JNEUR0SCI.3874-10.2010.

Yoshiyuki Kubota, Satoru Kondo, Masaki Nomura, Sayuri Hatada, Noboru Yamaguchi, Alsayed A. Mohamed, Fuyuki Karube, Joachim Lübke, and Yasuo Kawaguchi. Functional effects of distinct innervation styles of pyramidal cells by fast spiking cortical interneurons. eLife, 4(July 2015):1–27, 2015. ISSN 2050084X. doi: 10.7554/eLife.07919.

Yoshiyuki Kubota, Fuyuki Karube, Masaki Nomura, and Yasuo Kawaguchi. The Diversity of Cortical Inhibitory Synapses. Frontiers in Neural Circuits, 10:27, 2016. ISSN 1662-5110. doi: 10.3389/fncir.2016.00027. URL http://journal.frontiersin.org/Article/10.3389/fncir.2016.00027/abstract.

Arvind Kumar, Sven Schrader, Ad Aertsen, and Stefan Rotter. The High-Conductance State of Cortical Networks. Neural Computation, 20(1):1–43, jan 2008. ISSN 0899-7667. doi: 10.1162/neco.2008.20.1.1. URL http://www.mitpressjournals.org/doi/10.1162/neco.2008.20.1.1.

Fereshteh Lagzi and Stefan Rotter. Dynamics of competition between subnetworks of spiking neuronal networks in the balanced state. PLoS ONE, 10(9), 2015. ISSN 19326203. doi: 10.1371/journal.pone.0138947.

Sandrine Lefort, Christian Tomm, J. C. Floyd Sarria, and Carl C H Petersen. The Excitatory Neuronal Network of the C2 Barrel Column in Mouse Primary Somatosensory Cortex. Neuron, 61(2):301–316, jan 2009. ISSN 08966273. doi: 10.1016/j.neuron.2008.12.020. URL http://www.ncbi.nlm.nih.gov/pubmed/19186171http://linkinghub.elsevier.com/retrieve/pii/S0896627308010921.

J Leger, E Stern, A Aertsen, and D Heck. Synaptic integration in rat frontal cortex shaped by network activity. Journal of Neurophysiology, 93(93):281–293, 2005. ISSN 0022-3077. doi: 10.1152/jn.00067.2003. URL http://jn.physiology.org/content/93/1Z281.short.

David A. Lewis and Guillermo Gonzalez-Burgos. Intrinsic excitatory connections in the prefrontal cortex and the pathophysiology of schizophrenia. Brain Research Bulletin, 52(5):309–317, jul 2000. ISSN 03619230. doi: 10.1016/S0361-9230(99)00243-9. URL http://www.ncbi.nlm.nih.gov/pubmed/10922508.

John E. Lisman, Sridhar Raghavachari, and Richard W. Tsien. The sequence of events that underlie quantal transmission at central glutamatergic synapses. Nature Reviews Neuroscience, 8(8):597–609, 2007. ISSN 1471-003X. doi: 10.1038/nrn2191. URL http://www.nature.com/doifinder/10.1038/nrn2191.

Ashok Litwin-Kumar and Brent Doiron. Slow dynamics and high variability in balanced cortical networks with clustered connections. Nature Neuroscience, 15(11):1498–1505, nov 2012. ISSN 1097-6256. doi: 10.1038/nn.3220. URL http://www.nature.com/doifinder/10.1038/nn.3220.

J.-t. Lu, C-y. Li, J-P. Zhao, M-m. Poo, and X.-h. Zhang. Spike-Timing-Dependent Plasticity of Neocortical Excitatory Synapses on Inhibitory Interneurons Depends on Target Cell Type. Journal of Neuroscience, 27(36):9711–9720, sep 2007. ISSN 0270-6474. doi: 10.1523/JNEUROSCI.2513-07.2007. URL http://www.jneurosci.org/cgi/doi/10.1523/JNEUR0SCI.2513-07.2007.

A. Luczak, P. Bartho, S. L. Marguet, G. Buzsaki, and K. D. Harris. Sequential structure of neocortical spontaneous activity in vivo. Proceedings of the National Academy of Sciences, 104(1):347–352, jan 2007. ISSN 0027-8424. doi: 10.1073/pnas.0605643104. URL http://www.pnas.org/cgi/doi/10.1073/pnas.0605643104.

Wolfgang Maass, T Natschl\\”ager, and Henry Markram. Real-time computing without stable states: A new framework for neural computation based on perturbations. Neural computation, 14(11):2531–2560, 2002. ISSN 0899-7667. URL http://www.mitpressjournals.org/doi/abs/10.1162/089976602760407955?url{%}7B{_}{%}7Dver=Z39.88-2003{%}7B&{%}7Damp;rfr{%}7B{_}{%}7Did=ori:rid:crossref.org{%}7B&{%}7Damp;rfr{%}7B{_}{%}7Ddat=cr{%}7B{_}{%}7Dpub{%}7B{%}25{%}7D3Dncbi.nlm.nih.gov.

Wolfgang Maass, Thomas Natschl??ger, and Henry Markram. Fading memory and kernel properties of generic cortical microcircuit models. Journal of Physiology Paris, 98(4-6 SPEC. ISS.):315–330, jul 2004. ISSN 09284257. doi: 10.1016/j.jphysparis.2005.09.020. URL http://www.sciencedirect.com/science/article/pii/S0928425705000215?via{%}3Dihub.

Arianna Maffei. The many forms and functions of long term plasticity at GABAergic synapses. Neural Plasticity, 2011:1–9, 2011. ISSN 16875443. doi: 10.1155/2011/254724. URL http://www.hindawi.com/journals/np/2011/254724/.

J. Mappes and L. Lindstrom. How Did the Cuckoo Get Its Polymorphic Plumage? Science, 337(6094): 532–533, aug 2012. ISSN 0036-8075. doi: 10.1126/science.1225997. URL http://www.sciencemag.org/cgi/doi/10.1126/science.1225997.

Jorge Mariño, James Schummers, David C Lyon, Lars Schwabe, Oliver Beck, Peter Wiesing, Klaus Obermayer, and Mriganka Sur. Invariant computations in local cortical networks with balanced excitation and inhibition. Nature Neuroscience, 8(2):194–201, feb 2005. ISSN 1097-6256. doi: 10.1038/nn1391. URL http://www.nature.com/doifinder/10.1038/nn1391.

Florian Markowetz. All biology is computational biology. PLoS Biology, 15(3):e2002050, mar 2017. ISSN 15457885. doi: 10.1371/journal.pbio.2002050. URL http://www.ncbi.nlm.nih.gov/pubmed/28278152http://www.pubmedcentral.nih.gov/articlerender.fcgi?artid=PMC5344307.

Henry Markram. The blue brain project. Nature reviews. Neuroscience, 7(2):153–160, feb 2006. ISSN 1471-003X. doi: 10.1038/nrn1848. URL http://www.ncbi.nlm.nih.gov/pubmed/16429124.

Henry Markram, Eilif Muller, Srikanth Ramaswamy, Michael W Reimann, Marwan Abdellah, Carlos Aguado Sanchez, Anastasia Ailamaki, Lidia Alonso-Nanclares, Nicolas Antille, Selim Arsever, Guy Antoine Atenekeng Kahou, Thomas K Berger, Ahmet Bilgili, Nenad Buncic, Athanassia Chalimourda, Giuseppe Chindemi, Jean-Denis Courcol, Fabien Delalondre, Vincent Delattre, Shaul Druckmann, Raphael Dumusc, James Dynes, Stefan Eilemann, Eyal Gal, Michael Emiel Gevaert, Jean-Pierre Ghobril, Albert Gidon, Joe W Graham, Anirudh Gupta, Valentin Haenel, Etay Hay, Thomas Heinis, Juan B Hernando, Michael Hines, Lida Kanari, Daniel Keller, John Kenyon, Georges Khazen, Yihwa Kim, James G King, Zoltan Kisvarday, Pramod Kumbhar, Sébastien Lasserre, Jean-Vincent Le Bé, Bruno R C Magãlhaes, Angel Merchán-Pérez, Julie Meystre, Benjamin Roy Morrice, Jeffrey Muller, Alberto Muñoz-Céspedes, Shruti Muralidhar, Keerthan Muthurasa, Daniel Nachbaur, Taylor H Newton, Max Nolte, Aleksandr Ovcharenko, Juan Palacios, Luis Pastor, Rodrigo Perin, Rajnish Ranjan, Imad Riachi, José-Rodrigo Rodríguez, Juan Luis Riquelme, Christian Rössert, Konstantinos Sfyrakis, Ying Shi, Julian C Shillcock, Gilad Silberberg, Ricardo Silva, Farhan Tauheed, Martin Telefont, Maria Toledo-Rodriguez, Thomas Tränkler, Werner Van Geit, Jafet Villafranca Díaz, Richard Walker, Yun Wang, Stefano M Zaninetta, Javier DeFelipe, Sean L Hill, Idan Segev, and Felix Schürmann. Reconstruction and Simulation of Neocortical Microcircuitry. Cell, 163(2):456–92, oct 2015. ISSN 1097-4172. doi: 10.1016/j.cell.2015.09.029. URL http://www.ncbi.nlm.nih.gov/pubmed/26451489.

Shimon Marom. On the Precarious Path of Reverse Neuro-Engineering. Frontiers in Computational Neuroscience, 3(May):3–6, jan 2009. ISSN 16625188. doi: 10.3389/neuro.10.005.2009. URL http://journal.frontiersin.org/article/10.3389/neuro.10.005.2009/abstract.

Vivien Marx. A deep look at synaptic dynamics. Nature, 515(7526):293–297, 2014. ISSN 0028-0836. doi: 10.1038/515293a. URL http://www.nature.com/doifinder/10.1038/515293a.

David A. McCormick, Zhong Wang, and John Huguenard. Neurotransmitter control of neocortical neuronal activity and excitability. Cerebral Cortex, 3(5):387–398, sep 1993. ISSN 14602199. doi: 10.1093/cercor/3.5.387. URL http://cercor.oxfordjournals.org/content/3Z5/387.shorthttps://academic.oup.com/cercor/article-lookup/doi/10.1093/cercor/3.5.387.

Raoul Martin Memmesheimer and Marc Timme. Designing complex networks. Physica D: Nonlinear Phenomena, 224(1-2):182–201, 2006a. ISSN 01672789. doi: 10.1016/j.physd.2006.09.037.

Raoul Martin Memmesheimer and Marc Timme. Designing the dynamics of spiking neural networks. Physical Review Letters, 97(18):1881011–4, 2006b. ISSN 00319007. doi: 10.1103/PhysRevLett.97.188101.

S. Mensi, R. Naud, C. Pozzorini, M. Avermann, C. C. H. Petersen, and W. Gerstner. Parameter extraction and classification of three cortical neuron types reveals two distinct adaptation mechanisms. Journal of Neurophysiology, 107(6):1756–1775, 2012. ISSN 0022-3077. doi: 10.1152/jn.00408.2011. URL http://jn.physiology.org/cgi/doi/10.1152/jn.00408.2011.

David Meunier, Renaud Lambiotte, and Edward T. Bullmore. Modular and hierarchically modular organization of brain networks. Frontiers in Neuroscience, 4(DEC):1–11, 2010. ISSN 16624548. doi: 10.3389/fnins.2010.00200. URL http://www.ncbi.nlm.nih.gov/pmc/articles/PMC3000003/.

Gero Miesenböck. Optogenetic Control of Cells and Circuits. Annual Review of Cell and Developmental Biology, 27(1):731–758, nov 2011. ISSN 1081-0706. doi: 10.1146/annurev-cellbio-100109-104051. URL http://www.annualreviews.org/doi/10.1146/annurev-cellbio-100109-104051.

Alexandre W. Moreau and Dimitri M. Kullmann. NMDA receptor-dependent function and plasticity in inhibitory circuits. Neuropharmacology, 74:23–31, nov 2013. ISSN 18737064. doi: 10.1016/j.neuropharm.2013.03.4. URL http://www.ncbi.nlm.nih.gov/pubmed/23537500http://linkinghub.elsevier.com/retrieve/pii/S0028390813001007.

K. Morita. Possible Role of Dendritic Compartmentalization in the Spatial Working Memory Circuit. Journal of Neuroscience, 28(30):7699–7724, jul 2008. ISSN 0270-6474. doi: 10.1523/JNEUROSCI.0059-08.2008. URL http://www.jneurosci.org/cgi/doi/10.1523/JNEUR0SCI.0059-08.2008.

Abigail Morrison, Markus Diesmann, and Wulfram Gerstner. Phenomenological models of synaptic plasticity based on spike timing. Biological Cybernetics, 98(6):459–478, 2008. ISSN 03401200. doi: 10.1007/s00422-008-0233-1.

V. Mountcastle. An organizing principle for cerebral function: the unit model and the distributed system. In G M Edelman and V Mountcastle, editors, The Mindful Brain, pages 7–50. MIT Press, 1978. ISBN 0262550075. URL http://www.citeulike.org/group/8299/article/4545635.

Vernon B. Mountcastle. The columnar organization of the neocortex. Brain, 120(4):701–722, 1997. ISSN 00068950. doi: 10.1093/brain/120.4.701. URL http://www.ncbi.nlm.nih.gov/pubmed/9153131.

Sophia Mueller, Danhong Wang, Michael D. Fox, B. T Thomas Yeo, Jorge Sepulcre, Mert R. Sabuncu, Rebecca Shafee, Jie Lu, and Hesheng Liu. Individual Variability in Functional Connectivity Architecture of the Human Brain. Neuron, 77(3):586–595, 2013. ISSN 08966273. doi: 10.1016/j.neuron.2012.12.028. URL http://dx.doi.org/10.1016/j.neuron.2012.12.028.

C. I. O. Myme. The NMDA-to-AMPA Ratio at Synapses Onto Layer 2/3 Pyramidal Neurons Is Conserved Across Prefrontal and Visual Cortices. Journal of Neurophysiology, 90(2):771–779, 2003. ISSN 0022-3077. doi: 10.1152/jn.00070.2003. URL http://jn.physiology.org/cgi/doi/10.1152/jn.00070.2003.

G. T. Neske, S. L. Patrick, and B. W. Connors. Contributions of Diverse Excitatory and Inhibitory Neurons to Recurrent Network Activity in Cerebral Cortex. Journal of Neuroscience, 35(3):1089–1105, 2015. ISSN 0270-6474. doi: 10.1523/JNEUROSCI.2279-14.2015. URL http://www.jneurosci.org/cgi/doi/10.1523/JNEUR0SCI.2279-14.2015.

W. Nissen, A. Szabo, J. Somogyi, P. Somogyi, and K. P. Lamsa. Cell Type-Specific Long-Term Plasticity at Glutamatergic Synapses onto Hippocampal Interneurons Expressing either Parvalbumin or CB1 Cannabinoid Receptor. Journal of Neuroscience, 30(4):1337–1347, jan 2010. ISSN 0270-6474. doi: 10.1523/JNEUROSCI.3481-09.2010. URL http://www.jneurosci.org/cgi/doi/10.1523/JNEUR0SCI.3481-09.2010.

Daniel H. O’Connor, Simon P. Peron, Daniel Huber, and Karel Svoboda. Neural activity in barrel cortex underlying vibrissa-based object localization in mice. Neuron, 67(6):1048–1061, sep 2010. ISSN 08966273. doi: 10.1016/j.neuron.2010.08.026. URL http://www.ncbi.nlm.nih.gov/pubmed/20869600.

Michael Okun and Ilan Lampl. Balance of excitation and inhibition. Scholarpedia, 4(8):7467, 2009. ISSN 1941-6016. doi: 10.4249/scholarpedia.7467. URL https://link.springer.com/content/pdf/10.2991{%}2F978-94-6239-133-8{_}44.pdf{%}0Ahttp://www.scholarpedia.org/article/Balance{_}of{_}excitation{_}and{_}inhibition.

Open Science Collaboration. Estimating the reproducibility of psychological science. Science, 349 (6251):aac4716-aac4716, aug 2015. ISSN 0036-8075. doi: 10.1126/science.aac4716. URL http://www.sciencemag.org/cgi/doi/10.1126/science.aac4716.

Nancy A O’Rourke, Nicholas C Weiler, Kristina D Micheva, and Stephen J Smith. Deep molecular diversity of mammalian synapses: why it matters and how to measure it. Nature reviews. Neuroscience, 13(6):365–79, may 2012. ISSN 1471-0048. doi: 10.1038/nrn3170. URL http://www.pubmedcentral.nih.gov/articlerender.fcgi?artid=3670986{&}tool=pmcentrez{&}rendertype=abstract.

Adriane G. Otopalik, Alexander C. Sutton, Matthew Banghart, and Eve Marder. When complex neuronal structures may not matter. eLife, 6:e23508, feb 2017. ISSN 2050084X. doi: 10.7554/eLife.23508. URL https://elifesciences.org/articles/23508.

Marius Pachitariu, Carsen Stringer, Michael Okun, Peter Bartho, Kenneth Harris, Peter Latham, Maneesh Sahani, and Nicholas Lesica. Inhibitory control of shared variability in cortical networks. bioRxiv, (041103):041103, 2016. ISSN 2050-084X. doi: 10.1101/041103. URL http://biorxiv.org/lookup/doi/10.1101/041103.

N Palomero-Gallagher, Katrin Amunts, and Karl Zilles. Transmitter Receptor Distribution in the Human Brain. In A.W. Toga, editor, Brain Mapping, chapter 221, pages 261–275. Elsevier Academic Press, San Diego, 2015. doi: 10.1016/B978-0-12-397025-1.00221-9.

Nicola Palomero-Gallagher and Karl Zilles. Cortical layers: Cyto-, myelo-, receptor- and synaptic architecture in human cortical areas. NeuroImage, aug 2017. ISSN 10959572. doi: 10.1016/j.neuroimage.2017.08.35. URL http://linkinghub.elsevier.com/retrieve/pii/S1053811917306821.

Pierre Paoletti, Camilla Bellone, and Qiang Zhou. NMDA receptor subunit diversity: impact on receptor properties, synaptic plasticity and disease. Nature Reviews Neuroscience, 14(6):383–400, may 2013. ISSN 1471-003X. doi: 10.1038/nrn3504. URL http://www.nature.com/doifinder/10.1038/nrn3504.

H J Park and K Friston. Structural and functional brain networks: from connections to cognition. Science (New York, N.Y.), 342(6158):1238411, 2013. ISSN 0036-8075. doi: 10.1126/science.1238411.

R. D. Peng. Reproducible Research in Computational Science. Science, 334(6060):1226–1227, dec 2011. ISSN 0036-8075. doi: 10.1126/science.1213847. URL http://www.sciencemag.org/cgi/doi/10.1126/science.1213847.

R. Perin, T. K. Berger, and H. Markram. A synaptic organizing principle for cortical neuronal groups. Proceedings of the National Academy of Sciences, 108(13):5419–5424, mar 2011. ISSN 0027-8424. doi: 10.1073/pnas.1016051108. URL http://www.pnas.org/cgi/doi/10.1073/pnas.1016051108.

Volker Pernice, Moritz Deger, Stefano Cardanobile, and Stefan Rotter. The relevance of network micro-structure for neural dynamics. Frontiers in computational neuroscience, 7(June):72, 2013. ISSN 1662-5188. doi: 10.3389/fncom.2013.00072. URL http://www.pubmedcentral.nih.gov/articlerender.fcgi?artid=3671286{&}tool=pmcentrez{&}rendertype=abstract.

C. C. H. Petersen, T. T. G. Hahn, M. Mehta, A. Grinvald, and B. Sakmann. Interaction of sensory responses with spontaneous depolarization in layer 2/3 barrel cortex. Proceedings of the National Academy of Sciences, 100(23):13638–13643, nov 2003. ISSN 0027-8424. doi: 10.1073/pnas.2235811100. URL http://www.pnas.org/cgi/doi/10.1073/pnas.2235811100.

Carl C H Petersen and Sylvain Crochet. Synaptic Computation and Sensory Processing in Neocortical Layer 2/3. Neuron, 78(1):28–48, 2013. ISSN 08966273. doi: 10.1016/j.neuron.2013.03.020. URL http://linkinghub.elsevier.com/retrieve/pii/S0896627313002675.

Hyun-Jae Pi, Balázs Hangya, Duda Kvitsiani, Joshua I. Sanders, Z. Josh Huang, and Adam Kepecs. Cortical interneurons that specialize in disinhibitory control. Nature, 503(7477):521–524, 2013. ISSN 0028-0836. doi: 10.1038/nature12676. URL http://www.nature.com/doifinder/10.1038/nature12676.

Gualtiero Piccinini and Oron Shagrir. Foundations of computational neuroscience. Current Opinion in Neurobiology, 25:25–30, 2014. ISSN 09594388. doi: 10.1016/j.conb.2013.10.005. URL http://dx.doi.org/10.1016/j.conb.2013.10.005.

M. Pletikos, A. Sousa, G. Sedmak, K. Meyer, Y. Zhu, F. Cheng, M. Li, Y. Kawasawa, and N. Estan. Temporal specification and bilaterality of human neocortical topographic gene expression. Neuron, 81 (2):321–332, 2014. ISSN 08966273. doi: 10.1016/j.neuron.2013.11.018.

William F Podlaski, Alexander Seeholzer, Lukas N Groschner, Gero Miesenboeck, Rajnish Ranjan, and Tim P Vogels. ICGenealogy: Mapping the function of neuronal ion channels in model and experiment. bioRxiv, page 058685, jun 2016. doi: 10.1101/058685. URL https://www.biorxiv.org/content/early/2016/06/16/058685.

Tobias C. Potjans and Markus Diesmann. The cell-type specific cortical microcircuit: Relating structure and activity in a full-scale spiking network model. Cerebral Cortex, 24(3):785–806, 2014. ISSN 10473211. doi: 10.1093/cercor/bhs358.

James F A Poulet. Keeping an Eye on Cortical States. Neuron, 84(2):246–248, oct 2014. ISSN 10974199. doi: 10.1016/j.neuron.2014.10.005. URL http://www.ncbi.nlm.nih.gov/pubmed/25374350.

James F. A. Poulet and Carl C. H. Petersen. Internal brain state regulates membrane potential synchrony in barrel cortex of behaving mice. Nature, 454(7206):881–885, aug 2008. ISSN 0028-0836. doi: 10.1038/nature07150. URL http://www.nature.com/doifinder/10.1038/nature07150.

James F A Poulet, Laura M J Fernandez, Sylvain Crochet, and Carl C H Petersen. Thalamic control of cortical states. Nature Neuroscience, 15(3):370–372, mar 2012. ISSN 1097-6256. doi: 10.1038/nn.3035. URL http://www.nature.com/doifinder/10.1038/nn.3035.

Jean-Francois Poulin, Bosiljka Tasic, Jens Hjerling-Leffler, Jeffrey M Trimarchi, and Rajeshwar Awatramani. Disentangling neural cell diversity using single-cell transcriptomics. Nature Neuroscience, 19(9):1131–1141, aug 2016. ISSN 1097-6256. doi: 10.1038/nn.4366. URL http://www.nature.com/doifinder/10.1038/nn.4366.

Jonathan D. Power, Alexander L. Cohen, Steven M. Nelson, Gagan S. Wig, Kelly Anne Barnes, Jessica A. Church, Alecia C. Vogel, Timothy O. Laumann, Fran M. Miezin, Bradley L. Schlaggar, and Steven E. Petersen. Functional Network Organization of the Human Brain. Neuron, 72(4):665–678, nov 2011. ISSN 08966273. doi: 10.1016/j.neuron.2011.09.006. URL http://www.pubmedcentral.nih.gov/articlerender.fcgi?artid=3222858{&}tool=pmcentrez{&}rendertype=abstracthttp://linkinghub.elsevier.com/retrieve/pii/S0896627311007926.

Christian Pozzorini, Richard Naud, Skander Mensi, and Wulfram Gerstner. Temporal whitening by power-law adaptation in neocortical neurons. Nature Neuroscience, 16(7):942–948, 2013. ISSN 1097-6256. doi: 10.1038/nn.3431. URL http://www.nature.com/neuro/journal/v16/n7/full/nn.3431.html{%}5Cnhttp://www.nature.com/neuro/journal/v16/n7/extref/nn.3431-S1.pdf{%}5Cnhttp://www.nature.com/neuro/journal/v16/n7/pdf/nn.3431.pdf.

Cathy J. Price and Karl J. Friston. Degeneracy and cognitive anatomy. Trends in Cognitive Sciences, 6 (10):416–421, oct 2002. ISSN 13646613. doi: 10.1016/S1364-6613(02)01976-9. URL http://www.ncbi.nlm.nih.gov/pubmed/12413574.

Srikanth Ramaswamy, Jean-Denis Courcol, Marwan Abdellah, Stanislaw R. Adaszewski, Nicolas Antille, Selim Arsever, Guy Atenekeng, Ahmet Bilgili, Yury Brukau, Athanassia Chalimourda, Giuseppe Chindemi, Fabien Delalondre, Raphael Dumusc, Stefan Eilemann, Michael Emiel Gevaert, Padraig Gleeson, Joe W. Graham, Juan B. Hernando, Lida Kanari, Yury Katkov, Daniel Keller, James G. King, Rajnish Ranjan, Michael W. Reimann, Christian Rössert, Ying Shi, Julian C. Shillcock, Martin Telefont, Werner Van Geit, Jafet Villafranca Diaz, Richard Walker, Yun Wang, Stefano M. Zaninetta, Javier DeFelipe, Sean L. Hill, Jeffrey Muller, Idan Segev, Felix Schürmann, Eilif B. Muller, and Henry Markram. The neocortical microcircuit collaboration portal: a resource for rat somatosensory cortex. Frontiers in Neural Circuits, 9:44, oct 2015. ISSN 1662-5110. doi: 10.3389/fncir.2015.00044. URL http://journal.frontiersin.org/Article/10.3389/fncir.2015.00044/abstract.

A. Renart, J. de la Rocha, P. Bartho, L. Hollender, N. Parga, A. Reyes, and K. D. Harris. The Asynchronous State in Cortical Circuits. Science, 327(5965):587–590, jan 2010. ISSN 0036-8075. doi: 10.1126/science.1179850. URL http://www.sciencemag.org/cgi/doi/10.1126/science.1179850.

Alex Roxin. The Role of Degree Distribution in Shaping the Dynamics in Networks of Sparsely Connected Spiking Neurons. Frontiers in Computational Neuroscience, 5:8, 2011. ISSN 1662-5188. doi: 10.3389/fncom.2011.00008. URL http://journal.frontiersin.org/article/10.3389/fncom.2011.00008/abstract.

Ran Rubin, L. F. Abbott, and Haim Sompolinsky. Balanced Excitation and Inhibition are Required for High-Capacity, Noise-Robust Neuronal Selectivity. may 2017. ISSN 0027-8424. doi: 10.1073/pnas.1705841114. URL http://arxiv.org/abs/1705.01502.

T. J. H. Ruigrok, R. A. Hensbroek, and J. I. Simpson. Spontaneous Activity Signatures of Morphologically Identified Interneurons in the Vestibulocerebellum. Journal of Neuroscience, 31(2):712–724, jan 2011. ISSN 0270-6474. doi: 10.1523/JNEUROSCI.1959-10.2011. URL http://www.jneurosci.org/cgi/doi/10.1523/JNEUR0SCI.1959-10.2011.

B. L. Sabatini and W. G. Regehr. Timing of Synaptic Transmission. Annual Review of Physiology, 61(1):521–542, mar 1999. ISSN 0066-4278. doi: 10.1146/annurev.physiol.61.1.521. URL http://www.annualreviews.org/doi/10.1146/annurev.physiol.61.1.521.

Maximilian Schmidt, Rembrandt Bakker, Kelly Shen, Gleb Bezgin, Markus Diesmann, and Sacha J van Albada. Full-density multi-scale account of structure and dynamics of macaque visual cortex. 2015. URL http://arxiv.org/abs/1511.09364.

Michael A. Schwemmer, Adrienne L. Fairhall, Sophie Denéve, and Eric T. Shea-Brown. Constructing precisely computing networks with biophysical spiking neurons. The Journal of Neuroscience, 32 (28):10112–10134, 2014. ISSN 0270-6474. doi: 10.1523/JNEUROSCI.4951-14.2015. URL http://arxiv.org/abs/1411.3191.

M. N. Shadlen and W. T. Newsome. The variable discharge of cortical neurons: implications for connectivity, computation, and information coding. The Journal of neuroscience: the official journal of the Society for Neuroscience, 18(10):3870–96, may 1998. ISSN 0270-6474. doi: 0270-6474/98/183870-27$05.00/0. URL http://www.ncbi.nlm.nih.gov/pubmed/9570816.

Michael N. Shadlen and William T. Newsome. Noise, neural codes and cortical organization. Current Opinion in Neurobiology, 4(4):569–579, 1994. ISSN 09594388. doi: 10.1016/0959-4388(94)90059-0. URL http://www.sciencedirect.com/science/article/pii/0959438894900590.

Gordon M. Shepard and Sten Grillner. Handbook of Brain Microcircuits. 2010. ISBN 9780195389883. doi: 10.1093/med/9780195389883.001.0001. URL http://oxfordmedicine.com/view/10.1093/med/9780195389883.001.0001/med-9780195389883.

Masanori Shimono and John M. Beggs. Functional clusters, hubs, and communities in the cortical microconnectome. Cerebral Cortex, 25(10):3743–3757, 2015. ISSN 14602199. doi: 10.1093/cercor/bhu252. URL http://www.cercor.oxfordjournals.org/lookup/doi/10.1093/cercor/bhu252.

Shigeru Shinomoto, Keisetsu Shima, and Jun Tanji. Differences in spiking patterns among cortical neurons. Neural computation, 15(12):2823–2842, 2003. ISSN 0899-7667. doi: 10.1162/089976603322518759.

Shigeru Shinomoto, Hideaki Kim, Takeaki Shimokawa, Nanae Matsuno, Shintaro Funahashi, Keisetsu Shima, Ichiro Fujita, Hiroshi Tamura, Taijiro Doi, Kenji Kawano, Naoko Inaba, Kikuro Fukushima, Sergei Kurkin, Kiyoshi Kurata, Masato Taira, Ken Ichiro Tsutsui, Hidehiko Komatsu, Tadashi Ogawa, Kowa Koida, Jun Tanji, and Keisuke Toyama. Relating neuronal firing patterns to functional differentiation of cerebral cortex. PLoS Computational Biology, 5(7), 2009. ISSN 1553734X. doi: 10.1371/journal.pcbi.1000433.

Michael A. Silver and Sabine Kastner. Topographic maps in human frontal and parietal cortex. Trends in Cognitive Sciences, 13(11):488–495, nov 2009. ISSN 13646613. doi: 10.1016/j.tics.2009.08.005. URL http://www.pubmedcentral.nih.gov/articlerender.fcgi?artid=2767426{&}tool=pmcentrez{&}rendertype=abstract.

Wolf Singer. Complexity as Substrate for Neuronal Computations. Complexity and Analogy in Science: Theoretical, Methodological and Epistemological Aspects, 22:209–218, 2015. URL http://www.pas.va/content/dam/accademia/pdf/acta22/acta22-singer.pdf.

Sen Song, Per Jesper Sj??str??m, Markus Reigl, Sacha Nelson, and Dmitri B. Chklovskii. Highly nonrandom features of synaptic connectivity in local cortical circuits. PLoS Biology, 3(3):0507–0519, mar 2005. ISSN 15449173. doi: 10.1371/journal.pbio.0030068. URL http://www.pubmedcentral.nih.gov/articlerender.fcgi?artid=1054880&tool=pmcentrez&rendertype=abstract.

Nelson Spruston. Pyramidal neurons: dendritic structure and synaptic integration. Nature Reviews Neuroscience, 9(3):206–221, 2008. ISSN 1471-003X. doi: 10.1038/nrn2286. URL http://www.nature.com/doifinder/10.1038/nrn2286.

Armen Stepanyants and Dmitri B. Chklovskii. Neurogeometry and potential synaptic connectivity, jul 2005. ISSN 01662236. URL http://linkinghub.elsevier.com/retrieve/pii/S0166223605001311.

Thomas C. Südhof. Neurotransmitter release: The last millisecond in the life of a synaptic vesicle. Neuron, 80(3):675–690, 2013. ISSN 08966273. doi: 10.1016/j.neuron.2013.10.022.

Thomas C. Südhof and Robert C. Malenka. Understanding Synapses: Past, Present, and Future. Neuron, 60(3):469–476, nov 2008. ISSN 08966273. doi: 10.1016/j.neuron.2008.10.011. URL http://www.ncbi.nlm.nih.gov/pubmed/18995821http://www.pubmedcentral.nih.gov/articlerender.fcgi?artid=PMC3243741http://linkinghub.elsevier.com/retrieve/pii/S0896627308008842.

Susan M Sunkin, Lydia Ng, Chris Lau, Tim Dolbeare, Terri L Gilbert, Carol L Thompson, Michael Hawrylycz, and Chinh Dang. Allen Brain Atlas: an integrated spatio-temporal portal for exploring the central nervous system. Nucleic acids research, 41(Database issue):D996–D1008, jan 2013. ISSN 1362-4962. doi: 10.1093/nar/gks1042. URL http://www.ncbi.nlm.nih.gov/pubmed/23193282http://www.pubmedcentral.nih.gov/articlerender.fcgi?artid=PMC3531093.

J. Szabadics. Excitatory Effect of GABAergic Axo-Axonic Cells in Cortical Microcircuits. Science, 311(5758):233–235, aug 2006. ISSN 0036-8075. doi: 10.1126/science.1121325. URL http://www.sciencemag.org/cgi/doi/10.1126/science.1121325.

Naoya Takahashi, Kazuo Kitamura, Naoki Matsuo, Mark Mayford, Masanobu Kano, Norio Matsuki, and Yuji Ikegaya. Locally Synchronized Synaptic Inputs. Science, 335(6066):353–356, jan 2012. ISSN 0036-8075,1095-9203. doi: 10.1126/science.1210362. URL http://www.sciencemag.org/content/335/6066/353{%}5Cnhttp://www.sciencemag.org/content/335/6066/353.full.pdf.

Christian Tetzlaff, Christoph Kolodziejski, Irene Markelic, and Florentin W??rg??tter. Time scales of memory, learning, and plasticity. Biological Cybernetics, 106(11-12):715–726, dec 2012. ISSN 14320770. doi: 10.1007/s00422-012-0529-z. URL http://link.springer.com/10.1007/s00422-012-0529-z.

Dominik Thalmeier, Marvin Uhlmann, Hilbert J. Kappen, and Raoul Martin Memmesheimer. Learning Universal Computations with Spikes. PLoS Computational Biology, 12(6):1–29, 2016. ISSN 15537358. doi: 10.1371/journal.pcbi.1004895.

Jean Philippe Thivierge and Gary F. Marcus. The topographic brain: from neural connectivity to cognition. Trends in Neurosciences, 30(6):251–259, jun 2007. ISSN 01662236. doi: 10.1016/j.tins.2007.04.004. URL http://www.ncbi.nlm.nih.gov/pubmed/17462748.

A. M. Thomson. Synaptic Connections and Small Circuits Involving Excitatory and Inhibitory Neurons in Layers 2-5 of Adult Rat and Cat Neocortex: Triple Intracellular Recordings and Biocytin Labelling In Vitro. Cerebral Cortex, 12(9):936–953, sep 2002. ISSN 14602199. doi: 10.1093/cercor/12.9.936. URL https://academic.oup.com/cercor/article-lookup/doi/10.1093/cercor/12.9.936.

Christian Tomm. Analysing Neuronal Network Architectures: From Weight Distributions to Structure and Back. PhD thesis, ECOLE POLYTECHNIQUE FÉDÉRALE DE LAUSANNE, 2012. URL https://infoscience.epfl.ch/record/174669/files/EPFL{_}TH5302.pdf.

Christian Tomm, Michael Avermann, C. Petersen, Wulfram Gerstner, and Tim P Vogels. Connection-type-specific biases make uniform random network models consistent with cortical recordings. Journal of Neurophysiology, 112(8):1801–1814, 2014. ISSN 0022-3077. doi: 10.1152/jn.00629.2013. URL http://classic.jn.physiology.org/content/early/2014/06/13/jn.00629.2013.abstract{%}5Cnhttp://jn.physiology.org/cgi/doi/10.1152/jn.00629.2013.

Shreejoy J. Tripathy, Judith Savitskaya, Shawn D. Burton, Nathaniel N. Urban, and Richard C. Gerkin. NeuroElectro: a window to the world’s neuron electrophysiology data. Frontiers in Neuroinformatics, 8 (April):40, 2014. ISSN 1662-5196. doi: 10.3389/fninf.2014.00040. URL http://journal.frontiersin.org/article/10.3389/fninf.2014.00040/abstract.

Shreejoy J. Tripathy, Shawn D. Burton, Matthew Geramita, Richard C. Gerkin, and Nathaniel N. Urban. Brain-wide analysis of electrophysiological diversity yields novel categorization of mammalian neuron types. Journal of Neurophysiology, 113(10):3474–3489, 2015. ISSN 0022-3077. doi: 10.1152/jn.00237.2015. URL http://jn.physiology.org/lookup/doi/10.1152/jn.00237.2015.

M Tsodyks and T Sejnowski. Rapid state switching in balanced cortical network models. Network: Computation in Neural Systems, 6(2):111–124, 1995. ISSN 0954-898X. doi: 10.1088/0954-898X/6/2/001. URL http://www.informaworld.com/openurl?genre=article&doi=10.1088/0954-898X/6/2/001&magic=crossref{%}7C{%}7CD404A21C5BB053405B1A640AFFD44AE3.

J K Tsotsos, S M Culhane, W Y K Wai, Yuzhong Lai, N Davis, and F Nuflo. Dynamics of Sparsely Conntected Networks of Excitatory and Inhibitory Spiking Neurons. Journal of Computational Neuroscience, 8:183–208, 2000.

Gina G. Turrigiano. Homeostatic plasticity in neuronal networks: The more things change, the more they stay the same. Trends in Neurosciences, 22(5):221–227, may 1999. ISSN 01662236. doi: 10.1016/S0166-2236(98)01341-1. URL http://www.ncbi.nlm.nih.gov/pubmed/10322495.

Karlijn I. Van Aerde and Dirk Feldmeyer. Morphological and physiological characterization of pyramidal neuron subtypes in rat medial prefrontal cortex. Cerebral Cortex, 25(3):788–805, mar 2015. ISSN 14602199. doi: 10.1093/cercor/bht278. URL http://www.ncbi.nlm.nih.gov/pubmed/24108807https://academic.oup.com/cercor/article-lookup/doi/10.1093/cercor/bht278.

David van den Broek, Marvin Uhlmann, Hartmut Fitz, Renato Duarte, Peter Hagoort, and Karl Magnus Petersson. The best spike filter kernel is a neuron. In Cognitive Computational Neuroscience, volume 1, 2017.

Gert van Dijck, Marc M. van Hulle, Shane A. Heiney, Pablo M. Blazquez, Hui Meng, Dora E. Angelaki, Alexander Arenz, Troy W. Margrie, Abteen Mostofi, Steve Edgley, Fredrik Bengtsson, Carl Fredrik Ekerot, Henrik Jürntell, Jeffrey W. Dalley, and Tahl Holtzman. Probabilistic Identification of Cerebellar Cortical Neurones across Species. PLoS ONE, 8(3), 2013. ISSN 19326203. doi: 10.1371/journal.pone.0057669.

C van Vreeswijk and H Sompolinsky. Chaos in neuronal networks with balanced excitatory and inhibitory activity. Science (New York, N.Y.), 274(5293):1724–6, dec 1996. ISSN 0036-8075. URL http://www.ncbi.nlm.nih.gov/pubmed/8939866.

C. van Vreeswijk and H. Sompolinsky. Chaotic Balanced State in a Model of Cortical Circuits. Neural Computation, 10(6):1321–1371, aug 1998. ISSN 0899-7667. doi: 10.1162/089976698300017214. URL http://www.mitpressjournals.org/doi/10.1162/089976698300017214.

David C. VanEssen. Cartography and connectomes. Neuron, 80(3):775–790, 2013. ISSN 08966273. doi: 10.1016/j.neuron.2013.10.027. URL http://dx.doi.org/10.1016/j.neuron.2013.10.027.

T. P. Vogels. Signal Propagation and Logic Gating in Networks of Integrate-and-Fire Neurons. Journal of Neuroscience, 25(46):10786–10795, 2005. ISSN 0270-6474. doi: 10.1523/JNEUROSCI.3508-05.2005. URL http://www.jneurosci.org/cgi/doi/10.1523/JNEUR0SCI.3508-05.2005.

T. P. Vogels, H. Sprekeler, F. Zenke, C. Clopath, and W. Gerstner. Inhibitory Plasticity Balances Excitation and Inhibition in Sensory Pathways and Memory Networks. Science, 334(6062):1569–1573, 2011. ISSN 0036-8075. doi: 10.1126/science.1211095. URL http://www.sciencemag.org/cgi/doi/10.1126/science.1211095.

Tim P Vogels and L F Abbott. Gating multiple signals through detailed balance of excitation and inhibition in spiking networks. Nature Neuroscience, 12(4):483–491, apr 2009. ISSN 1097-6256. doi: 10.1038/nn.2276. URL http://www.nature.com/doifinder/10.1038/nn.2276.

Tim P. Vogels, Kanaka Rajan, and L.F. Abbott. Neural Network Dynamics. Annual Review of Neuroscience, 28(1):357–376, jan 2005. ISSN 0147-006X. doi: 10.1146/annurev.neuro.28.061604.135637. URL http://www.annualreviews.org/doi/10.1146/annurev.neuro.28.061604.135637.

Giannis Voglis and Nektarios Tavernarakis. The role of synaptic ion channels in synaptic plasticity. EMBO reports, 7(11):1104–1110, 2006. ISSN 1469-221X. doi: 10.1038/sj.embor.7400830. URL http://embor.embopress.org/cgi/doi/10.1038/sj.embor.7400830.

Konrad Wagstyl, Lisa Ronan, Ian M. Goodyer, and Paul C. Fletcher. Cortical thickness gradients in structural hierarchies. NeuroImage, 111:241–250, 2015. ISSN 10959572. doi: 10.1016/j.neuroimage.2015.02.36. URL http://dx.doi.org/10.1016/j.neuroimage.2015.02.036.

J. Waters and F. Helmchen. Background Synaptic Activity Is Sparse in Neocortex. Journal of Neuroscience, 26(32):8267–8277, aug 2006. ISSN 0270-6474. doi: 10.1523/JNEUROSCI.2152-06.2006. URL http://www.jneurosci.org/cgi/doi/10.1523/JNEUR0SCI.2152-06.2006.

Alanna J. Watt, Mark C.W. Van Rossum, Katrina M. MacLeod, Sacha B. Nelson, and Gina G. Turrigiano. Activity coregulates quantal AMPA and NMDA currents at neocortical synapses. Neuron, 26(3): 659–670, 2000. ISSN 08966273. doi: 10.1016/S0896-6273(00)81202-7.

Philipp Weidel, Mikael Djurfeldt, Renato Duarte, and Abigail Morrison. Closed loop interactions between spiking neural network and robotic simulators based on MUSIC and ROS. Frontiers in Neuroinformatics, 10(31):1–19, 2016. ISSN 1662-5196. doi: 10.3389/fninf.2016.00031. URL http://arxiv.org/abs/1604.04764{%}0Ahttp://dx.doi.org/10.3389/fninf.2016.00031.

A. Wertz, S. Trenholm, K. Yonehara, D. Hillier, Z. Raics, M. Leinweber, G. Szalay, A. Ghanem, G. Keller, B. Rozsa, K.-K. Conzelmann, and B. Roska. Single-cell-initiated monosynaptic tracing reveals layer-specific cortical network modules. Science, 349(6243):70–74, jul 2015. ISSN 0036-8075. doi: 10.1126/science.aab1687. URL http://dx.doi.org/10.3389/fninf.2016.00031.

Nathan R. Wilson, Caroline A. Runyan, Forea L. Wang, and Mriganka Sur. Division and subtraction by distinct cortical inhibitory networks in vivo. Nature, 488(7411):343–348, aug 2012. ISSN 0028-0836. doi: 10.1038/nature11347. URL http://www.nature.com/doifinder/10.1038/nature11347.

B. T. Thomas Yeo, Fenna M. Krienen, Jorge Sepulcre, Mert R. Sabuncu, D. Lashkari, Marisa Hollinshead, Joshua L. Roffman, Jordan W. Smoller, Lilla Zollei, Jonathan R. Polimeni, Bruce Fischl, Hesheng Liu, and Randy L. Buckner. The organization of the human cerebral cortex estimated by intrinsic functional connectivity. Journal of neurophysiology, 106:1125–1165, 2011. ISSN 1522-1598. doi: 10.1152/jn.00338.2011. URL http://jn.physiology.org/content/106/3/1125.short.

Ofer Yizhar, Lief E. Fenno, Matthias Prigge, Franziska Schneider, Thomas J. Davidson, Daniel J. O’Shea, Vikaas S. Sohal, Inbal Goshen, Joel Finkelstein, Jeanne T. Paz, Katja Stehfest, Roman Fudim, Charu Ramakrishnan, John R. Huguenard, Peter Hegemann, and Karl Deisseroth. Neocortical excitation/inhibition balance in information processing and social dysfunction. Nature, 477(7363): 171–178, sep 2011. ISSN 0028-0836. doi: 10.1038/nature10360. URL http://www.nature.com/doifinder/10.1038/nature10360.

Yumiko Yoshimura and Edward M Callaway. Fine-scale specificity of cortical networks depends on inhibitory cell type and connectivity. Nature Neuroscience, 8(11):1552–1559, nov 2005. ISSN 1097-6256. doi: 10.1038/nn1565. URL http://www.nature.com/doifinder/10.1038/nn1565.

Yumiko Yoshimura, Jami L. M. Dantzker, and Edward M. Callaway. Excitatory cortical neurons form fine-scale functional networks. Nature, 433(7028):868–873, feb 2005. ISSN 0028-0836. doi: 10.1038/nature03252. URL http://www.nature.com/doifinder/10.1038/nature03252.

A. V. Zaitsev, N. V. Povysheva, G. Gonzalez-Burgos, and D. A. Lewis. Electrophysiological classes of layer 2/3 pyramidal cells in monkey prefrontal cortex. Journal of Neurophysiology, 108(2):595–609, 2012. ISSN 0022-3077. doi: 10.1152/jn.00859.2011. URL http://jn.physiology.org/cgi/doi/10.1152/jn.00859.2011.

Lyuba Zehl, Florent Jaillet, Adrian Stoewer, Jan Grewe, Andrey Sobolev, Thomas Wachtler, Thomas G. Brochier, Alexa Riehle, Michael Denker, and Sonja Grün. Handling Metadata in a Neurophysiology Laboratory. Frontiers in Neuroinformatics, 10:26, jul 2016. ISSN 1662-5196. doi: 10.3389/fninf.2016.00026. URL http://journal.frontiersin.org/Article/10.3389/fninf.2016.00026/abstract.

Friedemann Zenke and Wulfram Gerstner. Cooperation across timescales between Hebbian and homeostatic plasticity. Philosophical Transactions of the Royal Society B: Biological Sciences, ((In Review)), 2016.

K. Zilles, N. Palomero-Gallagher, C. Grefkes, F. Scheperjans, C. Boy, K. Amunts, and A. Schleicher. Architectonics of the human cerebral cortex and transmitter receptor fingerprints: Reconciling functional neuroanatomy and neurochemistry. European Neuropsychopharmacology, 12(6):587–599, 2002a. ISSN 0924977X. doi: 10.1016/S0924-977X(02)00108-6.

Karl Zilles and Katrin Amunts. Receptor mapping: architecture of the human cerebral cortex. Current Opinion in Neurology, 22(4):331–339, aug 2009. ISSN 1350-7540. doi: 10.1097/WCO.0b013e32832d95db. URL http://content.wkhealth.com/linkback/openurl?sid=WKPTLP:landingpage&an=00019052-200908000-00002.

Karl Zilles, Axel Schleicher, Nicola Palomero-Gallagher, and Katrin Amunts. Quantitative Analysis of Cyto- and Receptor Architecture of the Human Brain, volume 58. 2002b. ISBN 0385-5600 (Print) 0385-5600 (Linking). doi: 10.1016/B978-012693019-1/50023-X. URL http://www.epjap.org/10.1051/epjap/2012110475{%}5Cnhttp://linkinghub.elsevier.com/retrieve/pii/B978012693019150023X.

Karl Zilles, Nicola Palomero-Gallagher, and Axel Schleicher. Transmitter receptors and functional anatomy of the cerebral cortex. Journal of Anatomy, 205(6):417–432, 2004. ISSN 00218782. doi: 10.1111/j.0021-8782.2004.00357.x. URL http://eutils.ncbi.nlm.nih.gov/entrez/eutils/elink.fcgi?dbfrom=pubmed&id=15610391&retmode=ref&cmd=prlinks{%}5Cnfile:///Users/balarampooja/Library/ApplicationSupport/Papers2/Articles/2004/Zilles/JournalofAnatomy2004Zilles.pdf%5Cnpapers2://publication/doi/10.

Stefano Zucca, Giulia D’Urso, Valentina Pasquale, Dania Vecchia, Giuseppe Pica, Serena Bovetti, Claudio Moretti, Stefano Varani, Manuel Molano-Mazón, Michela Chiappalone, Stefano Panzeri, and Tommaso Fellin. An inhibitory gate for state transition in cortex. eLife, 6:e26177, may 2017. ISSN 2050084X. doi: 10.7554/eLife.26177.001. URL http://elifesciences.org/lookup/doi/10.7554/eLife.26177.

