## Supplementary Materials for "Leveraging heterogeneity for neural computation with fading memory in layer 2/3 cortical microcircuits"

Renato Duarte[1,2,3,4,*], Abigail Morrison[1,2,3,5]

[1] Institute of Neuroscience and Medicine (INM-6) and Institute for Advanced Simulation (IAS-6) and JARA BRAIN Institute I, Jülich Research Centre, Jülich, Germany
[2] Bernstein Center Freiburg, Albert-Ludwig University of Freiburg, Germany
[3] Faculty of Biology, Albert-Ludwig University of Freiburg, Freiburg im Breisgau, Germany
[4] Institute of Adaptive and Neural Computation, School of Informatics, University of Edinburgh, UK
[5] Institute of Cognitive Neuroscience, Faculty of Psychology, Ruhr-University Bochum, Bochum, Germany

*

### 1 Supplementary tables

| A: Model Summary | |
|---|---|
| **Populations** | 1 Excitatory, 2 Inhibitory |
| **Topology** | None |
| **Connectivity** | Sparse, random or structured ($\text{HET}_1$), with density $p^{\text{syn}}$ |
| **Neuron Model** | Adaptive leaky integrate-and-fire, fixed voltage threshold, fixed absolute refractory time, sub-threshold and spike-triggered adaptation |
| **Synapse Model** | Multi-receptor kinetics |
| **Plasticity** | None |
| **Input** | Stochastic background spikes and somatic current injection onto 25% E neurons |
| **Measurements** | Spiking activity, membrane potentials, synaptic currents/conductances |

| B: Populations | | |
|---|---|---|
| **Name** | **Elements** | **Size** |
| E | `iaf_cond_mtime` | 2000 |
| $I_1$ | `iaf_cond_mtime` | 175 |
| $I_2$ | `iaf_cond_mtime` | 325 |

| C: Neuron Models | |
|---|---|
| **Name** | Multi-adaptive integrate-and-fire neuron (`iaf_cond_mtime`) |
| **Subthreshold Dynamics** | if $\left(t > t^f + \tau_{\text{ref}}\right)$ <br> $C_m \frac{dV_i}{dt} = -\, g_{\text{leak}} \left(V_i(t) - E_L\right) - I_{i,\text{adapt}}(t) - \sum_{k\in\text{ syn}} \sum_{j\in\text{pre}} I^k_{ij}(t)$ <br> else <br> $V(t) = V_{\text{reset}}$ |
| **Intrinsic Adaptation** | $\tau_w \frac{dI_{i,\text{adapt}}}{dt} = -I_{i,\text{adapt}} + a\left(I_{i,\text{adapt}} - V_i\right) + b\sum_{t_f\in F(i)} \delta\left(t - t_f\right)$ |
| **Synaptic Transmission** | $I^{\text{syn}}_{\text{ij}}(t) = g^{\text{syn}}_{\text{ij}}(t, V_i)\left(V_i(t) - E_{\text{syn}}\right)$ |
| **Spiking** | If $V(t-) < V_{th}$ OR $V(t+) \geq V_{th}$ <br> 1. set $t^f = t$ 2. emit spike with time stamp $t^f$ |

| D: Synapse Models | |
|---|---|
| **Receptor Conductance** | $g^{\text{rec}}_{\text{ij}}(t, V_i) = \bar{g}_{\text{rec}} n_{\text{rec}}(V) w^{\text{rec}}_{\text{ij}}(t) \sum_{t_f\in F(j)} \left[1 - \exp\left(-\frac{t - t_f}{\tau^{\text{rec}}_{\text{rise}}}\right)\right] \left[r_{\text{rec}} \exp\left(-\frac{t - t_f}{\tau^{\text{rec}}_{\text{decay}_\text{f}}}\right) + (1 - r_{\text{rec}}) \exp\left(-\frac{t - t_f}{\tau^{\text{rec}}_{\text{decay}_\text{s}}}\right)\right] \Theta\left(t - t_f\right)$ |

| E: Input | | |
|---|---|---|
| **Type** | **Target** | **Description** |
| `poisson_generator` | [E, $I_1$, $I_2$] | Total rate $\nu_{\text{in}} K_{\text{in}}$ |
| `step_current_generator` | E | Step current amplitude $u[n]\rho_u$, changing every $\Delta t$ ms |

| F: Measurements |
|---|
| Spiking activity, membrane potentials, synaptic currents and conductances |

**Supplementary Table 1.** Tabular description of network model after [Nordlie et al., 2009].

| Parameter | Data | Model | | Description |
|---|---|---|---|---|
| | (*) | $\bar{x}$ | s | |
| $I_{rh}$[pA] | $[190, 250]$ | 262.73 | 45.88 | rheobase current |
| | $[190, 250]$ | 251.5 | 44.8 | |
| | $[80, 150]$ | 172.16 | 19.8 | |
| | $E \approx I_1 >> I_2$ | $E \approx I_1 >> I_2$ | | |
| Slope[Hz/pA] | $[140, 250]$ | 226.79 | 36.1 | slope of the fI curve |
| | $[240, 500]$ | 529.46 | 252.16 | |
| | $[150, 400]$ | 305.82 | 76.21 | |
| | $I_1 > I_2 > E$ | $I_1 > I_2 > E$ | | |
| $\nu_{min}$[Hz] | $[0.5, 10]$ | 4.84 | 2.7 | minimum firing rate |
| | $[15, 30]$ | 15.57 | 3.3 | |
| | $[5, 20]$ | 8.23 | 3.42 | |
| | $I_1 > I_2 > E$ | $I_1 > I_2 > E$ | | |
| $\nu_{max}$[Hz] | $[15, 80]$ | 130.83 | 21.31 | maximum firing rate |
| | $[180, 250]$ | 330.28 | 164.93 | |
| | $[40, 200]$ | 208.69 | 38.8 | |
| | $I_1 > I_2 > E$ | $I_1 > I_2 > E$ | | |

**Supplementary Table 2.** Discrepancies between electrophysiological parameters reported in the literature and model results. The results obtained after careful choice of the individual parameters for the different neuronal classes did not exactly match the experimental reports, but the relative relations between classes are retained. (*) Note that the ranges reported in this table are a rough approximation to the range of mean values reported in different studies (see below). Naturally, values like the maximum rate ($\nu_{max}$[Hz]) depend entirely on the range of input current considered in a given experiment, so in this case, only the relative ratio is pertinent.

#### 2 Primary data sources

Primary literature sources considered and used to constrain and guide the choices for all the different parameter sets. Note that most of the sources for the neuronal parameters correspond to individual data points included in the NeuroElectro [Tripathy et al., 2014, Tripathy et al., 2015] database. Additionally, it is worth mentioning that, apart from direct experimental measurements, we also included the values of some parameters used in previous biophysical and phenomenological models, provided they accounted for the relevant phenomena.

- Neurons:

  [Avermann et al., 2012, Harrison et al., 2015, Lu et al., 2007, Neske et al., 2015, Pi et al., 2013, Helmstaedter et al., 2009, Kröner et al., 2007, Levy and Reyes, 2012, Tyler et al., 2015, Lacroix et al., 2015, Cruikshank et al., 2012, Zaitsev et al., 2012, Cho et al., 2010, Yang et al., 2013b, Oswald and Reyes, 2008, van der Velden et al., 2012, Yang et al., 2013a, Povysheva et al., 2006, Valdés-Sánchez et al., 2007, Gonzalez-Burgos et al., 2015, González-Burgos et al., 2004, Tateno, 2004, Waters and Helmchen, 2006, Sippy and Yuste, 2013, Pillai et al., 2014, Dong et al., 2004, Yang et al., 2013b, Sessler et al., 1998, Lee et al., 2005, Nowak, 2002, Yang and Benardo, 1997, Guan et al., 2007, Barnes et al., 2015, Brill and Huguenard, 2010, Cummings et al., 2009, Elstrott et al., 2014, Aramakis et al., 2000, Shruti et al., 2008, Kobayashi et al., 2008, Luebke et al., 2015, Ren et al., 2014, Waters and Helmchen, 2006, González-Burgos et al., 2004, Tateno, 2004, Lemtiri-Chlieh and Levine, 2007, Luhmann et al., 1998, Krimer, 2005, Wayman et al., 2015, Andjelic et al., 2008, Castro et al., 2002, Amatrudo et al., 2012, Takei et al., 2010, Huggenberger et al., 2009, Tanaka et al., 2011, Gulledge et al., 2009, Oswald and Reyes, 2008, Van Aerde and Feldmeyer, 2015, Sutor et al., 2000, Tyler et al., 2015, Hadjilambreva, 2004, Beggs et al., 2000, Abel, 2003, Cheetham et al., 2007, Lambe et al., 2000, Hirai et al., 2012, Chen et al., 2015, Pineda et al., 1998, Salling et al., 2014, Cho et al., 2010, Lee et al., 2007, Mowery et al., 2015, Telfeian and Connors, 2003, Wang and Gao, 2010, Blatow et al., 2003, Takesian et al., 2012, Zhou and Roper, 2011, Rose, 2005, Aerde et al., 2015, Schiff and Reyes, 2012, Yang et al., 2013a, Oswald and Reyes, 2011, Zhong and Yan, 2016, Gao et al., 2003, Imbrosci et al., 2014, Erisir et al., 1999, Gonzalez-Burgos, 2004, Karayannis et al., 2007, Tateno and Robinson, 2008, Gorelova, 2002, Zaitsev et al., 2009, Tateno and Robinson, 2008, Dougherty et al., 2014, Abe et al., 2011, Kawaguchi and Kubota, 1998, Povysheva et al., 2008, Miyoshi et al., 2007, Akgul and Wollmuth, 2013, Cho et al., 2010, Akgul and Wollmuth, 2013, Chadderton et al., 2009, Szabadics, 2006]

- Receptors:

  [Destexhe et al., 1994, Destexhe et al., 1998, Hestrin, 1993, McCormick et al., 1993, Angulo et al., 1999, Myme, 2003, Watt et al., 2000, Hoffmann et al., 2015, Connors et al., 1988, Hill et al., 2000, Paoletti et al., 2013, Wang and Gao, 2010, Jahr and Stevens, 1990, Hestrin et al., 1990, Moreau and Kullmann, 2013, Nissen et al., 2010, Izhikevich et al., 2004, Izhikevich and Edelman, 2008, Dayan and Abbott, 2001, Gerstner and Kistler, 2002, Gerstner et al., 2014, Brunel and Wang, 2001, Destexhe and Sejnowski, 1995, McCormick et al., 1993, Mason et al., 1991, Southan et al., 2016, Sah et al., 1990, Uzun et al., 2010, Köhr et al., 1993]

- Synapses:

  [Avermann et al., 2012, Lu et al., 2007, Feldmeyer et al., 2006, Hoffmann et al., 2015, Thomson, 2002, Monier et al., 2008, Kubota et al., 2015, Lefort et al., 2009, Helmstaedter et al., 2009, Helmstaedter et al., 2008, Gonzalez-Burgos et al., 2015, Povysheva et al., 2006, Valdés-Sánchez et al., 2007]

- Connectivity:

  [Tomm, 2012, Tomm et al., 2014, Koulakov et al., 2009, Avermann et al., 2012, Lu et al., 2007, Lefort et al., 2009, Fino and Yuste, 2011, Levy and Reyes, 2012, Feldmeyer et al., 2006, Helmstaedter et al., 2009, Helmstaedter et al., 2008, Song et al., 2005, Thomson, 2002, Perin et al., 2011, Yoshimura et al., 2005, Yoshimura and Callaway, 2005, Potjans and Diesmann, 2014, Kubota et al., 2015, Hagen et al., 2016]

### 3 Supplementary figures

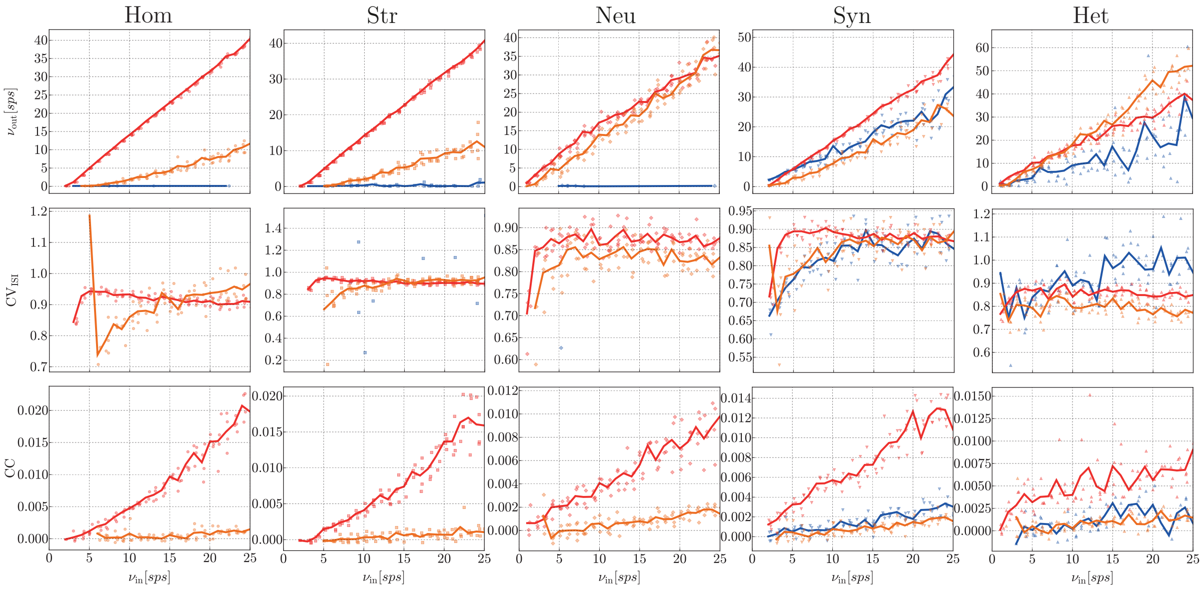


**Supplementary Figure 1.** Characteristics of population spiking activity in response to background, Poissonian input (quiet state) in the various conditions analysed and for the 3 different population types (E, blue; $\text{I}_1$, red; $\text{I}_2$, orange), as a function of the input rate $\nu_{\text{in}}$ for a total simulation time of 10 seconds. The top row corresponds the population rate transfer functions, showing that E neurons fire extremely sparsely and synaptic heterogeneity is strictly required to obtain an active E population. The middle and bottom row depict the measured irregularity ($\text{CV}_{\text{ISI}}$) and synchrony (CC) in all conditions analysed. Note that in many conditions the spiking activity in the E population is so sparse that it is not possible to compute these metrics, since the total number of spikes is insufficient.

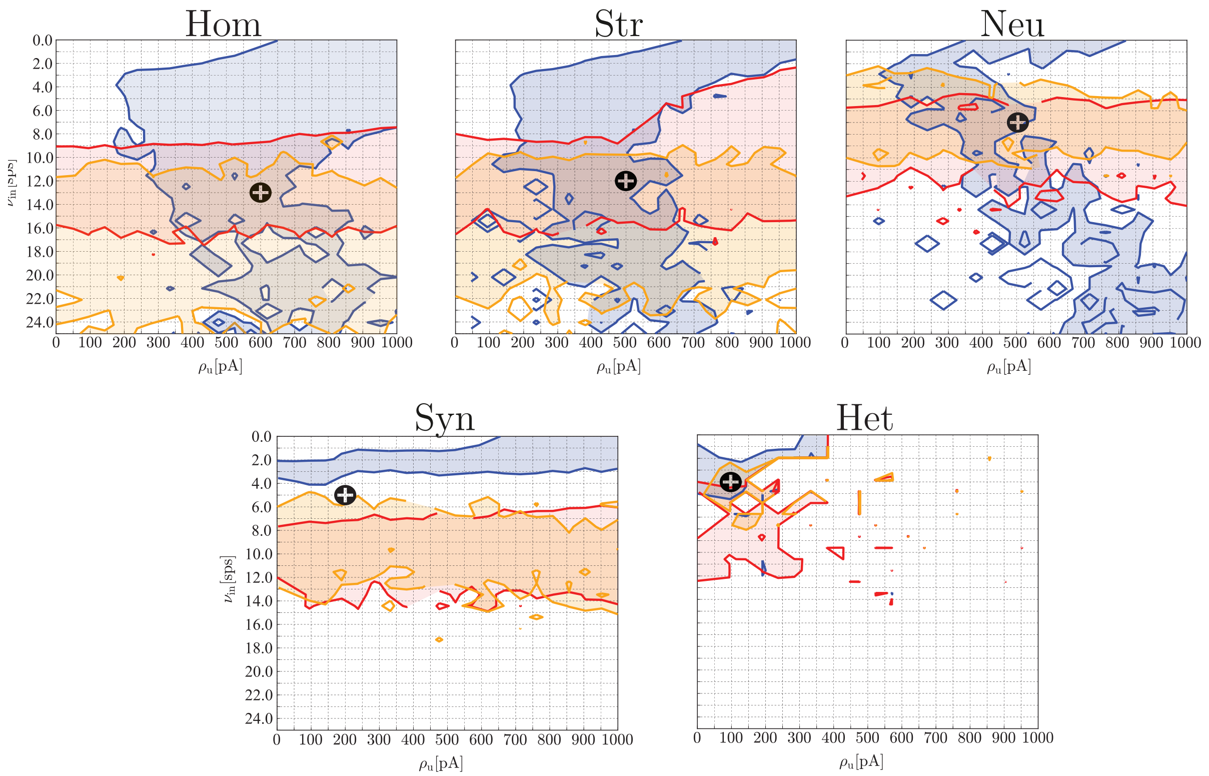


**Supplementary Figure 2.** Tuning the input parameters for the active state. To emulate an active processing condition, an extra input current of maximum amplitude $\rho_u$ is given to a randomly chosen subset of 25% excitatory neurons. The circuits in the different conditions exhibit different degrees of sensitivity to their inputs. To achieve adequate and comparable responses, we attempt to find combination of input parameters that allows the mean firing rates to remain within realistic bounds ($\nu_E \in [0.5, 5]$, $\nu_{I_1} \in [10, 25]$, $\nu_{I_2} \in [3, 15]$).

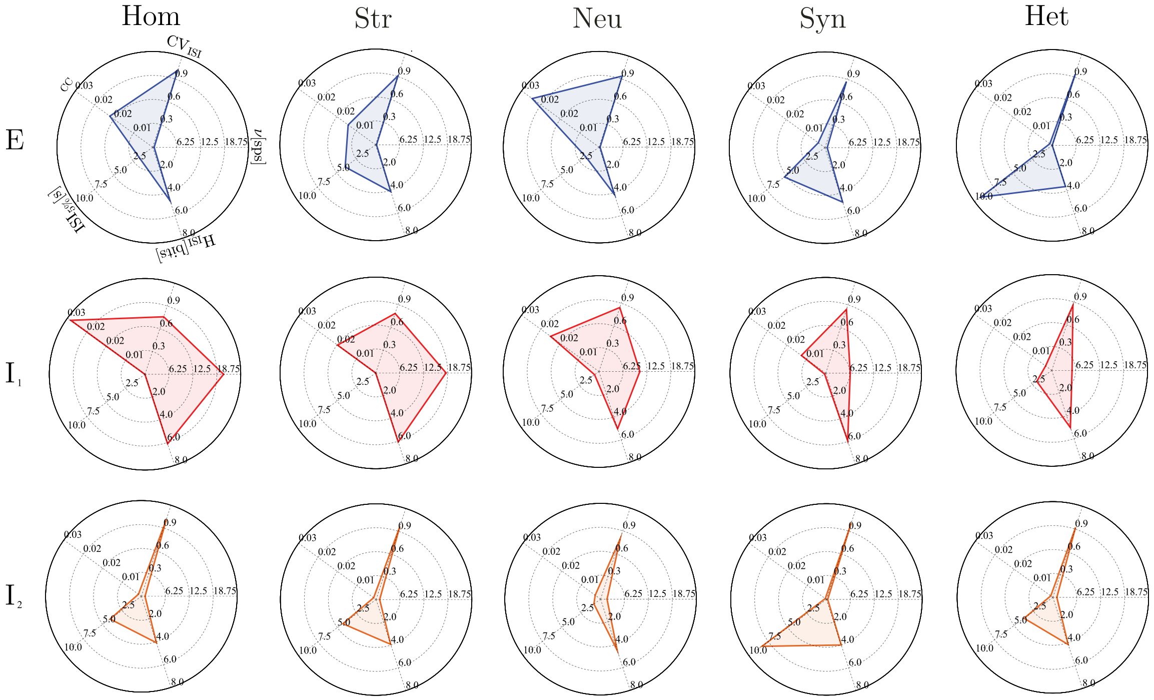


**Supplementary Figure 3.** Complete statistics of population spiking activity in the active state for the different neuron classes: E (top, blue), $I_1$ (middle, red) and $I_2$ (bottom, orange) and for the different conditions (columns). The radial axes in each plot correspond to: regularity ($CV_{ISI}$), synchrony (CC), burstiness ($ISI_{5\%}$), entropy of the ISI distribution ($H_{ISI}$), and the mean firing rate ($\nu$). All statistics were computed for an observation period of 10s, in a single realization for each condition, with all input parameters fixed and set to the values determined in Supplementary Figure 2.

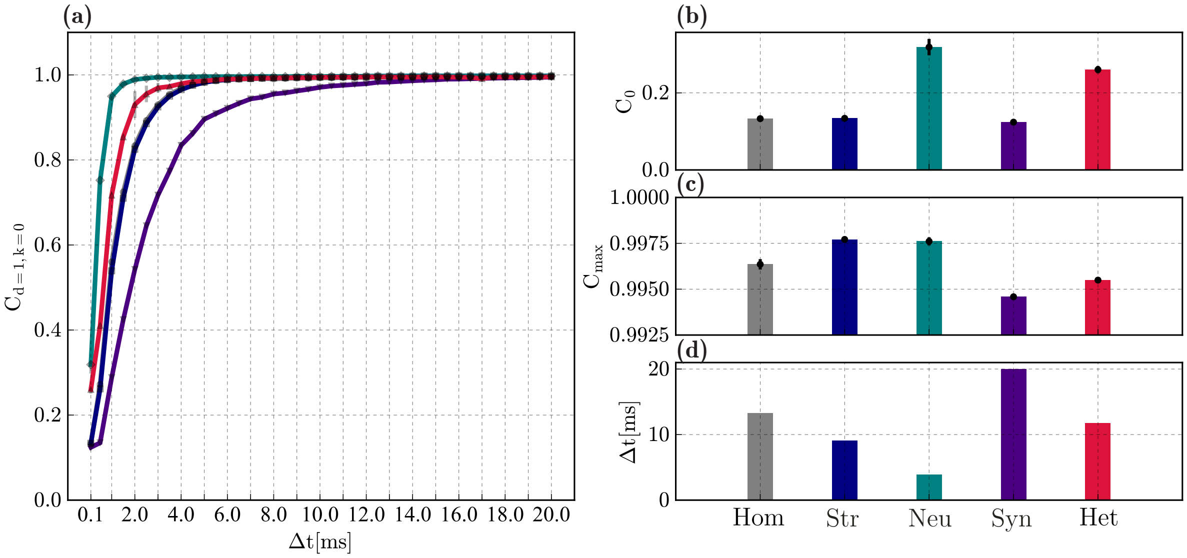


**Supplementary Figure 4.** Temporal receptivity of the microcircuits analysed. **(a)** capacity to reconstruct the original input signal at zero lag (i.e. maximum polynomial degree $d = 1$, maximum delay $k = 0$, $C_{d=1,k=0}$) as a function of the signal resolution ($\Delta t$). Since the capacity values converge asymptotically to 1, we determine the optimal resolution as the minimum $\Delta t$ at which $C_{d=1,k=0} \geq 0.99$. **(b)** Decoding capacity at minimum resolution $\Delta t = 0.1$ ms (equal to the simulation resolution). **(c)** Capacity at the maximum resolution tested ($\Delta t = 20$ ms). **(d)** Optimal resolution for each condition. All results correspond to the mean and standard deviations for 10 simulations per condition.

### 4 Reproducibility and Replication

Considerable efforts were taken to ensure transparency, openness and reproducibility. We provide all the relevant data, materials and code, necessary to replicate and scrutinize all the results presented in this manuscript.

#### 4.1 OSF Project

Through the Open Science Framework (osf.io), an open-access, curated and registered repository is openly available at [Duarte and Morrison, 2017][1]. The project contains all the relevant information necessary to replicate and scrutinize the present work, divided into the following components:

- Data - linked to the Sciebo campus cloud [Vogl et al., 2015], where the data is hosted. Due to the large size of the data (totalling $\approx 550GB$), we do not provide a complete data package as a supplement. If there are any difficulties accessing the data through this OSF component, please contact the authors.
- Figures - all the figures in the manuscript, in various formats (hosted on Sciebo as well)
- Presentations - public presentations of the project
- Software - all dedicated and modified software that is necessary to run and replicate the experiments (see below):
  - NEST 2.10.0 modified
  - NMSAT v0.2
  - Project Code
- Manuscript - all the relevant manuscript documents and files (hosted directly on OSF).
- Bibliography - linked to the Mendeley database containing all the references used in this manuscript

#### 4.2 Software and source code

The code package provided as a supplement (Supplementary File 1[2], also available in the OSF repository) implements project-specific functionality to NMSAT [Duarte et al., 2017], which is a tailor-made python package that provides a generic set of tools to build, simulate and analyse neuronal microcircuit models with any degree of complexity, as exemplified in this study. It provides a high-level wrapper for PyNEST (used as the core simulation engine). The specificities of this project require the installation and use of a specific, modified version of NEST 2.10.0 [Bos et al., 2015] (available in the Software component of OSF or upon request), since it relies on 2 models that are not currently available in the main release (`iaf_cond_mtime`: neuron model with complex synaptic kinetics, `multiport_synapse`: connection model allowing the spike-triggered conductances onto the different receptors of a given synapse type). To use the provided software:

1. Setup - After ensuring that all dependencies are satisfied, NMSAT[3] needs to be downloaded and setup, as explained in the provided documentation[4].
2. Project code - The code package for this project should then be extracted onto the `projects/` folder. The provided code has the following structure:
   `heterogeneity_project/`

[1] The project is embargoed and only accessible to reviewers, until the manuscript is accepted for publication
[2] will only be provided after the manuscript is accepted
[3] https://github.com/rcfduarte/nmsat
[4] https://rcfduarte.github.io/nmsat/

```
├── parameters/
│   └── preset/
├── computations/
├── scripts/
└── read_data/
```

where `read_data` contains all the analysis scripts necessary to read, analyse and plot the data (see Supplementary Table 4); `scripts` contains all the main experiments as a complete script, mostly for debugging purposes. The main simulations are run using combinations of parameters files with the corresponding computation function (see Supplementary Table 3 for a description of the experiments provided and the standard use case in the code documentation[5] for instructions).

3. Running a simulation - Specific experiments can be run from scratch using the provided code. Modify the specific parameters as desired (paying particular attention to the system specificities) and execute the experiment:

```
$ python main.py -f {parameters_file} -c {computation} --extra {computation_parameters}
```

4. Replicating an experiment - Alternatively, experiments can be re-run using the original parameter file (`{data_label}_ParameterSpace.py` file, see Supplementary Table 4) when executing the experiment. Simply execute as above, replacing `{parameters_file}` with the full path to the stored parameters. Note, however, that the system parameters need to be edited in these cases, as they were executed on different machines.

| **Experiment** | **Parameters file** | **Computation** |
|---|---|---|
| Single neuron fI curves and related parameters and distributions (Fig.1) | `single_neuron_fI` | `singleneuron_dcinput` |
| Single receptor kinetics (Fig. 2b) | `synaptic_response_receptors` | `synaptic_response` |
| PSP/PSC kinetics (Fig. 2c, d) | `synaptic_response_rest` | `synaptic_response` |
| Connectivity and degree distributions (Fig. 3) and illustrations of population activity (Fig. 4d (background) and Fig. 5c,d) | `noise_driven_dynamics` | `illustrate_activity(*)` |
| Population rate transfer functions and responses to background noise (Fig. 4 and Supp. Fig. 1) | `noise_driven_dynamics` | `noisedriven_dynamics` |
| State transitions and statistics of active states (Fig. 5a and b, Fig. 6 and Supp. Fig. 3) | `state_transition` | `characterize_state_transition` |
| Stimulus parameters, memory and capacity (Fig. 7b and c, Fig. 8, 9 and Supp. Fig. 4) | `stimulus_driven` | `measure_capacity` |
| Capacity analysis on pre-stored data | `stimulus_driven` | `measure_capacity_offline` |

**Supplementary Table 3.** Summary of all the numerical experiments that can be run using the provided source code. Some are very memory and especially time-consuming (the capacity analysis, in particular). The item marked with (*) can be found in the scripts folder.

[5] https://rcfduarte.github.io/nmsat/

##### 4.3 Datasets

Currently, the data is only accessible through the OSF repository but, upon publication, it will be registered and made publicly available via a suitable platform. We provide all the datasets necessary to replicate the main experiments, as well as the original parameters used for each simulation (for scrutiny) and the main results. The datasets are organized as follows:

```
data
├── neuron_parameters
├── receptor_parameters
├── synapse_parameters
├── population_RTF
├── state_transitions
├── active_state
├── input_tuning
└── capacity
    ├── data_label_ParameterSpace.py
    └── data_label/
        ├── Activity/
        ├── Figures/
        ├── Inputs/
        ├── Parameters/
        ├── Results/
        └── Output/
```

The specific sub-folder structure varies depending on the dataset, but typically follows the organization described above for `capacity/`, used as an example. For the `data_label` of specific experiments consult Supplementary Table 4. The code package we provide as supplement contains all the analysis scripts (`read_data` folder).

| Description | Data label | Analysis script |
|---|---|---|
| Single neuron fI curves and distributions (Fig. 1) | single_neuron_dcinput | neuron_fI |
| Adaptation parameters for the different neuron classes $(a, b,$ data not shown) | single_neuron_excitability | singleneuron_excitability |
| Receptor conductance kinetics (data not shown) | *neuron_rec* | receptor_conductances |
| Distribution of PSP amplitudes and latencies (Fig. 2d) | *syn*_heterogeneous | plot_het_synapse |
| Population rate transfer functions (Fig. 4a, b, c and Supp. Fig. 1) | populationRTF_normal_*het* | populationRTF_spiking |
| Sub-threshold responses to background noise (Fig. 4d and Fig. 5a, b) | populationRTF_normal_*het* | populationRTF_subthreshold |
| State transitions and characterization of quiet and active states, single trial (Fig. 4e and 7a) | state_transitions_fixed_rate_*het* | plot_state_transitions |
| Spiking statistics in the active state (Fig. 9 and Supp. Fig. 3) | active_state_*het* | active_state_spiking |
| Input amplitude tuning (Supp. Fig. 2) | stimulus_tuning_*het* | stimulus_amplitude |
| Stimulus resolution (Supp. Fig. 4) | stimulus_resolution_*het* | stimulus_resolution |
| Memory capacity (Fig. 7b and c) | capacity_*het* | memory_capacity |
| Total processing capacity (Fig. 9) | capacity_*het* | total_capacity |
| Receptor composition and memory capacity (Fig. 8) | receptor_ratios_*het* | capacity_parameter_scan |

**Supplementary Table 4.** Summary of all datasets and corresponding scripts to analyse and plot the data. Note: For legacy reasons, the conditions are labelled as: HOM (homogeneous), HET1 (structural), HET2 (neuronal), HET3 (synaptic), HET or HETall (fully heterogeneous).

### References


Abe et al., 2011. Abe, Y., Namba, H., Kato, T., Iwakura, Y., and Nawa, H. (2011). Neuregulin-1 signals from the periphery regulate AMPA receptor sensitivity and expression in GABAergic interneurons in developing neocortex. *The Journal of neuroscience : the official journal of the Society for Neuroscience*, 31(15):5699–709.

Abel, 2003. Abel, H. J. (2003). Relationships Between Intracellular Calcium and Afterhyperpolarizations in Neocortical Pyramidal Neurons. *Journal of Neurophysiology*, 91(1):324–335.

Aerde et al., 2015. Aerde, K. I. V., Qi, G., and Feldmeyer, D. (2015). Cell type-specific effects of adenosine on cortical neurons. *Cerebral Cortex*, 25(3):772–787.

Akgul and Wollmuth, 2013. Akgul, G. and Wollmuth, L. P. (2013). Synapse-Associated Protein 97 Regulates the Membrane Properties of Fast-Spiking Parvalbumin Interneurons in the Visual Cortex. *Journal of Neuroscience*, 33(31):12739–12750.

Amatrudo et al., 2012. Amatrudo, J. M., Weaver, C. M., Crimins, J. L., Hof, P. R., Rosene, D. L., and Luebke, J. I. (2012). Influence of Highly Distinctive Structural Properties on the Excitability of Pyramidal Neurons in Monkey Visual and Prefrontal Cortices. *Journal of Neuroscience*, 32(40):13644–13660.

Andjelic et al., 2008. Andjelic, S., Gallopin, T., Cauli, B., Hill, E. L., Roux, L., Badr, S., Hu, E., Tamas, G., and Lambolez, B. (2008). Glutamatergic Nonpyramidal Neurons From Neocortical Layer VI and Their Comparison With Pyramidal and Spiny Stellate Neurons. *Journal of Neurophysiology*, 101(2):641–654.

Angulo et al., 1999. Angulo, M. C., Rossier, J., and Audinat, E. (1999). Postsynaptic glutamate receptors and integrative properties of fast-spiking interneurons in the rat neocortex. *Journal of neurophysiology*, 82(3):1295–1302.

Aramakis et al., 2000. Aramakis, V. B., Hsieh, C. Y., Leslie, F. M., and Metherate, R. (2000). A critical period for nicotine-induced disruption of synaptic development in rat auditory cortex. *The Journal of neuroscience : the official journal of the Society for Neuroscience*, 20(16):6106–6116.

Avermann et al., 2012. Avermann, M., Tomm, C., Mateo, C., Gerstner, W., and Petersen, C. C. H. (2012). Microcircuits of excitatory and inhibitory neurons in layer 2/3 of mouse barrel cortex. *Journal of Neurophysiology*, 107(11):3116–3134.

Barnes et al., 2015. Barnes, S. J., Cheetham, C. E., Liu, Y., Bennett, S. H., Albieri, G., Jorstad, A. A., Knott, G. W., and Finnerty, G. T. (2015). Delayed and Temporally Imprecise Neurotransmission in Reorganizing Cortical Microcircuits. *Journal of Neuroscience*, 35(24):9024–9037.

Beggs et al., 2000. Beggs, J. M., Moyer, J. R., McGann, J. P., and Brown, T. H. (2000). Prolonged synaptic integration in perirhinal cortical neurons. *Journal of neurophysiology*, 83(6):3294–3298.

Blatow et al., 2003. Blatow, M., Rozov, A., Katona, I., Hormuzdi, S. G., Meyer, A. H., Whittington, M. A., Caputi, A., and Monyer, H. (2003). A novel network of multipolar bursting interneurons generates theta frequency oscillations in neocortex. *Neuron*, 38(5):805–817.

Bos et al., 2015. Bos, H., Morrison, Abigail Peyser, Alexander Hahne, J., Helias, M., Kunkel, S., Ippen, T., Eppler, J. M., Schmidt, M., Seeholzer, A., Djurfeldt, M., Diaz, S., Morén, J., Deepu, R., Stocco, T., Deger, M., Michler, F., and Plesser, H. E. (2015). Nest 2.10.0. page DOI: 10.5281/ZENODO.44222.

Brill and Huguenard, 2010. Brill, J. and Huguenard, J. R. (2010). Enhanced infragranular and supragranular synaptic input onto layer 5 pyramidal neurons in a rat model of cortical dysplasia. *Cerebral Cortex*, 20(12):2926–2938.

Brunel and Wang, 2001. Brunel, N. and Wang, X. J. (2001). Effects of neuromodulation in a cortical network model of object working memory dominated by recurrent inhibition. *Journal of Computational Neuroscience*, 11(1):63–85.

Castro et al., 2002. Castro, P. A., Pleasure, S. J., and Baraban, S. C. (2002). Hippocampal heterotopia with molecular and electrophysiological properties of neocortical neurons. *Neuroscience*, 114(4):961–972.

Chadderton et al., 2009. Chadderton, P., Agapiou, J. P., McAlpine, D., and Margrie, T. W. (2009). The Synaptic Representation of Sound Source Location in Auditory Cortex. *Journal of Neuroscience*, 29(45):14127–14135.

Cheetham et al., 2007. Cheetham, C. E. J., Hammond, M. S. L., Edwards, C. E. J., and Finnerty, G. T. (2007). Sensory Experience Alters Cortical Connectivity and Synaptic Function Site Specifically. *Journal of Neuroscience*, 27(13):3456–3465.

Chen et al., 2015. Chen, I.-W., Helmchen, F., and Lutcke, H. (2015). Specific Early and Late Oddball-Evoked Responses in Excitatory and Inhibitory Neurons of Mouse Auditory Cortex. *Journal of Neuroscience*, 35(36):12560–12573.

Cho et al., 2010. Cho, K.-h., Jang, J. H., Jang, H.-j., Kim, M.-j., Hee, S., Fukuda, T., Tennigkeit, F., Singer, W., Rhie, D.-j., and Yoon, S. H. (2010). Subtype-Specific Dendritic Calcium Dynamics of Inhibitory Interneurons in the Rat Visual Cortex. *Journal of Neurophysiology*, 104(June 2010):840–853.

Connors et al., 1988. Connors, B. W., Malenka, R. C., and Silva, L. R. (1988). Two inhibitory postsynaptic potentials, and GABAA and GABAB receptor-mediated responses in neocortex of rat and cat. *The Journal of Physiology*, 406(1):443–468.

Cruikshank et al., 2012. Cruikshank, S. J., Ahmed, O. J., Stevens, T. R., Patrick, S. L., Gonzalez, A. N., Elmaleh, M., and Connors, B. W. (2012). Thalamic Control of Layer 1 Circuits in Prefrontal Cortex. *Journal of Neuroscience*, 32(49):17813–17823.

Cummings et al., 2009. Cummings, D. M., Andre, V. M., Uzgil, B. O., Gee, S. M., Fisher, Y. E., Cepeda, C., and Levine, M. S. (2009). Alterations in Cortical Excitation and Inhibition in Genetic Mouse Models of Huntington's Disease. *Journal of Neuroscience*, 29(33):10371–10386.

Dayan and Abbott, 2001. Dayan, P. and Abbott, L. F. (2001). *Theoretical Neuroscience: Computational and Mathematical Modeling of Neural Systems*, volume 15. The MIT Press, Cambridge, MA.

Destexhe et al., 1994. Destexhe, A., Mainen, Z. F., and Sejnowski, T. J. (1994). Synthesis of models for excitable membranes, synaptic transmission and neuromodulation using a common kinetic formalism. *Journal of Computational Neuroscience*, 1(3):195–230.

Destexhe et al., 1998. Destexhe, A., Mainen, Z. F., and Sejnowski, T. J. (1998). Kinetic models of synaptic transmission. In Koch, C. and Segev, I., editors, *Methods in Neuronal Modeling*, pages 1–25. MIT Press, Cambridge, MA, 2nd editio edition.

Destexhe and Sejnowski, 1995. Destexhe, A. and Sejnowski, T. J. (1995). G protein activation kinetics and spillover of gamma-aminobutyric acid may account for differences between inhibitory responses in the hippocampus and thalamus. *Proceedings of the National Academy of Sciences*, 92(21):9515–9519.

Dong et al., 2004. Dong, H., Shao, Z., Nerbonne, J. M., and Burkhalter, A. (2004). Differential depression of inhibitory synaptic responses in feedforward and feedback circuits between different areas of mouse visual cortex. *Journal of Comparative Neurology*, 475(3):361–373.

Dougherty et al., 2014. Dougherty, S. E., Bartley, A. F., Lucas, E. K., Hablitz, J. J., Dobrunz, L. E., and Cowell, R. M. (2014). Mice lacking the transcriptional coactivator PGC-1$\alpha$ exhibit alterations in inhibitory synaptic transmission in the motor cortex. *Neuroscience*, 271:137–148.

Duarte and Morrison, 2017. Duarte, R. and Morrison, A. (2017). Leveraging heterogeneity for neural computations with fading memory. *Open Science Framework*.

Duarte et al., 2017. Duarte, R., Zajzon, B., and Morrison, A. (2017). Neural Microcircuit Simulation And Analysis Toolkit. *Zenodo*.

Elstrott et al., 2014. Elstrott, J., Clancy, K. B., Jafri, H., Akimenko, I., and Feldman, D. E. (2014). Cellular mechanisms for response heterogeneity among L2/3 pyramidal cells in whisker somatosensory cortex. *Journal of Neurophysiology*, 112(2):233–248.

Erisir et al., 1999. Erisir, A., Lau, D., Rudy, B., and Leonard, C. S. (1999). Function of Specific K+ Channels in Sustained High-Frequency Firing of Fast-Spiking Neocortical Interneurons. *J Neurophysiol*, 82(5):2476–2489.

Feldmeyer et al., 2006. Feldmeyer, D., Lübke, J., and Sakmann, B. (2006). Efficacy and connectivity of intracolumnar pairs of layer 2/3 pyramidal cells in the barrel cortex of juvenile rats. *The Journal of Physiology*, 575(2):583–602.

Fino and Yuste, 2011. Fino, E. and Yuste, R. (2011). Dense inhibitory connectivity in neocortex. *Neuron*, 69(6):1188–1203.

Gao et al., 2003. Gao, W. J., Wang, Y., and Goldman-Rakic, P. S. (2003). Dopamine modulation of perisomatic and peridendritic inhibition in prefrontal cortex. *J Neurosci*, 23(5):1622–1630.

Gerstner and Kistler, 2002. Gerstner, W. and Kistler, W. M. (2002). *Spiking Neuron Models*. Cambridge University Press.

Gerstner et al., 2014. Gerstner, W., Kistler, W. M., Naud, R., and Paninski, L. (2014). *Neuronal Dynamics - from single neurons to networks and models of cognition*. Cambridge University Press.

Gonzalez-Burgos, 2004. Gonzalez-Burgos, G. (2004). Functional Properties of Fast Spiking Interneurons and Their Synaptic Connections With Pyramidal Cells in Primate Dorsolateral Prefrontal Cortex. *Journal of Neurophysiology*, 93(2):942–953.

González-Burgos et al., 2004. González-Burgos, G., Krimer, L. S., Urban, N. N., Barrionuevo, G., and Lewis, D. A. (2004). Synaptic Efficacy during Repetitive Activation of Excitatory Inputs in Primate Dorsolateral Prefrontal Cortex. *Cerebral Cortex*, 14(5):530–542.

Gonzalez-Burgos et al., 2015. Gonzalez-Burgos, G., Miyamae, T., Pafundo, D. E., Yoshino, H., Rotaru, D. C., Hoftman, G., Datta, D., Zhang, Y., Hammond, M., Sampson, A. R., Fish, K. N., Ermentrout, G. B., and Lewis, D. A. (2015). Functional maturation of GABA synapses during postnatal development of the monkey dorsolateral prefrontal cortex. *Cerebral Cortex*, 25(11):4076–4093.

Gorelova, 2002. Gorelova, N. (2002). Mechanisms of Dopamine Activation of Fast-Spiking Interneurons That Exert Inhibition in Rat Prefrontal Cortex. *Journal of Neurophysiology*, 88(6):3150–3166.

Guan et al., 2007. Guan, D., Lee, J. C. F., Higgs, M. H., Spain, W. J., and Foehring, R. C. (2007). Functional Roles of Kv1 Channels in Neocortical Pyramidal Neurons. *Journal of Neurophysiology*, 97(3):1931–1940.

Gulledge et al., 2009. Gulledge, A. T., Bucci, D. J., Zhang, S. S., Matsui, M., and Yeh, H. H. (2009). M1 Receptors Mediate Cholinergic Modulation of Excitability in Neocortical Pyramidal Neurons. *Journal of Neuroscience*, 29(31):9888–9902.

Hadjilambreva, 2004. Hadjilambreva, G. (2004). Neuromodulation by a Cytokine: Interferon- Differentially Augments Neocortical Neuronal Activity and Excitability. *Journal of Neurophysiology*, 93(2):843–852.

Hagen et al., 2016. Hagen, E., Dahmen, D., Stavrinou, M. L., Lindén, H., Tetzlaff, T., Van Albada, S. J., Grün, S., Diesmann, M., and Einevoll, G. T. (2016). Hybrid scheme for modeling local field potentials from point-neuron networks. *Cerebral Cortex*, 26(12):4461–4496.

Harrison et al., 2015. Harrison, P. M., Badel, L., Wall, M. J., and Richardson, M. J. (2015). Experimentally Verified Parameter Sets for Modelling Heterogeneous Neocortical Pyramidal-Cell Populations. *PLoS Computational Biology*, 11(8):e1004165.

Helmstaedter et al., 2009. Helmstaedter, M., Sakmann, B., and Feldmeyer, D. (2009). L2/3 Interneuron groups defined by multiparameter analysis of axonal projection, dendritic geometry, and electrical excitability. *Cerebral Cortex*, 19(4):951–962.

Helmstaedter et al., 2008. Helmstaedter, M., Staiger, J. F., Sakmann, B., and Feldmeyer, D. (2008). Efficient Recruitment of Layer 2/3 Interneurons by Layer 4 Input in Single Columns of Rat Somatosensory Cortex. *Journal of Neuroscience*, 28(33):8273–8284.

Hestrin, 1993. Hestrin, S. (1993). Different glutamate receptor channels mediate fast excitatory synaptic currents in inhibitory and excitatory cortical neurons. *Neuron*, 11(6):1083–1091.

Hestrin et al., 1990. Hestrin, S., Sah, P., and Nicoll, R. A. (1990). Mechanisms generating the time course of dual component excitatory synaptic currents recorded in hippocampal slices. *Neuron*, 5(3):247–253.

Hill et al., 2000. Hill, E., Kalloniatis, M., and Tan, S. S. (2000). Glutamate, GABA and precursor amino acids in adult mouse neocortex: cellular diversity revealed by quantitative immunocytochemistry. *Cerebral cortex (New York, N.Y. : 1991)*, 10(11):1132–42.

Hirai et al., 2012. Hirai, Y., Morishima, M., Karube, F., and Kawaguchi, Y. (2012). Specialized Cortical Subnetworks Differentially Connect Frontal Cortex to Parahippocampal Areas. *Journal of Neuroscience*, 32(5):1898–1913.

Hoffmann et al., 2015. Hoffmann, J. H., Meyer, H. S., Schmitt, A. C., Straehle, J., Weitbrecht, T., Sakmann, B., and Helmstaedter, M. (2015). Synaptic conductance estimates of the connection between local inhibitor interneurons and pyramidal neurons in layer 2/3 of a cortical column. *Cerebral Cortex*, 25(11):4415–4429.

Huggenberger et al., 2009. Huggenberger, S., Vater, M., and Deisz, R. A. (2009). Interlaminar differences of intrinsic properties of pyramidal neurons in the auditory cortex of mice. *Cerebral Cortex*, 19(5):1008–1018.

Imbrosci et al., 2014. Imbrosci, B., Neitz, A., and Mittmann, T. (2014). Focal cortical lesions induce bidirectional changes in the excitability of fast spiking and non fast spiking cortical interneurons. *PLoS ONE*, 9(10):e111105.

Izhikevich and Edelman, 2008. Izhikevich, E. M. and Edelman, G. M. (2008). Large-scale model of mammalian thalamocortical systems. *Proceedings of the National Academy of Sciences*, 105(9):3593–3598.

Izhikevich et al., 2004. Izhikevich, E. M., Gally, J. A., and Edelman, G. M. (2004). Spike-timing dynamics of neuronal groups. *Cerebral Cortex*, 14(8):933–944.

Jahr and Stevens, 1990. Jahr, C. E. and Stevens, C. F. (1990). Voltage dependence of NMDA-activated macroscopic conductances predicted by single-channel kinetics. *The Journal of neuroscience*, 10(9):3178–3182.

Karayannis et al., 2007. Karayannis, T., Huerta-Ocampo, I., and Capogna, M. (2007). GABAergic and pyramidal neurons of deep cortical layers directly receive and differently integrate callosal input. *Cerebral Cortex*, 17(5):1213–1226.

Kawaguchi and Kubota, 1998. Kawaguchi, Y. and Kubota, Y. (1998). Neurochemical features and synaptic connections of large physiologically-identified GABAergic cells in the rat frontal cortex. *Neuroscience*, 85(3):677–701.

Kobayashi et al., 2008. Kobayashi, M., Sasabe, T., Shiohama, Y., and Koshikawa, N. (2008). Activation of ??1-adrenoceptors increases firing frequency through protein kinase C in pyramidal neurons of rat visual cortex. *Neuroscience Letters*, 430(2):175–180.

Köhr et al., 1993. Köhr, G., De Koninck, Y., and Mody, I. (1993). Properties of NMDA receptor channels in neurons acutely isolated from epileptic (kindled) rats. *The Journal of neuroscience : the official journal of the Society for Neuroscience*, 13(8):3612–3627.

Koulakov et al., 2009. Koulakov, A. A., Hromadka, T., and Zador, A. M. (2009). Correlated Connectivity and the Distribution of Firing Rates in the Neocortex. *Journal of Neuroscience*, 29(12):3685–3694.

Krimer, 2005. Krimer, L. S. (2005). Cluster Analysis-Based Physiological Classification and Morphological Properties of Inhibitory Neurons in Layers 2-3 of Monkey Dorsolateral Prefrontal Cortex. *Journal of Neurophysiology*, 94(5):3009–3022.

Kröner et al., 2007. Kröner, S., Krimer, L. S., Lewis, D. A., and Barrionuevo, G. (2007). Dopamine increases inhibition in the monkey dorsolateral prefrontal cortex through cell type-specific modulation of interneurons. *Cerebral Cortex*, 17(5):1020–1032.

Kubota et al., 2015. Kubota, Y., Kondo, S., Nomura, M., Hatada, S., Yamaguchi, N., Mohamed, A. A., Karube, F., Lübke, J., and Kawaguchi, Y. (2015). Functional effects of distinct innervation styles of pyramidal cells by fast spiking cortical interneurons. *eLife*, 4(July 2015):1–27.

Lacroix et al., 2015. Lacroix, A., Toussay, X., Anenberg, E., Lecrux, C., Ferreiros, N., Karagiannis, A., Plaisier, F., Chausson, P., Jarlier, F., Burgess, S. A., Hillman, E. M. C., Tegeder, I., Murphy, T. H., Hamel, E., and Cauli, B. (2015). COX-2-Derived Prostaglandin E2 Produced by Pyramidal Neurons Contributes to Neurovascular Coupling in the Rodent Cerebral Cortex. *Journal of Neuroscience*, 35(34):11791–11810.

Lambe et al., 2000. Lambe, E. K., Goldman-Rakic, P. S., and Aghajanian, G. K. (2000). Serotonin induces EPSCs preferentially in layer V pyramidal neurons of the frontal cortex in the rat. *Cerebral cortex (New York, N.Y. : 1991)*, 10(10):974–980.

Lee et al., 2007. Lee, C. M., Chang, W. C., Chang, K. B., and Shyu, B. C. (2007). Synaptic organization and input-specific short-term plasticity in anterior cingulate cortical neurons with intact thalamic inputs. *European Journal of Neuroscience*, 25(9):2847–2861.

Lee et al., 2005. Lee, J. C. F., Callaway, J. C., and Foehring, R. C. (2005). Effects of Temperature on Calcium Transients and Ca 2+ -Dependent Afterhyperpolarizaions in Neocortical Pyramidal Neurons. *Journal of Neurophysiology*, 93(4):2012–2020.

Lefort et al., 2009. Lefort, S., Tomm, C., Floyd Sarria, J. C., and Petersen, C. C. H. (2009). The Excitatory Neuronal Network of the C2 Barrel Column in Mouse Primary Somatosensory Cortex. *Neuron*, 61(2):301–316.

Lemtiri-Chlieh and Levine, 2007. Lemtiri-Chlieh, F. and Levine, E. S. (2007). Lack of Depolarization-Induced Suppression of Inhibition (DSI) in Layer 2/3 Interneurons That Receive Cannabinoid-Sensitive Inhibitory Inputs. *Journal of Neurophysiology*, 98(5):2517–2524.

Levy and Reyes, 2012. Levy, R. B. and Reyes, A. D. (2012). Spatial Profile of Excitatory and Inhibitory Synaptic Connectivity in Mouse Primary Auditory Cortex. *Journal of Neuroscience*, 32(16):5609–5619.

Lu et al., 2007. Lu, J.-t., Li, C.-y., Zhao, J.-P., Poo, M.-m., and Zhang, X.-h. (2007). Spike-Timing-Dependent Plasticity of Neocortical Excitatory Synapses on Inhibitory Interneurons Depends on Target Cell Type. *Journal of Neuroscience*, 27(36):9711–9720.

Luebke et al., 2015. Luebke, J. I., Medalla, M., Amatrudo, J. M., Weaver, C. M., Crimins, J. L., Hunt, B., Hof, P. R., and Peters, A. (2015). Age-related changes to layer 3 pyramidal cells in the rhesus monkey visual cortex. *Cerebral Cortex*, 25(6):1454–1468.

Luhmann et al., 1998. Luhmann, H. J., Karpuk, N., Qü, M., Zilles, K., Brill, J., Huguenard, J. R., Hutnick, L. K., Golshani, P., Namihira, M., Xue, Z., Matynia, A., Yang, W., Silva, A. J., Schweizer, F. E., and Fan, G. (1998). Characterization of Neuronal Migration Disorders in Neocortical Structures . II . Intracellular In Vitro Recordings Characterization of Neuronal Migration Disorders in Neocortical Structures . II . Intracellular In Vitro Recordings. *Journal of neurophysiology*, 80(1):92–102.

Mason et al., 1991. Mason, a., Nicoll, A., and Stratford, K. (1991). Synaptic transmission between individual pyramidal neurons of the rat visual cortex in vitro. *The Journal of Neuroscience*, 11(January):72–84.

McCormick et al., 1993. McCormick, D. A., Wang, Z., and Huguenard, J. (1993). Neurotransmitter control of neocortical neuronal activity and excitability. *Cerebral Cortex*, 3(5):387–398.

Miyoshi et al., 2007. Miyoshi, G., Butt, S. J. B., Takebayashi, H., and Fishell, G. (2007). Physiologically Distinct Temporal Cohorts of Cortical Interneurons Arise from Telencephalic Olig2-Expressing Precursors. *Journal of Neuroscience*, 27(29):7786–7798.

Monier et al., 2008. Monier, C., Fournier, J., and Frégnac, Y. (2008). In vitro and in vivo measures of evoked excitatory and inhibitory conductance dynamics in sensory cortices. *Journal of Neuroscience Methods*, 169(2):323–365.

Moreau and Kullmann, 2013. Moreau, A. W. and Kullmann, D. M. (2013). NMDA receptor-dependent function and plasticity in inhibitory circuits. *Neuropharmacology*, 74:23–31.

Mowery et al., 2015. Mowery, T. M., Kotak, V. C., and Sanes, D. H. (2015). Transient Hearing Loss Within a Critical Period Causes Persistent Changes to Cellular Properties in Adult Auditory Cortex. *Cerebral Cortex*, 25(8):2083–2094.

Myme, 2003. Myme, C. I. O. (2003). The NMDA-to-AMPA Ratio at Synapses Onto Layer 2/3 Pyramidal Neurons Is Conserved Across Prefrontal and Visual Cortices. *Journal of Neurophysiology*, 90(2):771–779.

Neske et al., 2015. Neske, G. T., Patrick, S. L., and Connors, B. W. (2015). Contributions of Diverse Excitatory and Inhibitory Neurons to Recurrent Network Activity in Cerebral Cortex. *Journal of Neuroscience*, 35(3):1089–1105.

Nissen et al., 2010. Nissen, W., Szabo, A., Somogyi, J., Somogyi, P., and Lamsa, K. P. (2010). Cell Type-Specific Long-Term Plasticity at Glutamatergic Synapses onto Hippocampal Interneurons Expressing either Parvalbumin or CB1 Cannabinoid Receptor. *Journal of Neuroscience*, 30(4):1337–1347.

Nordlie et al., 2009. Nordlie, E., Gewaltig, M. O., and Plesser, H. E. (2009). Towards reproducible descriptions of neuronal network models. *PLoS Computational Biology*, 5(8):e1000456.

Nowak, 2002. Nowak, L. G. (2002). Electrophysiological Classes of Cat Primary Visual Cortical Neurons In Vivo as Revealed by Quantitative Analyses. *Journal of Neurophysiology*, 89(3):1541–1566.

Oswald and Reyes, 2008. Oswald, A.-M. M. and Reyes, A. D. (2008). Maturation of Intrinsic and Synaptic Properties of Layer 2/3 Pyramidal Neurons in Mouse Auditory Cortex. *Journal of Neurophysiology*, 99(6):2998–3008.

Oswald and Reyes, 2011. Oswald, A. M. M. and Reyes, A. D. (2011). Development of inhibitory timescales in auditory cortex. *Cerebral Cortex*, 21(6):1351–1361.

Paoletti et al., 2013. Paoletti, P., Bellone, C., and Zhou, Q. (2013). NMDA receptor subunit diversity: impact on receptor properties, synaptic plasticity and disease. *Nature Reviews Neuroscience*, 14(6):383–400.

Perin et al., 2011. Perin, R., Berger, T. K., and Markram, H. (2011). A synaptic organizing principle for cortical neuronal groups. *Proceedings of the National Academy of Sciences*, 108(13):5419–5424.

Pi et al., 2013. Pi, H.-J., Hangya, B., Kvitsiani, D., Sanders, J. I., Huang, Z. J., and Kepecs, A. (2013). Cortical interneurons that specialize in disinhibitory control. *Nature*, 503(7477):521–524.

Pillai et al., 2014. Pillai, A. G., Henckens, M. J. A. G., Fernández, G., and Joëls, M. (2014). Delayed effects of corticosterone on slow after-hyperpolarization potentials in mouse hippocampal versus prefrontal cortical pyramidal neurons. *PLoS ONE*, 9(6):e99208.

Pineda et al., 1998. Pineda, J. C., Waters, R. S., and Foehring, R. C. (1998). Specificity in the interaction of HVA Ca2+ channel types with Ca2+-dependent AHPs and firing behavior in neocortical pyramidal neurons. *J Neurophysiol*, 79(5):2522–2534.

Potjans and Diesmann, 2014. Potjans, T. C. and Diesmann, M. (2014). The cell-type specific cortical microcircuit: Relating structure and activity in a full-scale spiking network model. *Cerebral Cortex*, 24(3):785–806.

Povysheva et al., 2006. Povysheva, N. V., Gonzalez-Burgos, G., Zaitsev, A. V., Kr??ner, S., Barrionuevo, G., Lewis, D. A., and Krimer, L. S. (2006). Properties of excitatory synaptic responses in fast-spiking interneurons and pyramidal cells from monkey and rat prefrontal cortex. *Cerebral Cortex*, 16(4):541–552.

Povysheva et al., 2008. Povysheva, N. V., Zaitsev, A. V., Rotaru, D. C., Gonzalez-Burgos, G., Lewis, D. A., and Krimer, L. S. (2008). Parvalbumin-positive basket interneurons in monkey and rat prefrontal cortex. *Journal of neurophysiology*, 100(4):2348–60.

Ren et al., 2014. Ren, M., Cao, V., Ye, Y., Manji, H. K., and Wang, K. H. (2014). Arc Regulates Experience-Dependent Persistent Firing Patterns in Frontal Cortex. *Journal of Neuroscience*, 34(19):6583–6595.

Rose, 2005. Rose, H. J. (2005). Auditory Thalamocortical Transmission Is Reliable and Temporally Precise. *Journal of Neurophysiology*, 94(3):2019–2030.

Sah et al., 1990. Sah, P., Hestrin, S., and Nicoll, R. a. (1990). Properties of excitatory postsynaptic currents recorded in vitro from rat hippocampal interneurones. *The Journal of physiology*, 430:605–616.

Salling et al., 2014. Salling, M. C., Harrison, N. L., Abernathy, K., Chandler, L., Woodward, J., Amatrudo, J., Weaver, C., Crimins, J., Hof, P., Rosene, D., Luebke, J., Badanich, K., Mulholland, P., Beckley, J., Trantham-Davidson, H., Woodward, J., Bai, D., Zhu, G., Pennefather, P., Jackson, M., MacDonald, J., Orser, B., Belelli, D., Harrison, N., Maguire, J., Macdonald, R., Walker, M., Cope, D., Bosman, L., Rosahl, T., Brussaard, A., Brickley, S., Cull-Candy, S., Farrant, M., Chattipakorn, S., McMahon, L., Chattipakorn, S., McMahon, L., Chudasama, Y., Robbins, T., Cope, D., Hughes, S., Crunelli, V., Dahchour, A., Hoffman, A., Deitrich, R., de Witte, P., Day, M., Carr, D., Ulrich, S., Ilijic, E., Tkatch, T., Surmeier, D., DeFelipe, J., Lopez-Cruz, P., Benavides-Piccione, R., Bielza,

C., Larranaga, P., Anderson, S., Burkhalter, A., Cauli, B., Fairen, A., Feldmeyer, D., Fishell, G., Fitzpatrick, D., Freund, T., Gonzalez-Burgos, G., Hestrin, S., Hill, S., Hof, P., Huang, J., Jones, E., Kawaguchi, Y., Kisvarday, Z., Kubota, Y., Lewis, D., Marin, O., Markram, H., McBain, C., Meyer, H., Monyer, H., Nelson, S., Rockland, K., Rossier, J., Rubenstein, J., Rudy, B., Scanziani, M., Shepherd, G., Sherwood, C., Staiger, J., Tamas, G., Thomson, A., Wang, Y., Yuste, R., Ascoli, G., Drasbek, K., Jensen, K., Elliott, R., Baker, S., Rogers, R., O'Leary, D., Paykel, E., Frith, C., Dolan, R., Sahakian, B., Farrant, M., Nusser, Z., Fino, E., Packer, A., Yuste, R., Flint, A., Liu, X., Kriegstein, A., Franklin, K., Paxinos, G., Franklin, K., Paxinos, G., Fritschy, J., Mohler, H., Goldstein, R., Volkow, N., Harsing, L., Matyus, P., Hedrick, T., Waters, J., Hoestgaard-Jensen, K., Dalby, N., Wolinsky, T., Murphey, C., Jones, K., Rottlander, M., Frederiksen, K., Watson, W., Jensen, K., Ebert, B., Horio, M., Kohno, M., Fujita, Y., Ishima, T., Inoue, R., Mori, H., Hashimoto, K., Jia, F., Chandra, D., Homanics, G., Harrison, N., Jia, F., Goldstein, P., Harrison, N., Jia, F., Pignataro, L., Schofield, C., Yue, M., Harrison, N., Goldstein, P., Jonsson, S., Kerekes, N., Hyytia, P., Ericson, M., Soderpalm, B., Kawaguchi, Y., Kirsch, J., Krettek, J., Price, J., Kunz, P., Burette, A., Weinberg, R., Philpot, B., Lu, Y., Ye, J., Malosio, M., Marqueze-Pouey, B., Kuhse, J., Betz, H., Mori, M., Gahwiler, B., Gerber, U., Murakami, T., Furuse, M., Nusser, Z., Mody, I., Olah, S., Fule, M., Komlosi, G., Varga, C., Baldi, R., Barzo, P., Tamas, G., Ongur, D., Price, J., Peng, Z., Hauer, B., Mihalek, R., Homanics, G., Sieghart, W., Olsen, R., Houser, C., Petersen, C., Crochet, S., Pirker, S., Schwarzer, C., Wieselthaler, A., Sieghart, W., Sperk, G., Pribilla, I., Takagi, T., Langosch, D., Bormann, J., Betz, H., Rao, S., Williams, G., Goldman-Rakic, P., Rudy, B., Fishell, G., Lee, S., Hjerling-Leffler, J., Salin, P., Tseng, G., Hoffman, S., Parada, I., Prince, D., Saxena, N., Macdonald, R., Seamans, J., Lapish, C., Durstewitz, D., Semyanov, A., Walker, M., Kullmann, D., Thuault, S., Malleret, G., Constantinople, C., Nicholls, R., Chen, I., Zhu, J., Panteleyev, A., Vronskaya, S., Nolan, M., Bruno, R., Siegelbaum, S., Kandel, E., Uylings, H., Groenewegen, H., Kolb, B., Vandenberg, R., Aubrey, K., Vardya, I., Drasbek, K., Dosa, Z., Jensen, K., Vicini, S., Ferguson, C., Prybylowski, K., Kralic, J., Morrow, A., Homanics, G., Weinberger, D., Berman, K., Zec, R., Weitlauf, C., Woodward, J., Yamada, J., Furukawa, T., Ueno, S., Yamamoto, S., Fukuda, A., Yoon, J., Minzenberg, M., Ursu, S., Walter, B. R., Wendelken, C., Ragland, J., Carter, C., Zhang, L., Gong, N., Fei, D., Xu, L., Xu, T., Zhang, X., Sun, G., Liu, L., Yu, F., and Xu, T. (2014). Strychnine-sensitive glycine receptors on pyramidal neurons in layers II/III of the mouse prefrontal cortex are tonically activated. *Journal of neurophysiology*, 112(5):1169–78.

Schiff and Reyes, 2012. Schiff, M. L. and Reyes, A. D. (2012). Characterization of thalamocortical responses of regular-spiking and fast-spiking neurons of the mouse auditory cortex in vitro and in silico. *Journal of Neurophysiology*, 107(5):1476–1488.

Sessler et al., 1998. Sessler, F. M., Hsu, F. C., Felder, T. N., Zhai, J., Lin, R. C., Wieland, S. J., and Kosobud, a. E. (1998). Effects of ethanol on rat somatosensory cortical neurons. *Brain research*, 804(2):266–74.

Shruti et al., 2008. Shruti, S., Clem, R. L., and Barth, A. L. (2008). A seizure-induced gain-of-function in BK channels is associated with elevated firing activity in neocortical pyramidal neurons. *Neurobiology of Disease*, 30(3):323–330.

Sippy and Yuste, 2013. Sippy, T. and Yuste, R. (2013). Decorrelating Action of Inhibition in Neocortical Networks. *Journal of Neuroscience*, 33(23):9813–9830.

Song et al., 2005. Song, S., Sj??str??m, P. J., Reigl, M., Nelson, S., and Chklovskii, D. B. (2005). Highly nonrandom features of synaptic connectivity in local cortical circuits. *PLoS Biology*, 3(3):0507–0519.

Southan et al., 2016. Southan, C., Sharman, J. L., Benson, H. E., Faccenda, E., Pawson, A. J., Alexander, S. P., Buneman, O. P., Davenport, A. P., McGrath, J. C., Peters, J. A., Spedding, M., Catterall, W. A., Fabbro, D., and Davies, J. A. (2016). The IUPHAR/BPS Guide to PHARMACOLOGY in 2016: Towards curated quantitative interactions between 1300 protein targets and 6000 ligands. *Nucleic Acids Research*, 44(D1):D1054–D1068.

Sutor et al., 2000. Sutor, B., Schmolke, C., Teubner, B., Schirmer, C., and Willecke, K. (2000). Myelination Defects and Neuronal Hyperexcitability in the Neocortex of Connexin 32-deficient Mice. *Cerebral Cortex*, 10(7):684–697.

Szabadics, 2006. Szabadics, J. (2006). Excitatory Effect of GABAergic Axo-Axonic Cells in Cortical Microcircuits. *Science*, 311(5758):233–235.

Takei et al., 2010. Takei, H., Fujita, S., Shirakawa, T., Koshikawa, N., and Kobayashi, M. (2010). Insulin facilitates repetitive spike firing in rat insular cortex via phosphoinositide 3-kinase but not mitogen activated protein kinase cascade. *Neuroscience*, 170(4):1199–1208.

Takesian et al., 2012. Takesian, A. E., Kotak, V. C., and Sanes, D. H. (2012). Age-dependent effect of hearing loss on cortical inhibitory synapse function. *Journal of Neurophysiology*, 107(3):937–947.

Tanaka et al., 2011. Tanaka, Y. R., Tanaka, Y. H., Konno, M., Fujiyama, F., Sonomura, T., Okamoto-Furuta, K., Kameda, H., Hioki, H., Furuta, T., Nakamura, K. C., and Kaneko, T. (2011). Local Connections of Excitatory Neurons to Corticothalamic Neurons in the Rat Barrel Cortex. *Journal of Neuroscience*, 31(50):18223–18236.

Tateno, 2004. Tateno, T. (2004). Threshold Firing Frequency-Current Relationships of Neurons in Rat Somatosensory Cortex: Type 1 and Type 2 Dynamics. *Journal of Neurophysiology*, 92(4):2283–2294.

Tateno and Robinson, 2008. Tateno, T. and Robinson, H. (2008). Integration of Broadband Conductance Input in Rat Somatosensory Cortical Inhibitory Interneurons: An Inhibition-Controlled Switch Between Intrinsic and Input-Driven Spiking in Fast-Spiking Cells. *Journal of Neurophysiology*, 101(2):1056–1072.

Telfeian and Connors, 2003. Telfeian, A. E. and Connors, B. W. (2003). Widely integrative properties of layer 5 pyramidal cells support a role for processing of extralaminar synaptic inputs in rat neocortex. *Neuroscience Letters*, 343(2):121–124.

Thomson, 2002. Thomson, A. M. (2002). Synaptic Connections and Small Circuits Involving Excitatory and Inhibitory Neurons in Layers 2-5 of Adult Rat and Cat Neocortex: Triple Intracellular Recordings and Biocytin Labelling In Vitro. *Cerebral Cortex*, 12(9):936–953.

Tomm, 2012. Tomm, C. (2012). *Analysing Neuronal Network Architectures: From Weight Distributions to Structure and Back*. PhD thesis, ÉCOLE POLYTECHNIQUE FÉDÉRALE DE LAUSANNE.

Tomm et al., 2014. Tomm, C., Avermann, M., Petersen, C., Gerstner, W., and Vogels, T. P. (2014). Connection-type-specific biases make uniform random network models consistent with cortical recordings. *Journal of Neurophysiology*, 112(8):1801–1814.

Tripathy et al., 2015. Tripathy, S. J., Burton, S. D., Geramita, M., Gerkin, R. C., and Urban, N. N. (2015). Brain-wide analysis of electrophysiological diversity yields novel categorization of mammalian neuron types. *Journal of Neurophysiology*, 113(10):3474–3489.

Tripathy et al., 2014. Tripathy, S. J., Savitskaya, J., Burton, S. D., Urban, N. N., and Gerkin, R. C. (2014). NeuroElectro: a window to the world's neuron electrophysiology data. *Frontiers in Neuroinformatics*, 8(April):40.

Tyler et al., 2015. Tyler, W. A., Medalla, M., Guillamon-Vivancos, T., Luebke, J. I., and Haydar, T. F. (2015). Neural Precursor Lineages Specify Distinct Neocortical Pyramidal Neuron Types. *Journal of Neuroscience*, 35(15):6142–6152.

Uzun et al., 2010. Uzun, S., Kozumplik, O., Jakovljević, M., and Sedić, B. (2010). Side effects of treatment with benzodiazepines. *Psychiatria Danubina*, 22(1):90–93.

Valdés-Sánchez et al., 2007. Valdés-Sánchez, L., Escámez, T., Echevarria, D., Ballesta, J. J., Tabarés-Seisdedos, R., Reiner, O., Martinez, S., and Geijo-Barrientos, E. (2007). Postnatal alterations of the inhibitory synaptic responses recorded from cortical pyramidal neurons in the Lis1/sLis1 mutant mouse. *Molecular and Cellular Neuroscience*, 35(2):220–229.

Van Aerde and Feldmeyer, 2015. Van Aerde, K. I. and Feldmeyer, D. (2015). Morphological and physiological characterization of pyramidal neuron subtypes in rat medial prefrontal cortex. *Cerebral Cortex*, 25(3):788–805.

van der Velden et al., 2012. van der Velden, L., van Hooft, J. A., and Chameau, P. (2012). Altered dendritic complexity affects firing properties of cortical layer 2/3 pyramidal neurons in mice lacking the 5-HT3A receptor. *Journal of Neurophysiology*, 108(5):1521–1528.

Vogl et al., 2015. Vogl, R., Rudolph, D., Angenent, H., Thoring, A., Schild, C., Stieglitz, S., and Meske, C. (2015). sciebo - the Campuscloud: a Sync and Share Cloud Storage Service for the Academic and Research Community in North Rhine-Westphalia. *PIK - Praxis der Informationsverarbeitung und Kommunikation*, 38(3-4):105–112.

Wang and Gao, 2010. Wang, H.-X. and Gao, W.-J. (2010). Development of calcium-permeable AMPA receptors and their correlation with NMDA receptors in fast-spiking interneurons of rat prefrontal cortex. *The Journal of Physiology*, 588(15):2823–2838.

Waters and Helmchen, 2006. Waters, J. and Helmchen, F. (2006). Background Synaptic Activity Is Sparse in Neocortex. *Journal of Neuroscience*, 26(32):8267–8277.

Watt et al., 2000. Watt, A. J., Van Rossum, M. C., MacLeod, K. M., Nelson, S. B., and Turrigiano, G. G. (2000). Activity coregulates quantal AMPA and NMDA currents at neocortical synapses. *Neuron*, 26(3):659–670.

Wayman et al., 2015. Wayman, W. N., Chen, L., Napier, T. C., and Hu, X.-T. (2015). Cocaine self-administration enhances excitatory responses of pyramidal neurons in the rat medial prefrontal cortex to human immunodeficiency virus-1 Tat. *European Journal of Neuroscience*, 41(9):1195–1206.

Yang et al., 2013a. Yang, J.-M., Zhang, J., Chen, X.-J., Geng, H.-Y., Ye, M., Spitzer, N. C., Luo, J.-H., Duan, S.-M., and Li, X.-M. (2013a). Development of GABA Circuitry of Fast-Spiking Basket Interneurons in the Medial Prefrontal Cortex of erbb4-Mutant Mice. *Journal of Neuroscience*, 33(50):19724–19733.

Yang and Benardo, 1997. Yang, L. and Benardo, L. S. (1997). Epileptogenesis following neocortical trauma from two sources of disinhibition. *Journal of neurophysiology*, 78(5):2804–2810.

Yang et al., 2013b. Yang, W., Carrasquillo, Y., Hooks, B. M., Nerbonne, J. M., and Burkhalter, A. (2013b). Distinct Balance of Excitation and Inhibition in an Interareal Feedforward and Feedback Circuit of Mouse Visual Cortex. *Journal of Neuroscience*, 33(44):17373–17384.

Yoshimura and Callaway, 2005. Yoshimura, Y. and Callaway, E. M. (2005). Fine-scale specificity of cortical networks depends on inhibitory cell type and connectivity. *Nature Neuroscience*, 8(11):1552–1559.

Yoshimura et al., 2005. Yoshimura, Y., Dantzker, J. L. M., and Callaway, E. M. (2005). Excitatory cortical neurons form fine-scale functional networks. *Nature*, 433(7028):868–873.

Zaitsev et al., 2012. Zaitsev, A. V., Povysheva, N. V., Gonzalez-Burgos, G., and Lewis, D. A. (2012). Electrophysiological classes of layer 2/3 pyramidal cells in monkey prefrontal cortex. *Journal of Neurophysiology*, 108(2):595–609.

Zaitsev et al., 2009. Zaitsev, A. V., Povysheva, N. V., Gonzalez-Burgos, G., Rotaru, D., Fish, K. N., Krimer, L. S., and Lewis, D. A. (2009). Interneuron diversity in layers 2-3 of monkey prefrontal cortex. *Cerebral Cortex*, 19(7):1597–1615.

Zhong and Yan, 2016. Zhong, P. and Yan, Z. (2016). Distinct Physiological Effects of Dopamine D4 Receptors on Prefrontal Cortical Pyramidal Neurons and Fast-Spiking Interneurons. *Cerebral Cortex*, 26(1):180–191.

Zhou and Roper, 2011. Zhou, F. W. and Roper, S. N. (2011). Altered firing rates and patterns in interneurons in experimental cortical Dysplasia. *Cerebral Cortex*, 21(7):1645–1658.
